# Molecular control of temporal integration matches decision-making to motivational state

**DOI:** 10.1101/2024.03.01.582988

**Authors:** Aditya K. Gautham, Lauren E. Miner, Marco N. Franco, Stephen C. Thornquist, Michael A. Crickmore

**Author notes:** Laboratory of Integrative Brain Function, The Rockefeller University (Stephen C. Thornquist).

## Abstract

Motivations bias our responses to stimuli, producing behavioral outcomes that match our needs and goals. We describe a mechanism behind this phenomenon: adjusting the time over which stimulus-derived information is permitted to accumulate toward a decision. As a Drosophila copulation progresses, the male becomes less likely to continue mating through challenges. We show that a set of Copulation Decision Neurons (CDNs) flexibly integrates information about competing drives to mediate this decision. Early in mating, dopamine signaling restricts CDN integration time by potentiating CaMKII activation in response to stimulatory inputs, imposing a high threshold for changing behaviors. Later into mating, the timescale over which the CDNs integrate termination-promoting information expands, increasing the likelihood of switching behaviors. We suggest scalable windows of temporal integration at dedicated circuit nodes as a key but underappreciated variable in state-based decision-making.

---

The ability to change behavioral responses to information collected from the world is a defining feature of motivation^1^. Studies of motivational scaling have focused primarily on the potentiation of sensory stimuli when our internal states make us more likely to act on them^2–11^, perhaps in part because sensory responses are reliable and easily measured. Sensory scaling would likely help animals to identify relevant environmental stimuli, but many signals are either ambiguous or relevant to multiple motivations, requiring modifications at other sites of control. By localizing a motivated decision to a small group of neurons governing an innate behavior and subjecting them to extensive electrical and molecular manipulations, we find another solution to the problem of motivational response tuning. We find that motivation biases a decision by altering the time over which information is retained and allowed to accumulate in sensory-agnostic decision-making neurons.

*Drosophila* copulation provides a clear and quantitative readout of motivated decision-making. Mating precludes many other behaviors, so to respond to changing conditions flies must first stop mating. Undisturbed copulations last ∼23 minutes, but if a dangerous situation arises the male^12–14^ may decide to truncate the mating (**Figure 1a, Extended Data Figure 1f**), depending on both the severity of the threat and how far the mating has progressed. For the first several minutes after initiating copulation, the male will sacrifice his (and his partner’s) life in the face of a potentially lethal threat, presumably to ensure successful fertilization. But his persistence (or propensity to keep mating when challenged) decreases as time passes, reflecting the increasing likelihood that the goals of mating have been achieved^12–14^.

**Figure 1:**
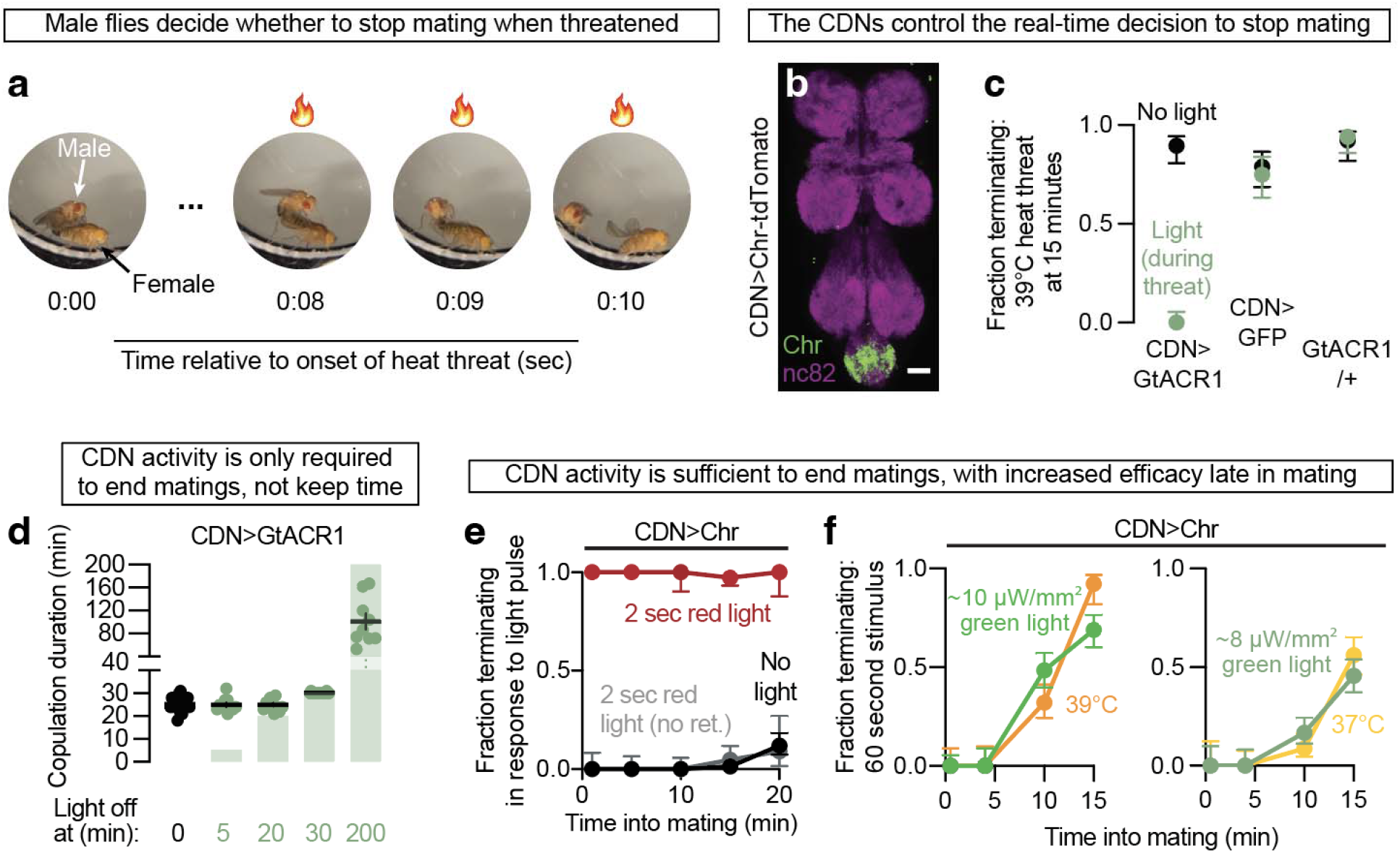
Copulation Decision Neuron (CDN) activity controls the real-time decision to end matings. **(a)** Male flies decide whether to stop mating when threatened. The male (red eyes) maintains a stereotyped posture while mating. When challenged (in this case with a 41°C threat) he may decide to detach himself from the female (yellow eyes) and end the mating. **(b)** The CDNs (labeled by NP2719-Gal4^12^) reside in the abdominal ganglion of the ventral nervous system (neuropil at bottom of image). Scale bar is 20 μm. **(c)** Acute silencing of the CDNs using the light-gated chloride channel GtACR1 prevents termination in response to a one-minute-long 39°C heat threat. Error bars for proportions here and in all other figures (unless otherwise stated) represent 67% credible intervals, chosen to resemble the standard error of the mean. For the number of samples in each experiment, see **Supplementary Table 3**. For statistical tests, see **Supplementary Table 4**. **(d)** Electrical activity in the CDNs is only necessary at the time of mating termination. Tonic silencing of the CDNs results in extended mating duration (fifth column, mean: 101 minutes), but silencing from the beginning until near the natural end of mating does not affect copulation duration (third column). Matings in which the CDNs are silenced through the normal ∼23-minute termination time end seconds after the light is turned off (fourth column). Green rectangles represent the time during which the neurons were silenced. Error bars for copulation duration here and throughout represent standard error of the mean. **(e)** Acute optogenetic stimulation of the CDNs causes termination. Two seconds of stimulation is sufficient to terminate copulation regardless of how far the mating has progressed. “No ret.” refers to flies that were not fed retinal, the obligate chromophore for CsChrimson’s light sensitivity, showing that light on its own does not cause termination of mating. **(f)** The termination response to minute-long green light stimulation (green lines; Left: 9.72 μW/mm^2^, Right: 8.03 μW/mm^2^) of the CDNs is potentiated as a mating progresses, similar to the response to real-world challenges like heat threats (orange lines).

We previously used the motivational dynamics of this system to study how neurons measure time on the scale of minutes^13^. That mechanism centered on the self-sustained activity of CaMKII, achieved through autophosphorylation at residue T287, a feature intensely studied for its role in memory formation and storage in the mammalian hippocampus^15,16^. Here, in a different set of neurons, we find another long-timescale function for CaMKII: matching decision-making to motivational state by scaling the amount of time over which information is integrated. These findings broaden the implications for CaMKII in regulating phenomena over timescales longer than can be supported by electrical activity alone^17,18^.

There are three main findings in this study: 1) that the changing integrative properties of the sexually dimorphic Copulation Decision Neurons (CDNs) underlie the male fly’s decrease in persistence as mating progresses; 2) that the timescale of integration is set by levels of CaMKII activity; and 3) that motivating dopaminergic signaling increases CaMKII’s excitability in the CDNs, restricting integration time and steer the resolution of conflicting drives. This mechanism supports flexible decision-making by allowing decision-relevant information to accumulate over timescales that depend on the relative values of the behaviors under consideration.

## RESULTS

### Copulation Decision Neurons (CDNs) set the probabilistic behavioral response to threats during mating

Our previous work showed that synaptic release of GABA from NP2719-Gal4 neurons (a set of abdominal ganglion interneurons referred to here as the CDNs; **Figure 1b, Extended Data Figure 1a, Extended Data Figures 16-20, Supplementary Discussion 1**) is required for matings to end at the appropriate time^12^ but with the tools available at the time could not resolve their role in the immediate decision to terminate the mating (**Supplementary Note 1**). Here we explored the relationship between acute CDN activity and decision-making. Silencing the CDNs with GtACR1^19^ (a light-gated chloride channel) during the presentation of a heat threat caused the male to persist through a strong challenge that would otherwise end nearly all matings (**Figure 1c** and **Video 1**). In unchallenged matings, tonic CDN inhibition extended copulation duration well beyond the natural ∼23-minute mark (**Figure 1d**), but if the inhibition ended after 20 minutes, matings were of normal duration. Tonic inhibition that continued beyond 23 minutes resulted in near-instantaneous termination after the inhibition concluded (**Figure 1d**). Inversely, CDN silencing that began just before 23 minutes extended matings as much as if the neurons had been inhibited throughout the mating (**Extended Data Figure 1b**). These results show that the electrical activity of the CDNs is not required to keep time during mating — only for the decision to end it, whether in response to threats or at its ∼23-minute conclusion.

Two seconds of optogenetic CDN stimulation using the red-light gated cation channel CsChrimson (Chr)^20^ terminated nearly 100% of matings (**Video 2** and **Figure 1e**) with a dismounting procedure resembling the response to threatening stimuli (**Video 3**). Termination occurred regardless of time into mating and with a variable latency of up to 30 seconds after the stimulation pulse (**Extended Data Figure 1d**). CDN stimulation in non-mating flies did not induce any obvious change in behavior and did not stop flies from courting or initiating mating (**Extended Data Figure 1c**), arguing against a startle reflex or activation of a motor pattern generator. Silencing the CDNs immediately after stimulation prevented terminations that had not already occurred (**Extended Data Figure 1e**), suggesting a requirement for sustained CDN electrical activity to make and execute the decision to switch behaviors.

The results from these initial optogenetic experiments were all-or-nothing, either suppressing the fly’s response to threats 100% of the time or inducing termination 100% of the time. But the fly’s responses to natural stimuli are probabilistic, with a gradually increasing fraction of matings being truncated in response to identical stimuli as the mating progresses^12^. If the CDNs were to adjust their output in response to direct stimulation according to mating time, then they might be the locus at which the termination probability is rescaled. To provide direct input to the CDNs without immediately terminating matings we delivered green light to mating pairs, which is less effective than red light at stimulating CsChrimson^20^ and slightly less effective at penetrating the fly’s cuticle^21^. The efficacy of minute-long green light CDN stimulation increased as mating progressed, with each condition quantitatively matching the response of a real-world threat of fixed severity (**Figure 1f**). These data argue that the CDNs treat sustained artificial stimulation similarly to their natural inputs, even though the excitation is non-synaptic and not matched to specific physiological dynamics. Since these experiments completely bypass sensory input by directly stimulating the CDNs, our results argue against an important role for motivational amplification of sensory stimuli upstream of the CDNs. Instead, together with the molecular results below, they argue that motivational scaling occurs within the CDNs.

### The CDNs more effectively retain termination-promoting information later into mating

In contrast to minute-long real-world threats and optogenetic stimulation bouts, termination to brief (500 ms or 1 sec), strong (i.e., bright red light) optogenetic pulses did not change from 10 to 15 minutes into mating (**Figure 2a**). The importance of stimulus duration in elevating termination probability late in mating raised the idea that the CDNs may change the timescale over which they integrate information rather than perceived stimulus intensity as mating progresses.

**Figure 2:**
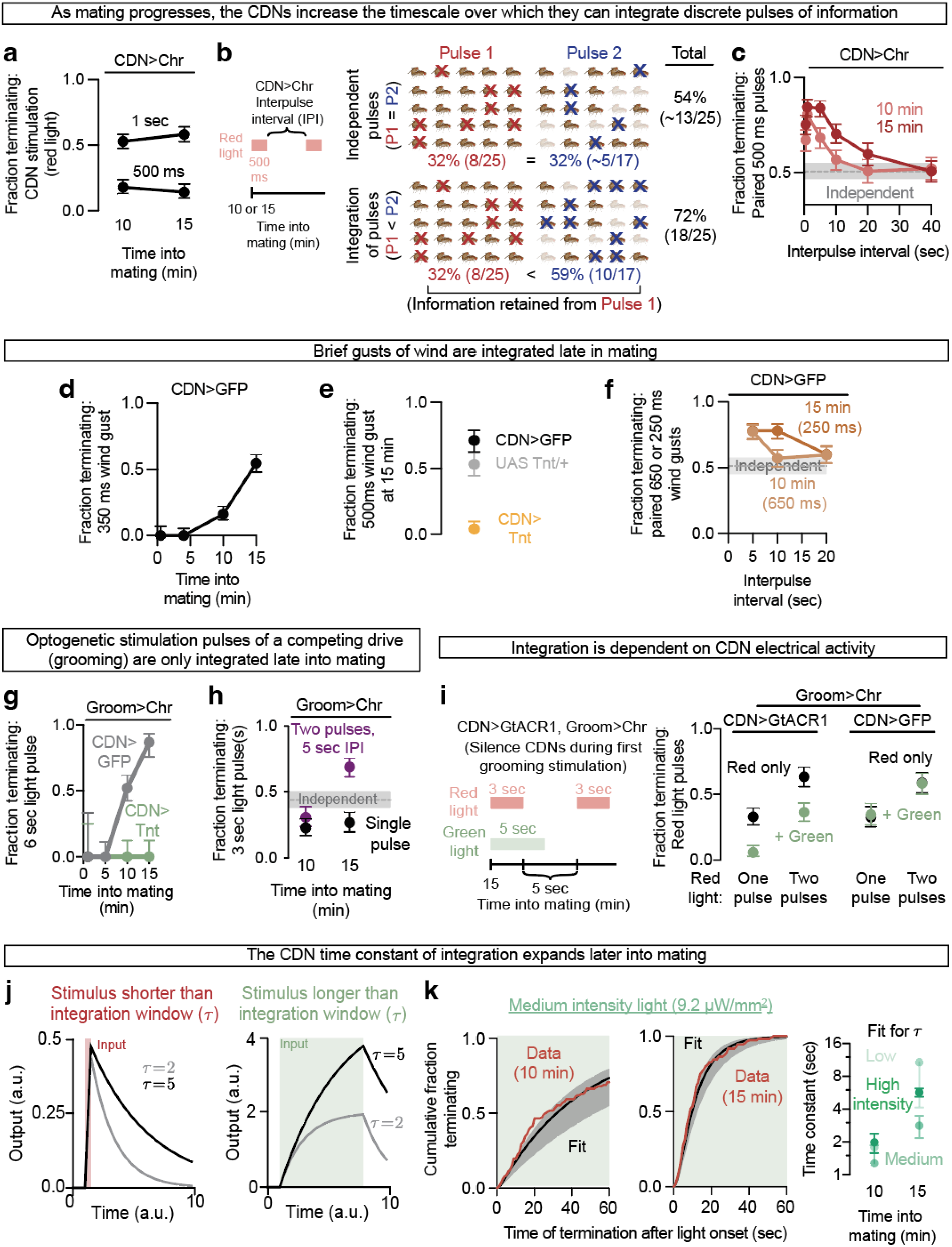
The CDNs integrate multimodal inputs over longer timescales as mating progresses. **(a)** Unlike longer challenges, the termination response to brief (500 ms and 1 sec), strong pulses of CDN stimulation does not increase as mating progresses. **(b)** Left: flies are given two strong 500 ms pulses of CsChrimson stimulation separated by an inter-pulse interval. Right: illustration of the probability of terminating the mating in response to two independent (top) or integrated (bottom) pulses. In the independent case, each pulse (first, red; second, blue) would terminate ∼32% of matings, and so in total would terminate ∼54% of matings. If information from the first pulse is integrated into the response to the second, then the overall proportion terminating the mating will be > 54% (in this example, 72% where the second pulse is augmented to 59%). **(c)** The probability of terminating the mating in response to two strong pulses of CDN stimulation is greater than expected if the pulses did not interact (grey line = 2p-p^2^, estimated using the single pulse response value (p)) so long as the inter-pulse interval is sufficiently short. The ability of pulses to be integrated over longer timescales is increased at 15, relative to 10, minutes into mating. **(d)** Male flies become progressively more likely to stop mating in response to a 350-millisecond gust of wind. Mating pairs are not blown apart by the force of the wind; rather, the male terminates the mating with some delay after the end of the wind gust (**Extended Data Figure 2** and see example in **Video 4**). **(e)** Constitutively silencing the CDNs with tetanus toxin (Tnt) prevents termination in response to wind (see example in **Video 5**). **(f)** Paired pulses of wind gusts (650 ms at 10 min, 250 ms at 15 min) are integrated over a longer timescale when delivered at 15 minutes into mating. **(g)** Six seconds of optogenetic grooming neuron stimulation causes termination with increasing propensity as the mating progresses. Grooming-induced termination is prevented at all time points by blocking CDN output with tetanus toxin (Tnt). **(h)** Two bouts of brief grooming neuron stimulation separated by 5 seconds results in a potentiation of the response to the second pulse, but only at 15 minutes into mating. A single brief pulse terminates the same fraction of matings at 10 and 15 minutes. The predicted value if the two pulses did not interact is indicated by the grey line. **(i)** Silencing the CDNs during a first demotivating stimulus prevents its integration with a later stimulus. Left: two bouts of grooming neuron stimulation were delivered at 15 minutes into mating, separated by 5 seconds. The CDNs were silenced during the first pulse of grooming neuron stimulation, as well as 2 seconds after the pulse offset. Right: silencing the CDNs during only the first of two optogenetic grooming pulses induces the same level of termination as if only one pulse (with no CDN inhibition) had been delivered. **(j)** Response of a model system to input pulses of equal strength but varying duration. The instantaneous probability of terminating a mating in response to a stimulus is: *p*_0_*τ* (1 − exp(− *t*/*τ*)). *p*_0_= perceived intensity, *τ* = time constant of integration, t = time since start of stimulus. Left: for pulses shorter than the integration window, the peak response is approximately the same regardless of *τ*, but the amount of information retained after pulse offset is greater with a greater *τ*. Right: for pulses longer than the integration window, information accumulates faster and peaks higher with a greater τ. Error bars represent estimated pointwise 95% coverage intervals. **(k)** Plotting the cumulative termination after the onset of green light and fitting the data (left) reveals a higher time constant of integration, *τ*, at 15 minutes into mating compared to 10 minutes (right). Error bars represent one standard error of the parameter fit estimated using the Cramér-Rao bound (see **Supplementary Note 2, Extended Data Figure 5**,**6**).

To examine the role of integration time in information processing and rule out confounding effects of sustained stimulation on the CsChrimson protein itself, we separated two, strong, 500 millisecond stimulation pulses by various time intervals (**Figure 2b**). These two pulses resulted in termination rates well above the expectation if each pulse were acting independently, but only when the pulses were presented sufficiently close together in time (**Figure 2c**). At 15 minutes into mating, the two pulses were integrated over a longer time interval than at 10 minutes, showing that integration cannot be accounted for by the dynamics of CsChrimson alone^20^ and instead arguing that information is retained over a longer time window as the mating progresses.

Because CsChrimson delivers depolarizing currents throughout the cell, rather than specifically at postsynaptic sites, we designed a preparation using naturalistic stimuli that could be delivered in quick pulses. We delivered sub-second gusts of wind to mating pairs (**Extended Data Figure 2,3**), as we had previously demonstrated that sustained exposure to intense airflow could interrupt mating^12^. Individual bursts of wind became more and more effective in causing termination as the mating progressed (**Figure 2d**), an effect that required CDN output (**Figure 2e**). Like direct stimulation of the CDNs with optogenetics, brief gusts of wind integrated supralinearly, and the time over which the pulses could interact increased as mating progressed (**Figure 2f**).

Like heat and wind threats, optogenetic stimulation of neurons that promote grooming behavior^22^ causes flies to terminate at higher levels when delivered later in mating, and this termination relies on CDN output (**Figure 2g**). Two short pulses of grooming neuron excitation were only integrated when presented later into mating, consistent with a gradually expanding integration window (**Figure 2h**). If the CDNs were silenced during the first grooming pulse, the second pulse acted as though the first had not been present at all (**Figure 2i**), demonstrating that the lingering information integrated from the first pulse required electrical activity of the CDNs, rather than acting in upstream circuitry or on the CsChrimson protein itself. We found similar mating time-dependent pulse integration by stimulating a separate neuronal population^23^ whose activity terminated mating (**Extended Data Figure 4**). These results argue that the CDNs integrate competing information from a variety of neuronal inputs — and that the temporal window of integration expands as motivation to sustain the mating decreases.

These results all indicate that the difference between the male’s higher motivational state at 10 minutes and lower motivational state at 15 minutes cannot be explained by a simple rescaling of synaptic inputs or a change in excitability. Instead, they suggest that the ability of the CDNs to retain information about demotivating inputs later in mating is a primary determinant of the fly’s decision-making under duress.

### Modeling the response to sustained inputs

For sustained inputs like our heat threats, increasing the amount of time over which information can accumulate would lead to pronounced differences in the response to a stimulus (schematized in **Figure 2j**). A longer integration time predicts that sustained stimulation would end matings not only more frequently, but also earlier into the stimulation, due to surpassing the saturation point of conditions with shorter time constants. Scoring the exact times of termination within minute-long optogenetic stimulation bouts at 10 or 15 minutes into mating supported this hypothesis (**Figure 2k**). We used the resulting cumulative termination curves to fit a simple linear dynamical system to produce quantitative estimates of information decay. The modeling estimated time constants of information retention, *τ* ≈ 1-2 seconds at 10 minutes and ≈ 3-10 seconds at 15 minutes (**Figure 2k, Supplementary Note 2, Extended Data Figures 5, 6, Methods** *[Modeling]*), in general agreement with our experimental findings above and below. r corresponds to the duration it takes each moment of input signal to decay by 1-1/e ≈ 63%, consistent with a discernible effect of integration as long as inputs are separated by less than 2r.

**Figure 3:**
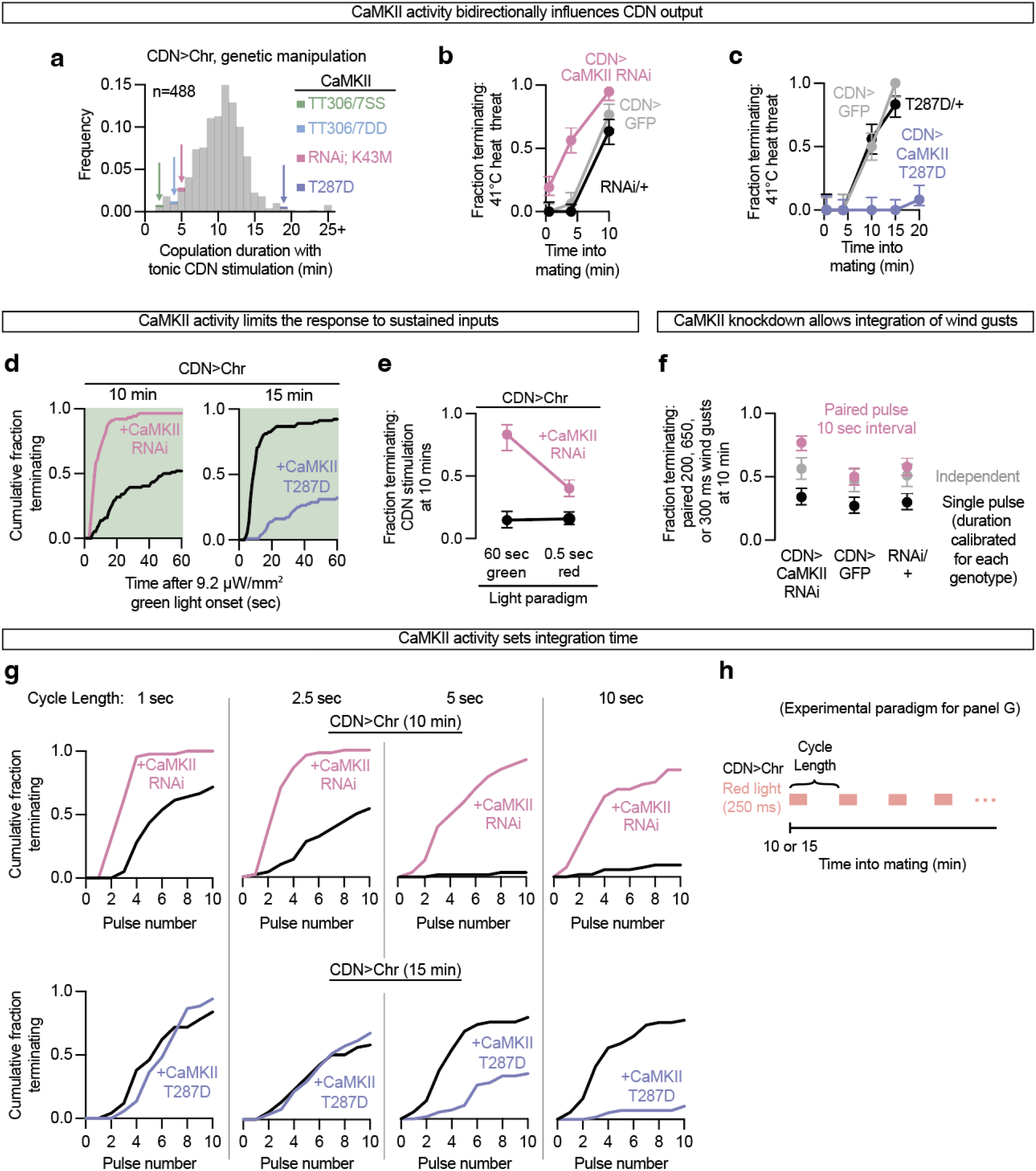
CaMKII activity in the CDNs sets the timescale of integration. **(a)** When the CDNs are tonically stimulated with white light throughout courtship and mating, copulation duration is reduced to ∼11 minutes. Out of ∼500 genetic manipulations (mostly RNAi), several manipulations of CaMKII in the CDNs strongly altered copulation duration. The average duration of each genotype (consisting of at least 6 flies) is rounded down and binned to a whole number, and bins containing CaMKII manipulations are indicated with colored arrows. **(b)** Knocking down CaMKII in the CDNs of males increases the sensitivity of matings to heat threats early into mating. **(c)** Expressing constitutively active CaMKII (T287D) in the CDNs protects matings from heat threats even late into mating. **(d)** Decreasing CaMKII activity via RNAi increases the rate and overall fraction of flies terminating to green light at 10 minutes whereas increasing CaMKII activity with T287D expression decreases the rate and overall fraction of flies terminating to green light at 15 minutes. **(e)** Knocking down CaMKII more strongly potentiates the termination response to sustained green light stimulation (8.03 μW/mm^2^) than a short pulse of red-light stimulation, though each protocol terminates the same proportion of control flies. **(f)** Knocking down CaMKII in the CDNs allows paired wind gusts to be integrated across a 10 second inter-gust interval at 10 minutes into mating. Single pulse lengths were calibrated so that flies would terminate at a rate of ∼30% for each genotype. **(g)** Cumulative fraction of flies terminating after each light pulse at 10 (top) and 15 (bottom) minutes into mating. At both 10 and 15 minutes into mating, males integrate pulses that are separated by less than 2.5 seconds, but integration across longer intervals was seen only at 15 minutes. CaMKII knockdown allows flies to integrate pulses at 10 minutes as if it were later into mating, while constitutively active CaMKII suppresses the time over which the CDNs can integrate pulses. **(h)** Ten 250 millisecond red light pulses with a set amount of time between each pulse were delivered to CDN>CsChrimson flies at either 10 or 15 minutes into mating.

**Figure 4:**
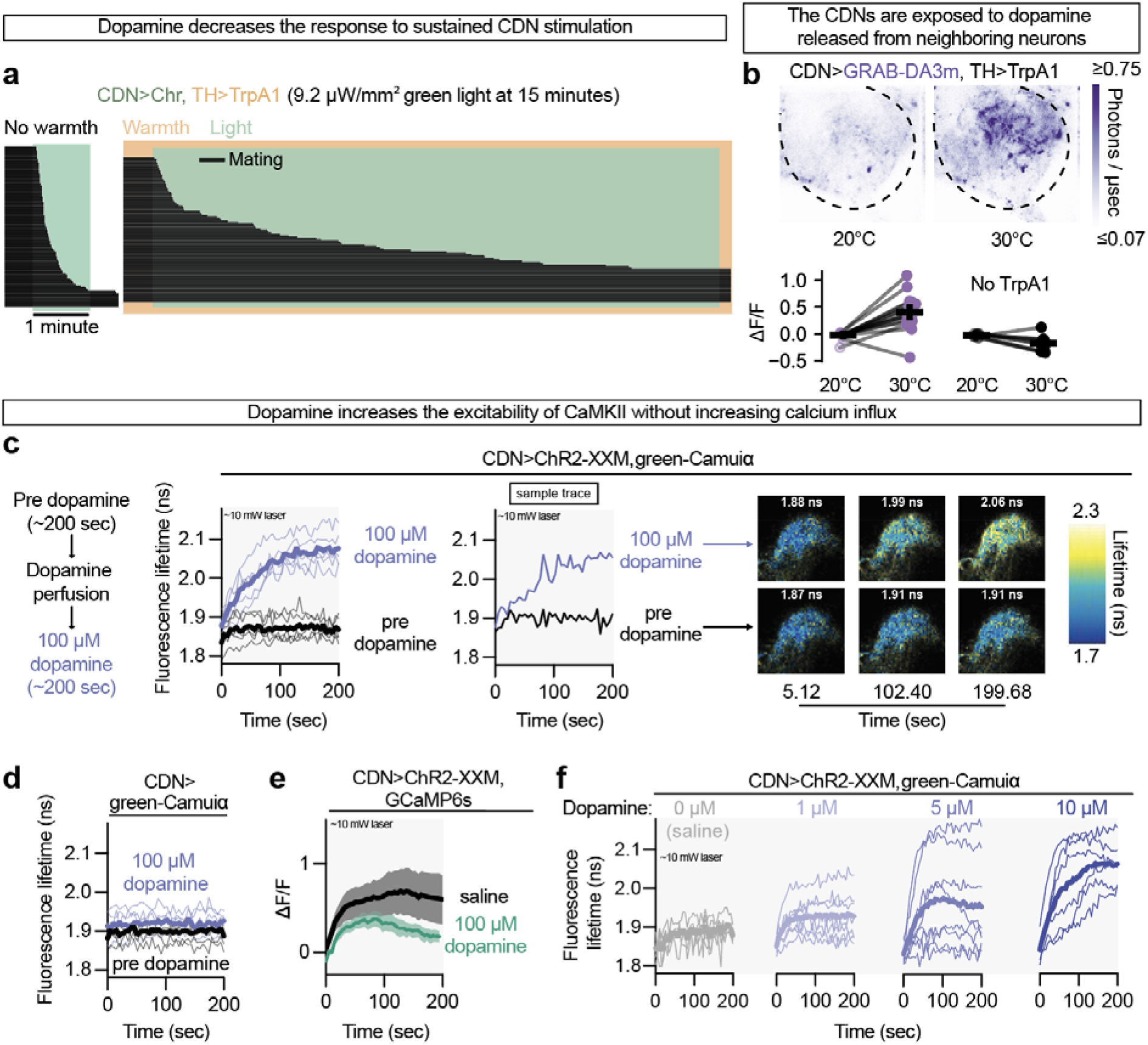
Dopamine restricts integration by facilitating CaMKII activation by calcium. **(a)** Dopamine motivates matings by restricting integration by the CDNs. Sustained thermogenetic stimulation of dopaminergic neurons with TrpA1 protects the mating against optogenetic stimulation of the CDNs at 15 minutes into mating. Each black stripe in the ethograms represents a single mating. **(b)** Thermogenetic stimulation of dopaminergic neurons drives release of dopamine onto the CDNs as measured by changes in the brightness of the genetically encoded fluorescent dopamine reporter GRAB-DA3m. Top: example abdominal ganglion (outlined with a black dashed line) at room temperature, with dim GRAB-DA3m fluorescence (left), and the same AG after increasing the temperature to stimulate local dopaminergic neurons (right). Bottom shows quantification of changes in fluorescence with temperature in flies expressing TrpA1 in the dopaminergic neurons (purple) or without TrpA1 (black). **(c)** Dopamine promotes the activation of CaMKII as reported by the fluorescence lifetime of the FRET sensor green-Camuiα. Left: ∼10-milliwatt, 920 nm laser stimulation of the Channelrhodpsin-2 variant ChR2-XXM only slightly increases CaMKII activity in the CDNs (black); subsequent perfusion of 100 μM dopamine allows the same stimulation of ChR2-XXM to strongly increase CaMKII activity (blue). NP5270-Gal4^12^ is used as the CDN driver for green-Camuiα and GCaMP6s imaging experiments (see **Methods** *[Imaging experiments, Region of interest]*). Each bold trace is the mean of the light traces. Middle: A representative sample trace. Right: fluorescence lifetime map of green-Camuiα signal in CDN axons from the example trace before and after dopamine perfusion. **(d)** Dopamine cannot increase CaMKII activity without ChR2-XXM stimulation. **(e)** Dopamine does not increase calcium influx in the CDNs as measured by GCaMP6s. Baseline fluorescence is calculated as the mean number of photons for the first 5.12 seconds of pre-perfusion recording, which was done at the same 10-milliwatt laser power (see **Methods** *[Imaging experiments, Optogenetic stimulation while imaging/Calcium imaging]*). **(f)** Increasing dopamine concentration increases the ability of ChR2-XXM stimulation to activate CaMKII. Each trace is a single fly. For the 0 μM concentration, saline was perfused while imaging.

**Figure 5:**
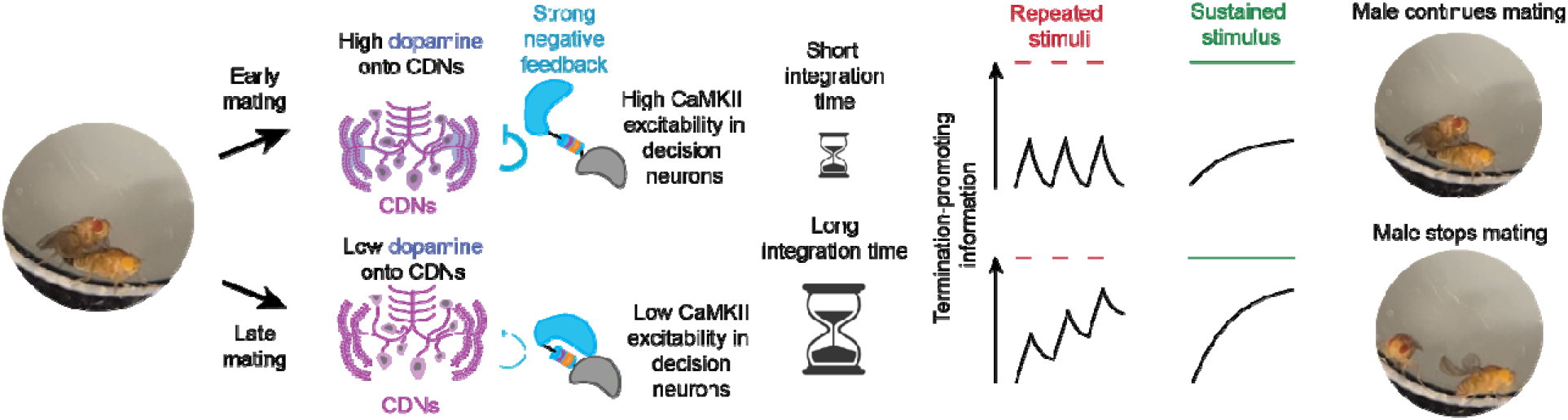
A motivating dopamine signal acts through CaMKII to bias behavioral choice by controlling retention of decision-relevant information.

### CaMKII activity adjusts the retention of information

We performed a screen for modifiers of CDN output in response to sustained, white-light stimulation, which on average took ∼11 minutes to terminate matings (**Figure 3a; see Methods** *[Behavioral experiments, Optogenetic stimulation during behavior, Screen experiments]* **and Supplementary Note 1**). Several of the most prominent outliers involved manipulations of the highly-conserved^24^ calcium/calmodulin-dependent kinase CaMKII. When a constitutively active CaMKII mutant^25^ (the phosphomimetic point mutation T287D) was expressed in the CDNs, matings were far less sensitive to sustained optogenetic stimulation (**Figure 3a**). Inversely, when CaMKII activity was reduced in the CDNs via an RNAi transgene, expression of a kinase dead mutant^26^ (K43M), or of mutated versions expected to reduce its activation by calmodulin binding^27^ (TT306/7DD, TT306/7SS), matings were more sensitive to sustained stimulation (**Figure 3a**). These results led us to investigate the role of CaMKII activity on temporal integration in the CDNs.

In assessing the role of CaMKII in the response to challenges, we found that RNAi-mediated knockdown in the CDNs made matings more susceptible to heat threats (**Figure 3b**), without affecting either the duration of undisturbed matings or the requirement for electrical activity to truncate matings (**Extended Data Figures 7a, b**). The constitutively active T287D mutation made matings insensitive to heat or wind threats (**Figure 3c, Extended Data Figure 8g**), effects that required CaMKII kinase activity (**Extended Data Figure 8b**) and had no bearing on the male’s fertility (**Extended Data Figure 8c**). Some males expressing T287D in the CDNs mated for over an hour (**Extended Data Figure 8d**), similar to males in which the CDNs were electrically or synaptically^12^ silenced, but expression of CaMKII-T287D had little to no effect on the ability of CsChrimson stimulation to evoke calcium transients (**Extended Data Figure 8a**). Males expressing the T287D mutation readily terminated matings in response to repeated CDN stimulation (see below, **Figure 3g**), indicating that the CDNs are still functional in the presence of sustained CaMKII activity. Using our cumulative termination paradigm, we found that increasing CaMKII activity decreased the rate at which males terminated matings in response to 1-minute green light stimulation, making them behave as if it were earlier into mating. Inversely, decreasing CaMKII levels made males behave as if it were later in mating (**Figure 3d**). These experiments suggest that CaMKII may exert control over the termination decision by decreasing the timescale of integration in the CDNs.

In another test of whether CaMKII controls the integration process or simply dampens CDN responses regardless of the input dynamics, we selected two conditions of optogenetic excitation that differed in intensity and duration by orders of magnitude but were balanced so that they were equally effective in terminating matings (∼15% termination probability). Knockdown of CaMKII increased the response to 60 seconds of weak stimulation far more effectively than the response to 500 milliseconds of strong excitation (**Figure 3e**). These results are inconsistent with a model in which CaMKII activity only impacts the magnitude of CDN output. Instead, they argue that CaMKII’s primary role is to control integration over many seconds.

Perhaps the most convincing demonstration of temporal integration in the CDNs is the ability of information from a first stimulatory input to combine with a later pulse to increase termination probability, as in **Figure 2b**. We performed a version of this experiment with wind gusts, finding a similarly increasing ability for spaced gusts to integrate later into mating. Knocking down CaMKII in the CDNs lengthened the integration window, allowing pulses to integrate when separated by 10 seconds at 10 minutes into mating (**Figure 3f**) — much longer than the normal time constant of integration at that point.

We noticed that the termination probability in response to single 500 ms red light pulses was altered by knockdown of CaMKII. This effect is explainable by adjustments to integration time, but it confounds a direct comparison across genotypes (**Extended Data Figure 9a, b**). We decreased the stimulation pulse length to 250 ms, which on its own almost never terminated matings (**Extended Data Figure 9c**) and supplied 10 pulses with varying cycle lengths — the rationale being that this provides many cycles over which a weak stimulus can accumulate (if the interpulse interval is less than the integration time of the system) without immediately saturating our readout (**Figure 3g**) (albeit with additional caveats, **Extended Data Figure 13**). At 10 minutes into mating, these 250 ms pulses accumulated to cause termination only when separated by less than 5 seconds. At 15 minutes into mating, integration was seen with up to 10 seconds between pulses (**Figure 3f**).

When CaMKII was knocked down, we found evidence of integration at 10 minutes into mating even when pulses were separated by as much as 10 seconds (**Figure 3f**). When CaMKII activity was increased in the CDNs via expression of the T287D mutant, pulses accumulated supralinearly with 1 second or 2.5 second cycle lengths, again showing that CaMKII does not simply prevent CDN output. Elevated CaMKII activity did, in contrast, suppress the accumulating response to pulses separated by 5 or 10 seconds (**Figure 3g**), arguing that the behavioral consequences of CaMKII activity are strongly dependent on the temporal dynamics of input to the CDNs. These results lead us to conclude that the CDNs are the locus of integration for termination-promoting information and that CaMKII activity adjusts the rate of information decay over time to scale behavioral responses.

### Dopamine facilitates CaMKII activation to prevent the accumulation of competing information

In the linear model, saturation and integration are both caused by negative feedback, with weak negative feedback corresponding to a longer time constant and higher saturation point. CaMKII is classically activated by calcium and calmodulin, and so would be poised to implement negative feedback – and manipulation of its calcium-dependent activation by mutating the 306 and 307 residues strongly changed the response to optogenetic excitation in our initial screen (**Figure 3a**). Because integration relies on the activity of the CDNs themselves rather than just receiving input (**Figure 2i**), we speculated that motivational inputs might adjust the magnitude of gain by affecting the relationship between CaMKII and activity of the CDNs, just as one would adjust the feedback term of the linear model.

Dopaminergic activity in the ventral nervous system (anatomically distinct from the courtship promoting dopaminergic neurons in the brain, **Extended Data Figure 11e**) motivates males to persist through threats during mating^12^ by signaling through the D2 receptor on the CDNs [Miner et. al^28^]. Thermogenetic stimulation of dopaminergic neurons using the warmth-gated cation channel TrpA1^29^ dramatically increased the duration of sustained optogenetic CDN stimulation required to end matings (**Figure 4a**), suggesting that dopamine might motivate matings by restricting CDN integration, possibly via CaMKII. Thermogenetically stimulating dopaminergic neurons while expressing the dopamine sensor GRAB-DA3m^30^ caused a temperature-dependent increase in fluorescence (**Figure 4b**) and FLIM signal (**Extended Data Figure 14**), arguing that activity-dependent dopamine secretion in the abdominal ganglion is capable of elevating dopamine levels to physiologically-meaningful levels near the CDNs.

In our previous work^13,14^ we noticed that the calcium-permeable and highly sensitive channelrhodopsin variant ChR2-XXM^31^ was activated by the near-infrared laser at high power, so we had kept power low to avoid stimulating the opsin while imaging. Here, we increased the laser power to provide constant stimulation of the CDNs while simultaneously monitoring calcium or CaMKII activity levels (see **Methods** *[Imaging experiments, Optogenetic stimulation while imaging]*). While this stimulation increased calcium levels in the CDNs as reported by GCaMP6s^32^ (see below), it had little effect on CaMKII activity (**Figure 4c**). However, when we perfused dopamine into the bath (at concentrations producing similar GRAB-DA3m FLIM changes as thermogenetic stimulation of dopaminergic neurons, **Extended Data Figure 14**), CaMKII was strongly activated by the same stimulation (**Figure 4c**). Dopamine perfusion also potentiated CaMKII activation in response to transient one-photon stimulation of ChR2-XXM (**Extended Data Figure 10b**). Perfusing dopamine while providing laser stimulation did not increase CaMKII activity in a different set of neurons, arguing against a generic effect of dopamine on ChR2-XXM or its relationship to CaMKII excitability. (**Extended Data Figure 10c**). CaMKII activity quickly relaxed back to baseline after stimulation ceased, illustrating that the sustained increase in CaMKII activity was maintained by the ongoing elevation of calcium (**Extended Data Figure 10d**), supported by the absence of a phenotype from preventing calcium-independent activity with the T287A mutation in **Figure 3a**. Perfusing dopamine without co-expression of ChR2-XXM only slightly increased fluorescence lifetime (**Figure 4d**), arguing against an effect of dopamine on the green-Camuiα sensor itself, and leading us to conclude that dopamine acts to enhance the ability of calcium to activate CaMKII in the CDNs.

An alternative hypothesis to explain these results might be that dopamine increases calcium influx, which in turn activates CaMKII. However, dopamine did not cause an increase in calcium driven by ChR2-XXM laser stimulation. Instead, high concentrations of dopamine decreased stimulation-evoked calcium in the CDNs despite effectively activating CaMKII (**Figure 4e**, further quantified in **Extended Data Figures 10e**,**f;** [Miner et. al^28^]). Increasing concentrations of dopamine increased the ability of ChR2-XXM stimulation to activate CaMKII (**Figure 4f**). These results support a model in which CaMKII acts as a rapid negative feedback regulator of CDN output, capping the retention of information supplied by excitatory inputs. In conditions of low dopamine, CaMKII is less activated by CDN calcium, allowing longer information retention and rendering the mating more susceptible to threats (**Figure 5**).

## Discussion

Compiling information over time is at the core of decision-making^33,34^, so control over the integration of that information seems at least as likely a target for motivational regulation as sensory processing—and it has several advantages. For example, control of decision-making via adjustments to the accumulation of information at decision centers avoids amplifying upstream signals, including decision-irrelevant inputs. The integration mechanism we propose serves as a tunable lowpass filter, promoting decisions based on accumulated evidence collected over time, instead of transient noise. This mechanism may be particularly well-suited to making decisions during innate behaviors, where the relevant sensory stimuli are more complex and varied than the learned sensory association tasks where decision-making is most frequently examined.

Our results, and work on sexually dimorphic fly behaviors more broadly^12–14,35–41^, add a new perspective to the prevailing view of decision-making as the result of anonymized population-level attractor dynamics^42^. In conventional analyses of circuit dynamics, such as region-wide measurements of intracellular calcium, CDN-like populations would likely not stand out among thousands of intermingled cells. The ability to identify and manipulate compact circuit elements provides clear evidence of decision-making bottlenecks that should be considered in the interpretation of widespread calcium or voltage recordings. Even having identified the CDNs, our data illustrate that calcium imaging would fail to reveal their changing integrative properties over the course of mating. We see the identification of compact decision-making centers as an opportunity to reduce the complexity of high-level processes and work to establish core molecular principles that enrich our understanding beyond a circuit-level explanation.

Our results add to a growing understanding of the neuronal populations that control the male’s changing responsiveness to challenges over time during mating (**Figure 5, Extended Data Figure 12, Supplementary Discussion 1**). A critical moment occurs six minutes after mating begins when the Crz eruption drives sperm transfer and opens the period of threat responsiveness^13,14^. The eruption is timed from within the Crz neurons by CaMKII’s self-sustained but slowly declining activity via autophosphorylation at T287. In this work we show that CaMKII works in a different set of neurons to again control long-timescale behavioral changes, but with key differences. In the CDNs, it is not clear to what extent CaMKII uses its classic self-sustaining abilities at T287. Instead, dopamine signaling guides the ability of CaMKII to be activated by excitatory inputs to the CDNs. Instead of delaying an all-or-nothing event as in the Crz neurons, CaMKII activity appears to work in real time to adjust the window of integration in the CDNs to match the local dopamine tone and guide the male’s response to challenges that arise during mating.

We believe that we have i) found strong evidence that motivational state tunes temporal integration to bias decision outcomes, ii) discovered the anatomical locus of this integration, and iii) identified key molecular factors that make the adjustment. But the physical nature of the information integrated over time and the means by which dopamine influences CaMKII activatability are not yet clear. One potential interaction between dopamine and CaMKII may come from the two phosphorylatable threonines at the 306 and 307 sites in the calmodulin binding region. Phosphorylation at these positions prevents calmodulin binding, and so decouples CaMKII activation from calcium/calmodulin levels. If dopamine signaling were to reduce phosphorylation rates at T306 and T307, it could directly control the relationship between CaMKII activity and electrical activity. Expression of mutant CaMKII proteins which mimic phosphorylation at these sites caused low motivation-like phenotypes in our screen.

Calcium levels could be the target of CaMKII activation as well as its impetus, matching the predictions of the linear model, in which negative pressure on the accumulating variable is produced by the accumulating variable itself (*dy*/*dt* = -*y/τ*). Lingering calcium over seconds-long timescales is also known to influence synaptic release in a process known as synaptic facilitation^43^, providing another means of controlling CDN output. But CaMKII manipulations with powerful impacts on integration produced only small changes in measured calcium after optogenetic stimulation (**Extended Data Figure 8a**), mostly visible near the bottom of the sensor’s dynamic range^32^. However, the calcium influx critical for controlling synaptic release is thought to be dominated by local conductances at the active zone^44,45^, and may only minimally contribute to our bulk cytosolic GCaMP measurements. Understanding how CaMKII activity expedites the purging of information will also require deeper investigation into this system. Even lacking these important details, our results provide a new and unexpected mechanistic explanation for motivational control over decision making. Given the broad conservation of the molecules involved and the generality of the problem, we expect this way of thinking about motivation to be useful in understanding motivated decisions across behaviors and animals.

## Supporting information

Full Supplement

## Acknowledgements

We thank: Dragana Rogulja for discussions, comments on the manuscript, and hosting us in her lab for the early stages of this project; Ryohei Yasuda and Long Yan for assistance building the two-photon microscope used in these experiments; the following undergraduates for assistance performing experiments related to or inspiring those in this work - Jiaxiang Zhang, Liam Kerrick, Roshinie Persaud, Peter Rifkin, Ashna Singh, Megan Hoffman, Evan Zheng, Mary Dello Russo, Gabriel Verderame; Barret Pfeiffer, David Anderson, and Gerry Rubin for sharing the UAS and LexAop2 Cs-Chrimson-tdTomato stocks before publication; Ofer Mazor and Pavel Gorelik (Harvard Medical School Neuroinstrumentation Core) for technical advice on designing the experimental apparatuses; Jazz Weisman for assistance machining the *ex vivo* physiology apparatus in Figure 4b and Extended Data Figure 14 and Ezgi Hacisuleyman for assistance stabilizing the preparation; Gaby Maimon for hosting SCT during the revision of this work; Yulong Li for sharing the 10x-UAS-GRAB-DA3m plasmid ahead of publication; members of the Maimon Lab for feedback on concepts and experiments; and current and former members of the Crickmore and Rogulja Labs for comments on the manuscript. This work was funded by the NIH (R01NS111441). SCT was supported by the National Science Foundation Graduate Research Fellowship (DGE1144152) and Helen Hay Whitney Postdoctoral Fellowship. LEM was supported by the National Science Foundation Graduate Research Fellowship (DGE2140743).

## Author Contributions

AKG and SCT performed the behavioral experiments, except **Figure 3a** which was performed by LEM. Fly genetics were performed by AKG, SCT, and MAC. The wind presentation chambers were designed and built by AKG. AKG performed the imaging experiments, except those in **Figure 4b** and **Extended Data Figure 14** which were performed and analysed by SCT. MANC EM analysis was performed by SCT. Statistical analysis code was written by SCT. Linear systems analyses and modeling were performed by SCT. MNF wrote the FLIM analysis code (based on advice from SCT). AKG, SCT, and MAC wrote the paper, with input from LEM and MNF.

## Notes

### Competing Interest Statement

The authors have declared no competing interest.

## References

1. Flavell, S. W., Gogolla, N., Lovett-Barron, M. & Zelikowsky, M. The emergence and influence of internal states. Neuron vol. 110 2545–2570 Preprint at 10.1016/j.neuron.2022.04.030 (2022).

2. Hindmarsh Sten, T., Li, R., Otopalik, A. & Ruta, V. Sexual arousal gates visual processing during Drosophila courtship. Nature 595, 549–553 (2021).

3. Ko, K. I. et al. Starvation promotes concerted modulation of appetitive olfactory behavior via parallel neuromodulatory circuits. Elife 4, (2015).

4. Aton, S. J. Set and setting: How behavioral state regulates sensory function and plasticity. Neurobiol Learn Mem 106, 1–10 (2013).

5. Lange, R. D. & Haefner, R. M. Characterizing and interpreting the influence of internal variables on sensory activity. Curr Opin Neurobiol 46, 84–89 (2017).

6. McLachlan, I. G. et al. Diverse states and stimuli tune olfactory receptor expression levels to modulate food-seeking behavior. 11, 79557 (2022).

7. Vogt, K. et al. Internal State Configures Olfactory Behavior and Early Sensory Processing in Drosophila Larvae. Sci. Adv vol. 7 https://www.science.org (2021).

8. Richman, E. B., Ticea, N., Allen, W. E., Deisseroth, K. & Luo, L. Neural landscape diffusion resolves conflicts between needs across time. Nature 623, 571–579 (2023).

9. Wu, Z. et al. Context-Dependent Decision Making in a Premotor Circuit. Neuron 106, 316-328.e6 (2020).

10. Burgess, C. R. et al. Hunger-Dependent Enhancement of Food Cue Responses in Mouse Postrhinal Cortex and Lateral Amygdala. Neuron 91, 1154–1169 (2016).

11. Allen, W. E. et al. Thirst regulates motivated behavior through modulation of brainwide neural population dynamics. Science (1979) 364, (2019).

12. Crickmore, M. A. & Vosshall, L. B. Opposing dopaminergic and GABAergic neurons control the duration and persistence of copulation in drosophila. Cell 155, 881 (2013).

13. Thornquist, S. C., Langer, K., Zhang, S. X., Rogulja, D. & Crickmore, M. A. CaMKII Measures the Passage of Time to Coordinate Behavior and Motivational State. Neuron 105, 334-345.e9 (2020).

14. Thornquist, S. C., Pitsch, M. J., Auth, C. S. & Crickmore, M. A. Biochemical evidence accumulates across neurons to drive a network-level eruption. Mol Cell 81, 675-690.e8 (2021).

15. Yasuda, R., Hayashi, Y. & Hell, J. W. CaMKII: a central molecular organizer of synaptic plasticity, learning and memory. Nature Reviews Neuroscience vol. 23 666–682 Preprint at 10.1038/s41583-022-00624-2 (2022).

16. Silva, A. J., Paylor, R., Wehner, J. M. & Tonegawa, S. Impaired Spatial Learning in α-Calcium-Calmodulin Kinase II Mutant Mice. Science (1979) 257, 206–211 (1992).

17. Sebastian Seung, H., Lee, D. D., Reis, B. Y. & Tank, D. W. Stability of the Memory of Eye Position in a Recurrent Network of Conductance-Based Model Neurons. Neuron 26, 259–271 (2000).

18. Major, G. & Tank, D. Persistent neural activity: prevalence and mechanisms. Curr Opin Neurobiol 14, 675–84 (2004).

19. Mohammad, F. et al. Optogenetic inhibition of behavior with anion channelrhodopsins. Nat Methods 14, 271–274 (2017).

20. Klapoetke, N. C. et al. Independent optical excitation of distinct neural populations. Nat Methods 11, 338–346 (2014).

21. Inagaki, H. K. et al. Optogenetic control of Drosophila using a red-shifted channelrhodopsin reveals experience-dependent influences on courtship. Nat Methods 11, 325–332 (2014).

22. Seeds, A. M. et al. A suppression hierarchy among competing motor programs drives sequential grooming in Drosophila. Elife 3, e02951 (2014).

23. Namiki, S., Dickinson, M. H., Wong, A. M., Korff, W. & Card, G. M. The functional organization of descending sensory-motor pathways in Drosophila. (2018) doi:10.7554/eLife.34272.001.

24. Tombes, R. M., Faison, M. O. & Turbeville, J. M. Organization and evolution of multifunctional Ca2+/CaM-dependent protein kinase genes. Gene vol. 322 17–31 Preprint at 10.1016/j.gene.2003.08.023 (2003).

25. Park, D., Coleman, M. J., Hodge, J. J. L., Budnik, V. & Griffith, L. C. Regulation of neuronal excitability in Drosophila by constitutively active CaMKII. J Neurobiol 52, 24–42 (2002).

26. Hanson, P. I., Meyer, T., Stryer, L. & Schulman, H. Dual Role of Calmodulin in Autophosphorylation of Multifunctional CaM Kinase May Underlie Decoding of Calcium Signals. Neuron vol. 12 (1994).

27. Elgersma, Y. et al. Inhibitory Autophosphorylation of CaMKII Controls PSD Association, Plasticity, and Learning. Neuron 36, (2002).

28. Miner, L. E., Gautham, A. K. & Crickmore, M. A. Local desensitization to dopamine devalues recurring behavior. BioRxiv doi:10.1101/2024.02.20.581276.

29. Hamada, F. N. et al. An internal thermal sensor controlling temperature preference in Drosophila. Nature 454, 217–220 (2008).

30. Zhuo, Y. et al. Improved green and red GRAB sensors for monitoring dopaminergic activity in vivo. Nat Methods (2023) doi:10.1038/s41592-023-02100-w.

31. Scholz, N. et al. Mechano-dependent signaling by Latrophilin/CIRL quenches cAMP in proprioceptive neurons. Elife (2017) doi:10.7554/eLife.28360.001.

32. Chen, T. W. et al. Ultrasensitive fluorescent proteins for imaging neuronal activity. Nature 499, 295–300 (2013).

33. Huk, A. C. & Shadlen, M. N. Neural activity in macaque parietal cortex reflects temporal integration of visual motion signals during perceptual decision making. Journal of Neuroscience 25, 10420–10436 (2005).

34. Wang, S., Faeder, J. R., Setlow, P. & Li, Y. Q. Memory of germinant stimuli in bacterial spores. mBio 6, (2015).

35. Wang, K. et al. Neural circuit mechanisms of sexual receptivity in Drosophila females. Nature 589, 577–581 (2021).

36. Kimura, K. ichi, Hachiya, T., Koganezawa, M., Tazawa, T. & Yamamoto, D. Fruitless and Doublesex Coordinate to Generate Male-Specific Neurons that Can Initiate Courtship. Neuron 59, 759–769 (2008).

37. Hoopfer, E. D., Jung, Y., Inagaki, H. K., Rubin, G. M. & Anderson, D. J. P1 interneurons promote a persistent internal state that enhances inter-male aggression in Drosophila. (2015) doi:10.7554/eLife.11346.001.

38. Feng, K., Palfreyman, M. T., Häsemeyer, M., Talsma, A. & Dickson, B. J. Ascending SAG neurons control sexual receptivity of Drosophila females. Neuron 83, 135–148 (2014).

39. Zhou, C., Pan, Y., Robinett, C. C., Meissner, G. W. & Baker, B. S. Central brain neurons expressing doublesex regulate female receptivity in Drosophila. Neuron 83, 149–163 (2014).

40. Zhang, S. X., Miner, L. E., Boutros, C. L., Rogulja, D. & Crickmore, M. A. Motivation, Perception, and Chance Converge to Make a Binary Decision. Neuron 99, 376-388.e6 (2018).

41. Kallman, B. R., Kim, H. & Scott, K. Excitation and inhibition onto central courtship neurons biases Drosophila mate choice. doi:10.7554/eLife.11188.001.

42. Ebitz, R. B. & Hayden, B. Y. The population doctrine in cognitive neuroscience. Neuron vol. 109 3055–3068 Preprint at 10.1016/j.neuron.2021.07.011 (2021).

43. Jackman, S. L. & Regehr, W. G. The Mechanisms and Functions of Synaptic Facilitation. Neuron vol. 94 447–464 Preprint at 10.1016/j.neuron.2017.02.047 (2017).

44. Südhof, T. C. Calcium control of neurotransmitter release. Cold Spring Harb Perspect Biol 4, (2012).

45. Zucker, R. S. & Regehr, W. G. Short-term synaptic plasticity. Annual Review of Physiology vol. 64 355–405 Preprint at 10.1146/annurev.physiol.64.092501.114547 (2002).

46. Takemura, S.-Y. et al. A Connectome of the Male Drosophila Ventral Nerve Cord. BioRxiv (2023) doi:10.1101/2023.06.05.543757.

47. Cheong, H. S. J. et al. Transforming descending input into behavior. BioRxiv doi:10.1101/2023.06.07.543976.

48. Tayler, T. D., Pacheco, D. A., Hergarden, A. C., Murthy, M. & Anderson, D. J. A neuropeptide circuit that coordinates sperm transfer and copulation duration in Drosophila. Proc Natl Acad Sci U S A 109, 20697–20702 (2012).

49. Randi, F., Sharma, A. K., Dvali, S. & Leifer, A. M. Neural signal propagation atlas of Caenorhabditis elegans. Nature 623, 406–414 (2023).

50. Talay, M. et al. Transsynaptic Mapping of Second-Order Taste Neurons in Flies by trans-Tango. Neuron 96, 783-795.e4 (2017).

51. Simões, J. M. et al. Robustness and plasticity in Drosophila heat avoidance. Nat Commun 12, (2021).

52. Mussells Pires, P., Zhang, L., Parache, V., Abbott, L. F. & Maimon, G. Converting an allocentric goal into an egocentric steering signal. Nature 626, 808–818 (2024).

53. Nern, A., Pfeiffer, B. D. & Rubin, G. M. Optimized tools for multicolor stochastic labeling reveal diverse stereotyped cell arrangements in the fly visual system. Proc Natl Acad Sci U S A 112, E2967–E2976 (2015).

