## Supplementary material for "Molecular control of temporal integration matches decision-making to motivational state": Full Supplement

### SUPPLEMENTARY DISCUSSION

##### Supplementary Discussion 1

##### Connectomic Analysis

We first manually examined each of the 704 neurons that only receive input within the AG, have release sites only within the AG, and do not have processes that travel through any nerves. We annotated whether they superficially resembled the Crz neurons or the CDNs, taking into consideration where they appear to have synaptic release sites (for example, the Crz neurons have release sites throughout their arbors (**Extended Data Fig 11a**), while the CDNs seem to restrict release sites to the lateral portions of the abdominal ganglion (**Extended Data Fig. 1a**)). All neurons of interest to this study, except possibly some dopaminergic neurons, also have somata restricted to the posteriormost abdominal neuromeres (the anteriormost neuromeres are dominated by neurons innervating the metathoracic ganglion). We arrived at a list of 39 potential Crz neurons and 142 potential CDNs. There appear to be several morphologically-distinct populations of dopaminergic neurons in the AG, and we have not yet managed to use genetic tools to discern which dopaminergic neurons were likely to be important, so we did not attempt to identify them in this first stage.

*EM Characterization of the putative Crz neurons:*

We then attempted to narrow down which of these 39 cells are the four Crz neurons, the most visually distinct of the abdominal ganglion neurons we have studied, based on a number of measures. First, we looked more closely at each neuron's anatomy for the following features: 1) arbors innerving the lateral portion of both sides of the AG, with a single thick neurite crossing through the center, 2) a single thin but short neurite extending up towards more anterior portions of the AG on one side only, 3) cell bodies located at the posterior end of the AG neuropil and 4) that the ventral portion of the neurites on each half of the AG did not cross or meet in the middle (though the dorsal portion often does), 5) identifiable dense core vesicles in the EM volume. In the end, we found 6 neurons that appeared to be Crz-like (**Extended Data Figure 15 a,a**), all of which innervated one another densely. This similarity was reflected in the serial set assignment of these neurons, an automated measure produced by the MANC annotation intended to coarsely cluster cells with similar synaptic partners[46,47], with all putative Crz neurons being assigned to serial set "11828". The six neurons split into two apparent groups by multiple analyses: cosine similarity of their synaptic output vectors (which compares whether they innervate the same general populations of neurons), coarse morphological parameters such as overall counts of synaptic input and output and volume, and predicted neurotransmitter profiles (**Extended Data Figure 15 c,d**). Both groups also strongly innervated a set of large descending neurons in the EM volume (a subset of the EN00B001 superclass) that resembled the CrzR neurons described in Tayler *et al*., 2012 known to drive ejaculation[48] (**Extended Data Figure 15e**). We presumed the set of four cells was the Crz neurons. Even still, no set of neurons convincingly matches the description of four identical cells – this suggests that our selection of putative Crz neurons is incorrect, that the Crz neurons can be further differentiated into two pairs of two, or that there are six Crz-like cells only four of which express Crz itself.

Of their postsynaptic neurons, their output synapses were divided into: onto one another (~5% of synapses), onto the two other Crz-like neurons (~5% of synapses), onto putative CrzR neurons (~8% of synapses), onto other neurons descending the abdominal trunk nerve (~19%), local cholinergic neurons (~20%), local GABAergic neurons (~7%), local glutamatergic neurons

(3.15%), local neurons of uncertain release type (~20%), neurons innervating at least one other neuropil (~20%), and synapses onto ascending neurons (<1%) (**Extended Data Figure 15f**). Their outputs appear broadly distributed at the level of synaptic connectivity, making many connections across many cell classes, consistent with serving a complex role in regulating copulation. They appear not to be a major synaptic input of, nor receive inhibition from, the CDNs themselves despite the proximity of Crz release sites to CDN neurites (**Extended Data Figure 11a**). An important caveat in interpreting these data, however, is that peptidergic signaling is not expected to be constrained to synapses as revealed, for example, in functional connectivity mapping in *C. elegans*[49]. Their fast excitatory neurotransmission likely is important in driving ejaculation through serotonergic descending neurons[48], but their effect on the fly's motivational state may act through other means. Their action on the CDNs' integrative properties resulting in a long-lasting switch in their excitability after the transient eruption of Crz activity (**Extended Data Figure 11c,d**) is inconsistent with typical predictions for direct synaptic connections, and we do not interpret the lack of apparent connectivity here as a lack of functional interaction (even ignoring the possibility of incorrect identification of either the Crz neurons or the CDNs).

The inputs to the Crz neurons are considerably simpler: 16% of their input synapses come from four neurons with highly similar anatomy (**Extended Data Figure 15g**), which we speculate to be the Dh44 neurons of the abdominal ganglion that reliably mark the Crz neurons when using *trans-Tango* labeling[50] driven with Dh44-Gal4 (unpublished). The next greatest contribution comes from the Crz neurons themselves, with 9% of their input synapses coming from other Crz neurons (consistent with previously-described reciprocal functional innervation[14]) (**Extended Data Figure 15g**). 94% of all input synapses are from local neurons of the AG, breaking down in a similar fashion to the outputs of the Crz neurons: 24% are of unknown transmitter type, 10% are predicted to be GABAergic, 7.5% are presumed glutamatergic, and 26% are assessed as cholinergic. The actual cell classes making up these remaining synapses are very distributed, consisting of many neurons each forming comparatively few synapses.

*EM characterization of the putative CDNs:*

The CDNs as a group show consistent morphological features from fly to fly but seem less consistent on a single neuron basis (**Extended Data Figure 16a**). However, all appear to have small somata, about half of which are in the ventral AG and half in the dorsal AG, receive sizable synaptic input in the central AG in thin neurites crossing the midline in segments (resembling a ladder), and have release sites in the AG's dorsolateral extent. Most CDNs appear to have unilateral projections (**Extended Data Figure 1a**), and IHC and genetic perturbations suggest most CDNs are GABAergic[12]. Published accounts vetting machine learning-based neurotransmitter classifications have shown them to be unreliable outside of the class of neurons on which they were trained, and in our own experience they have frequently mislabeled non-cholinergic cell types (unpublished), but for some cell classes (e.g. the ring neurons of the ellipsoid body), they have been extremely robust, and so we opted to use neurotransmitter classification as an initial screening condition. Of the 142 potential CDNs we identified in the EM volume, 54 are predicted to be GABAergic with high confidence.

While each CDN shows separate release sites in the dorsolateral abdominal ganglion, because they all appear to share the same input structure we first compared neurons by the cosine similarity of the vectors of presynaptic inputs to their postsynaptic sites. Ordering this similarity matrix using hierarchical clustering showed that most GABAergic neurons of the AG have unique collections of presynaptic inputs, but about half of the neurons could be sorted into one of five groups (**Extended Data Figure 16**). For each of these clusters, we examined the morphology of the individual neurons more closely. In clusters 1 and 2, each neuron had neurites in the center of the AG at which they received much of their inputs and release sites in the dorsolateral AG, though each neuron's morphology was idiosyncratic and the cell body positions were not stereotyped (**Extended Data Figure 16**). Cluster 2 consisted of 7 neurons, and ~8 of the 12 neurons labeled by the CDN driver line are expected to be GABAergic[12], so we labeled Cluster 2 "putative CDN"s. Because Cluster 1's outputs were less consistent than Cluster 2's (**Extended Data Figure 16**) and their morphology was also somewhat less reliably structured but reminiscent of the CDNs, we opted to label Cluster 1 as "CDN-like". The CDNs' output vectors should also appear to split into two similar groups, as Cluster 2 did (**Extended Data Figure 16**). We continue to use Clusters 3-5 in the subsequent analyses for comparison with Clusters 1 and 2.

Of the five clusters of GABAergic interneurons, 3-5 showed seemingly high direct recurrence, but Clusters 1 and 2 did not. We computed the Spearman correlation between the number of synapses a GABAergic interneuron made on a target cell against the number of synapses it received from that same cell. Clusters 1 and 2 had a Spearman correlation of 0.06 and 0.05, respectively, while Clusters 3, 4, and 5 had correlations of .19, 0.21, and .11, reflecting a much greater tendency to receive comparable direct feedback from a neuron that cells in those three clusters innervated (**Extended Data Figure 17a**). Pooling within a cluster, however, tells a different story: if the correlation is computed between total synapses from a cluster onto a postsynaptic cell and how many synapses that cell makes back onto any cell in the cluster, the correlation increases to match that of the other groups (**Extended Data Figure 17b**). However, Clusters 1 and 2 have a particularly large set of cells that either synapse far more onto the cluster than they receive or receive far more synapses than they reciprocate (long horizontal and vertical lines near the axes); the vast majority of the reciprocal pairings are from neurons that are only weakly connected to the CDNs.

We then examined the individual identities of the postsynaptic connections of the presumed CDNs. The major outputs were one of three classes: 1) inter-neuropil interneurons that only receive input in the abdominal neuromere and have release sites in the three leg neuropils, 2) large bilateral local interneurons, and 3) descending motorneurons through the abdominal trunk nerve (**Extended Data Figure 18a**).

The majority of the interneuropil neurons targeted by the CDNs project to multiple leg neuropil, and those which receive the most input from the CDNs (with the exception of two cells) have very few pre-synaptic sites in the abdominal ganglion (**Extended Data Figure 18b**). Most of these neurons themselves are excitatory and target either motorneurons of the leg neuropils or interneurons of those neuropil. In the case of targeted interneurons, the interneurons targeted were predicted to be either glutamatergic or GABAergic, in either case likely inhibiting the local circuitry controlling movement of that leg. These point to a complex relationship between CDN activity and the movements of the fly, plausibly driving escape or dismounting motor programs coordinated across legs, inhibiting specific patterns of activity aimed at keeping the male balanced on the female to encourage dismounting, or anything in between.

The large bilateral local interneurons all showed highly similar morphology: each neuron's processes filled both hemispheres of the abdominal ganglion with almost no innervation of the midline through its entire dorso-ventral extent (**Extended Data Figure 18d**). About half of these interneurons were putatively inhibitory and half putatively excitatory (**Extended Data Figure 18e**), with the strongest innervation being of cholinergic interneurons that did not innervate the CDNs highly, arguing against a direct role of the CDNs in computing information through recurrent processing. They did, however, innervate GABAergic neurons to a lesser extent which themselves synapse onto the CDNs, potentially allowing recurrent excitation through disinhibition. The confidence of the neurotransmitter prediction for many of these recurrent neurons was comparatively low, usually predicting GABA with less than twice the likelihood of either other option. These neurons may be peptidergic or aminergic (which the classifier does not include as categories), confounding direct interpretations of the sign of this recurrent connection. Unlike the other two classes of downstream cells, the CDNs made up a sizable fraction of synaptic inputs to many of these interneuron classes, in some instances constituting >10% of their total synaptic input (**Extended Data Figure 18**). Even those neurons where the total number of synapses from the CDNs was relatively small (e.g. those with too few synapses to be included in **Extended Data Figure 18a**) received a large fraction of their synaptic input from the CDNs (**Extended Data Figure 18**). It is likely that the CDNs play an important role in regulating activity in the abdominal ganglion through these neurons, while their effects on other neuropil and motor output is less substantive. In short, these data do not unequivocally rule out a role for recurrence in the integrative properties of the CDNs, but they do not tell a story in which that is a dominant feature of their connectivity.

A minority of the CDNs synapses targeted varied descending motor neurons. Of these, about 7 received sizable input comparable with interneuropil and abdominal ganglion interneurons (**Extended Data Figure 18a**). Unlike the other two postsynaptic cell classes, in no case did the CDNs make up more than a few % of the synaptic input of these motor neurons, and even those cases were rare (**Extended Data Figure 18**). The CDNs may contribute to the actions controlled by neurons descending the abdominal trunk nerve, such as those controlling the genitalia, but seem unlikely to be a primary determinant in their output.

Trans-Tango labeling of putative postsynaptic targets partially corroborates this analysis, labeling 6 large AG interneurons with cell bodies on the ventral face of the ganglion resembling the major interneuron targets of the CDNs, with a large gap in the center of the AG and dense innervation of both sides (**Extended Data Figure 18h**). One faint dorsal ascending neuron appears to be labeled as well, but no descending axons were visible. Because most descending and interneuropil targets receive only a very small fraction of their inputs from the CDNs, it is perhaps unsurprising that they are not easily visualized with this approach, even though the numerosity of the synapses from the CDNs is relatively high (in a few cases, several hundreds of synapses).

Finally, we examined the classes of neurons which innervate the CDNs. These break down into four main classes: descending neurons coming from the brain (making up ~8.5% of total synapses onto the CDNs), interneuropil neurons coming from the leg neuropils of the VNS (24.1%), local interneurons of the AG (52.1%), and ascending neurons climbing the abdominal nerve (6.7%) (**Extended Data Figure 19a**) Most of these neurons innervated all of the CDNs to a similar extent, despite the varied morphology of each CDN, or innervated about half of the CDNs (if they targeted a single half of the AG) (**Extended Data Figure 19b**).

The principal descending neuron classes innervating the putative CDNs all share one morphology: they descend the VNS along a dorsomedial tract, innervating the leg neuropil to a small extent but with the majority of their output synapses in the abdominal neuromere (**Extended Data Figures 19c,d**). For many of these descending neurons, the CDNs are their primary targets as well, in most cases making up nearly half of all of their output synapses in the abdominal neuromere (**Extended Data Figure 19e**). The identity of these cell types, in terms of known brain descending neurons, is currently unknown, and the main descending neurons are annotated as DNx177, DNx071, DNx094, DNxl130, and DNad002. They possibly convey threat information directly, but because of their small fractional input to the CDNs, are presumably not the sole source of threat (or other mating demotivating) signals.

The interneuropil neurons were similarly focused on the abdominal ganglion, with modest innervation of one or two other neuropil (usually T3, **Extended Data Figure 19f,g**) and heavy innervation of the AG. Of these, many synapsed onto the CDNs seemingly proportionate to their number of synapses overall, but one pair of interneurons seemed highly specific, almost exclusively synapsing onto the 7 cells annotated as CDNs with their 400 synapses (**Extended Data Figure 19h**), and these synapses were broadly distributed across the CDNs (**Extended Data Figure 19i**). These neurons were also the strongest singular inputs onto the CDNs (**Extended Data Figure 19a**). A second pair, the second strongest inputs to the CDNs, provide almost as many synapses, but a lower fraction of their total synaptic output (**Extended Data Figure 19l**). Because these interneuropil cells seem to constitute a direct line to the CDN population, we provide a cursory examination of their inputs as well. Each of these neurons receives a much-higher-than-typical fraction of their input synapses from descending neurons (**Extended Data Figure 19j**), and many of the remaining synapses come from large local interneurons of the other leg neuropil. These inputs to the CDN appear to aggregate information coming from many sources, possibly providing access to some of the combined demotivating input the CDNs would need to integrate to perform their generic demotivating function.

The anatomy of the interneurons targeting the CDNs is highly varied. We plot the top 24 interneuron inputs to the CDNs in **Extended Data Figure 20a**, showing that there is no consistent anatomy between them. We speculated that a subset of these may be the dopaminergic neurons, and that they might be identified by the inability of the machine learning classifier to categorize their synapses. Two major inputs of the putative CDNs had near-total uncertainty in their synapse classification, as measured by the entropy of the prediction probabilities (**Extended Data Figure 20b**), with two others showing only modest confidence. We suspect either of these pairs may be relevant sources of dopamine for the CDNs. These cells may also be peptidergic or secrete other monoamines, and suggest there remain modulatory inputs to the CDNs we have not yet characterized. The majority of the inputs to the CDNs, especially strong inputs, were predicted to be GABAergic or glutamatergic, and very few of these connections received input from the CDNs. Together, these facts suggest they do not principally receive excitation within the AG, even from disinhibiting recurrent AG connections (as speculated above) and rely on the interneuropil connections and descending neurons to increase their activity (**Extended Data Figure 20c**).

### SUPPLEMENTARY NOTES

#### Supplementary Note 1

In our previous work[12] we showed that thermogenetically stimulating the CDNs throughout courtship and copulation shortened mating duration (an effect we here recreate with optogenetic stimulation, **Extended Data Figure 1c**), leading us to hypothesize that electrical activity of the CDNs is responsible for timing the duration of a mating. However, a thorough optogenetic interrogation of the CDNs (**Figure 1**) reveals that electrical activity is only necessary at the time of termination. We do not understand why flies can habituate to many minutes of intense CDN stimulation if it begins before mating, but not if it begins after mating starts. We used a modified version of this paradigm as a first-pass screen for genetic manipulations that would alter the effectiveness of CDN stimulation to end matings (**Figure 3a**) because it is higher throughput than manually delivering heat or light to flies at a certain time into mating – it is possible that some molecular hits from the screen may specifically control this habituation property of the CDNs.

#### Supplementary Note 2

Increasing the amount of time over which information accumulates would lead to pronounced differences in the response to a stimulus, but only when the input is sustained for a period comparable to the duration over which information can be retained. In the simplest case, a linear dynamical system with a "time constant" parameter $\tau$ (i.e. $dy/dt = -y/\tau + f(t)$ with $f(t)$ the time-varying stimulus) allows a single quantitative variable to stand in for this memory-like capacity, where $\tau$ would vary slowly over many minutes of mating (so it can be modeled as approximately constant for any individual stimulus). This model recapitulates our findings in which strong but short-lived inputs — ones much shorter than even the smallest values of $\tau$, such as the 500 millisecond pulse — show little change in the probability of terminating mating between presentation at 10 minutes and 15 minutes (**Figure 2a**), while weak inputs on the order of $\tau$ or longer scale termination response dramatically (**Figure 1f**).

To derive quantitative estimates for the time constant of integration in different motivational states we modeled the instantaneous probability of terminating mating with the linear dynamical system above, so that $y(t) = probability(\text{terminating mating at time } t \text{ if still mating by time } t)$, presuming that the nonlinearity above operates near its linear regime with very weak stimuli. The precise control over the onset times and intensity of input in the sustained optogenetic stimulation experiments enabled us to fit the cumulative distribution of actual times of termination to the equation

$$\sigma(t) = 1 - exp\left[-p_0\left(t - \tau\left(1 - e^{-\frac{t}{\tau}}\right)\right)\right]$$

(where $p_0$ is a constant corresponding to the strength of the demotivating input to the CDNs) to jointly estimate the parameters of the model (**Extended Data Figure 2a**, see **Methods** for more information). A shorter $\tau$ caps the peak termination probability for long threats of fixed intensity, whereas extending integration to longer timescales leads to an increased output and, therefore, increased termination probability. Increases in $p_0$, presuming that the stimulus itself is identical, would reflect an increasing sensitivity to all stimuli regardless of their dynamics. Using this model allows explicit quantitative reasoning about the extent to which temporal integration changes across experimental conditions and across time.

We plotted the cumulative termination probabilities during varying intensities of CDN stimulation beginning either 10 or 15 minutes into mating (**Extended Data Figure 2d)**. These plots show that the data are well fit by a linear integration process in time, with $\tau \approx$ 1-2 seconds at 10 minutes and $\approx$3-10 seconds at 15 minutes (**Extended Data Figure 2b**). Importantly, without either constraint being explicitly imposed, the model fit only predicted a change in the overall perceived strength of input when the intensity of the light was changed (**Extended Data Figure 2b**; $p_0 \approx 10^{-3}$ for low intensity light, 0.02 for medium intensity, and 0.1 for high intensity). The fit is orders of magnitude better when temporal integration is included than if the flies are assumed not to integrate inputs over time (**Extended Data Figure 2c**).

### EXTENDED DATA FIGURES

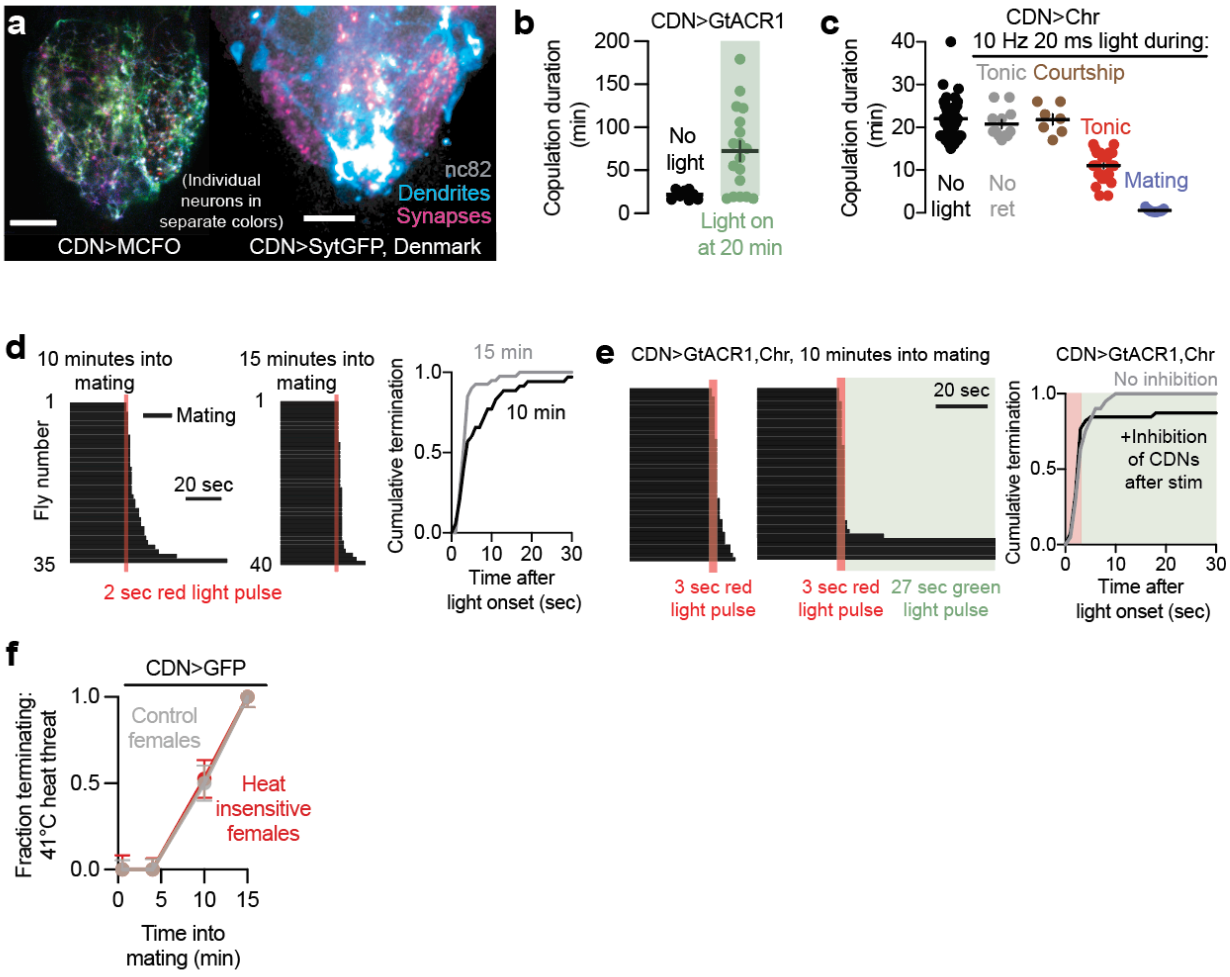


**Extended Data Figure 1: Sustained CDN activity is necessary and sufficient to end matings**
**(a)** Left: Individual CDNs labelled via MultiColor FlpOut (image of a single optical section of the abdominal ganglion). Right: CDN dendrites selectively cover the midline tracts of the abdominal ganglion (blue); the CDNs send axonal projections throughout the abdominal ganglion (magenta). Scale bars are 20 μm.
(**b**) Electrical activity of the CDNs is only necessary around the time of termination to end the mating: silencing that begins just before the natural time of termination (20 min) is still sufficient to prolong the mating – 5 flies stopped mating before the light was turned on.
**(c)** Optogenetic stimulation of the CDNs using CsChrimson preceding the onset of mating does not affect copulation duration if only supplied during courtship (brown), but shortens copulation by several minutes if continued into the mating (i.e. flashing red light throughout the duration of the experiment, “tonic”, red). These results closely resemble the results of thermogenetic activation in previous work[12] that did find immediate termination of the mating upon CDN activation. Providing the same optogenetic activation only after mating begins results in near-immediate termination of copulation (blue).
(**d**) Flies end matings in response to 2 seconds of optogenetic CDN stimulation with varying latencies. Left: ethogram, right: cumulative distribution plot.

**(e)** Stimulation of the CDNs followed by immediate electrical silencing largely prevents the termination of matings that had not ended before the silencing began. Left: ethogram, right: cumulative distribution plot.
**(f)** Mating with a heat-insensitive, Gr28b.d;TrpA1 double mutant female[51] does not change the male's decision to stop mating when threatened by heat.

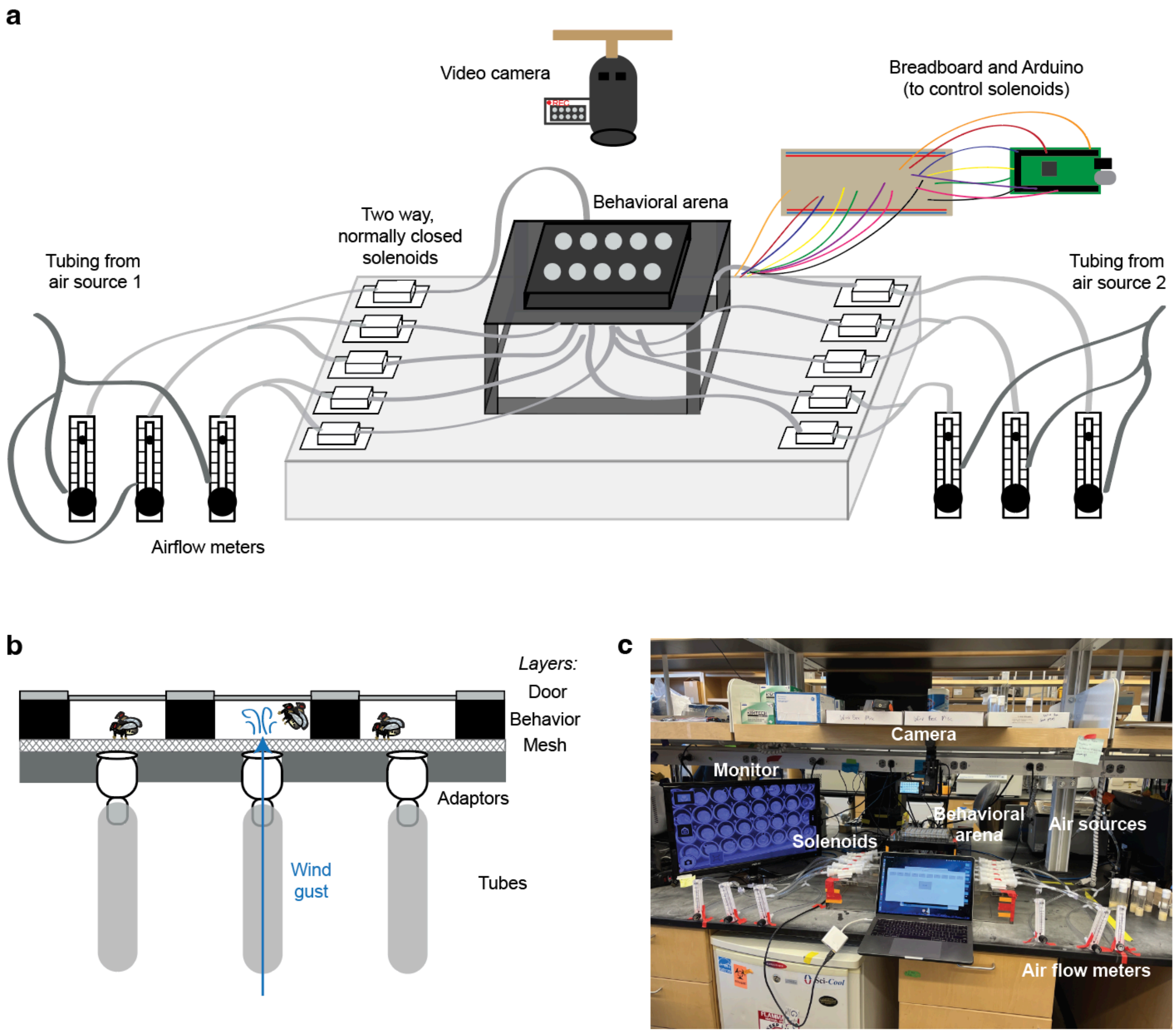


**Extended Data Figure 2: An experimental setup to deliver brief pulses of wind to mating flies**
**(a)** Flies were placed in elevated behavioral arenas connected to two-way solenoids. Compressed air sources were fed into airflow meters and then into the solenoids, which gate the delivery of the wind. By controlling solenoid opening via Arduino, specific gusts of wind of timed duration can be delivered to behavioral arenas. A camera was suspended above the behavioral arenas. For more information, see **Methods**.
**(b)** Side view of each well. Tubes were connected into wells via adapters, and a mesh layer was placed over the hole in the floor so the flies did not fall in.
**(c)** Photo of the setup. Solenoids are controlled via a computer connected to the Arduino (not in view).

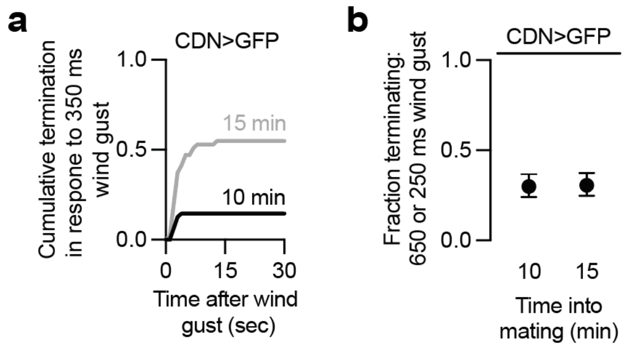


**Extended Data Figure 3: Further characterization of wind gust data**
**(a)** Flies end matings in response to a 350-millisecond wind gust with varying latencies. Cumulative distribution plot of the 10- and 15-minute data in **Figure 2d**.
**(b)** Single 650-millisecond wind gusts at 10 minutes into mating, and 250-millisecond wind gusts at 15 minutes into mating each terminate ~30% of matings. Data used to calculate independent probability of paired pulse experiment in **Figure 2f**.

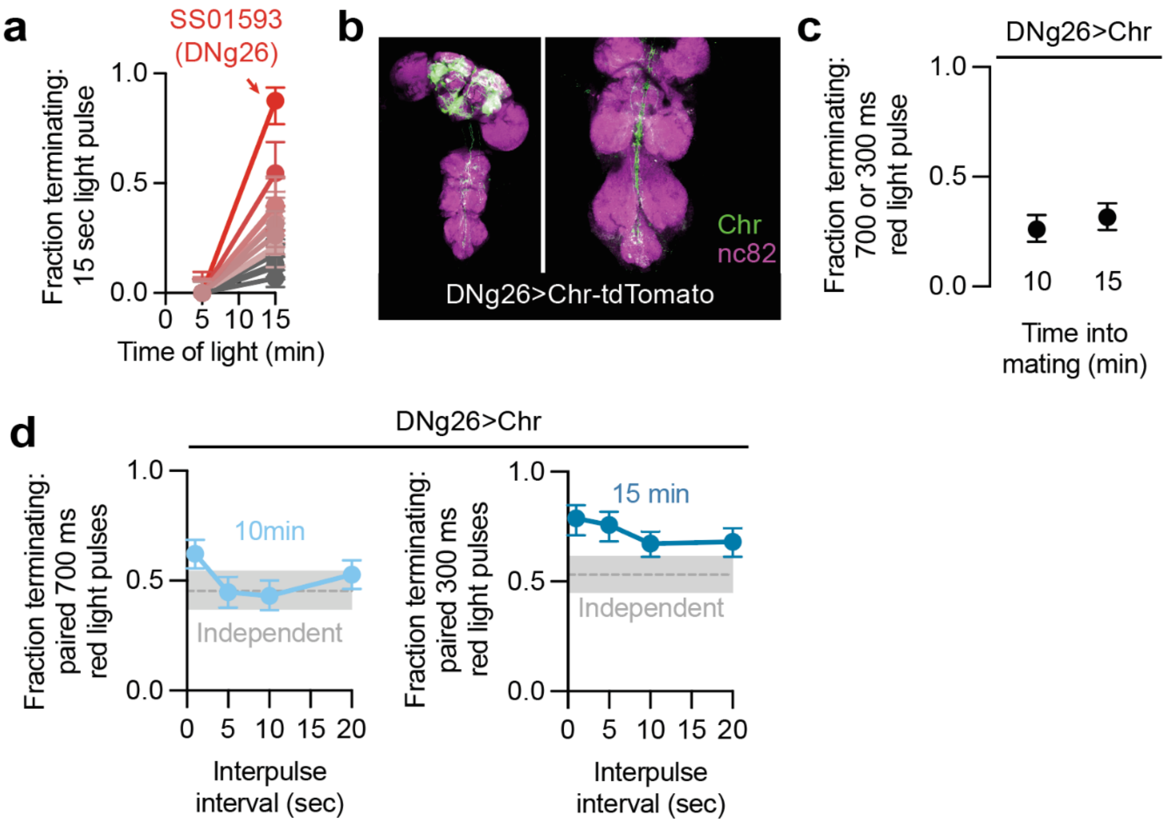


**Extended Data Figure 4: Pulses of descending neuron stimulation are integrated late in mating**

**(a)** Of 164 descending-interneuron-labeling lines screened, none terminated mating at 5 minutes when stimulated for 15 seconds with CsChrimson. SS01593 (containing the serotonergic descending neuron DNg26, as well as labeling other cells) is the most effective line at terminating mating at 15 minutes.

**(b)** DNg26 (SS01593) sends projections to the abdominal ganglion.

**(c)** 700 milliseconds of DNg26 stimulation at 10 minutes into mating, or 300 milliseconds of DNg26 stimulation at 15 minutes into mating terminate ~30% of matings.

**(d)** Paired pulse stimulation (700 ms at 10 min, 300 ms at 15 min) of DNg26 is integrated over a longer timescale when delivered at 15 minutes into mating.

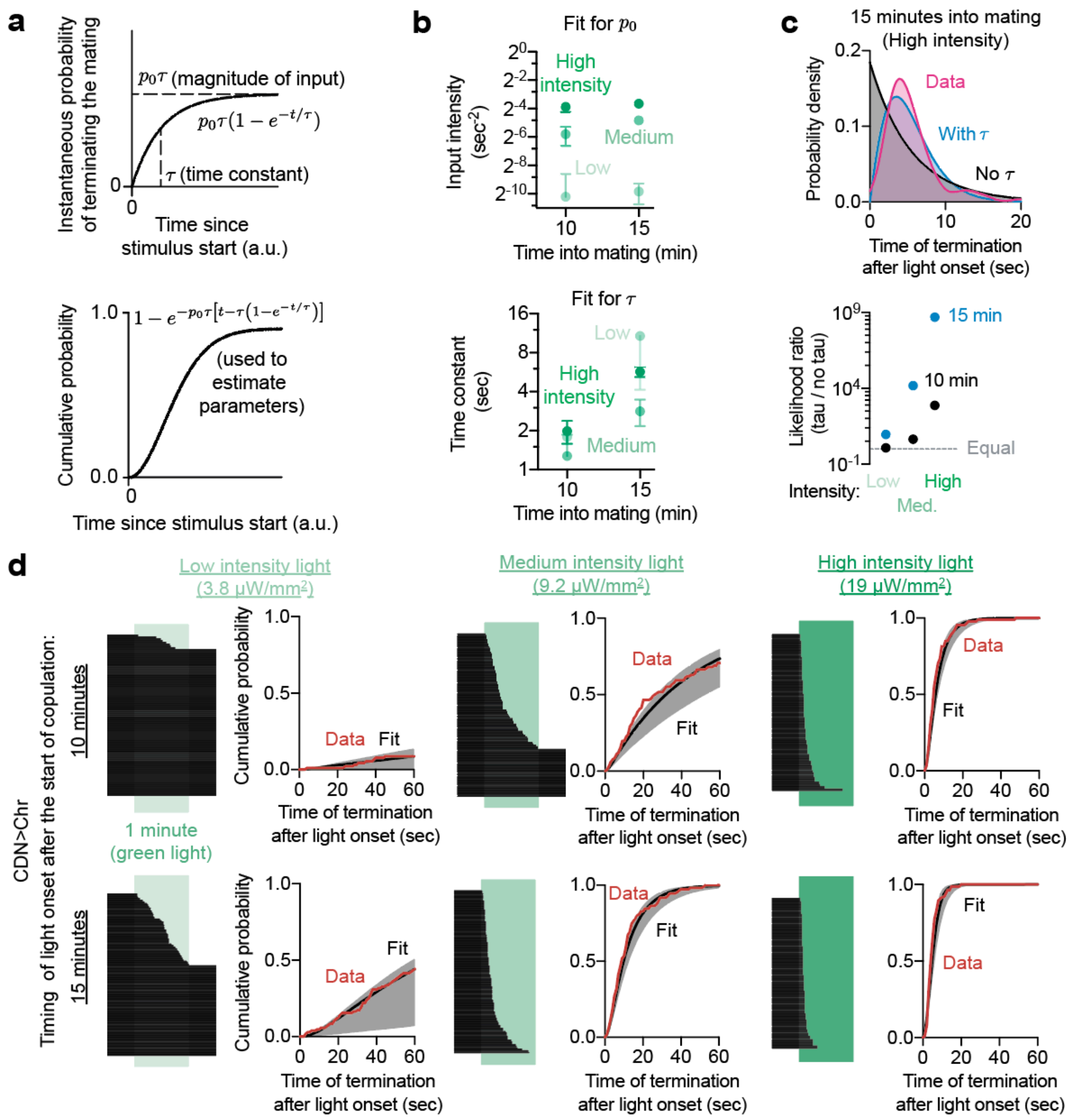


**Extended Data Figure 5: Quantitative estimation of the changing time constant of integration**

**(a)** If the instantaneous probability of terminating a mating in response to sustained stimulation ramps up as an exponential (top), then the cumulative probability of a mating ending by a particular time into a sustained stimulation follows the function $\sigma(t) = 1 - exp(-p_0\tau(t - (\tau(1 - exp(-t/\tau)))))$ (bottom).

**(b)** Parameter estimates for $\tau$ (time constant, bottom) and $p_0$ (intensity, top) across timepoints and conditions. $p_0$ is sensitive to stimulation intensity but not time into mating, while $\tau$ scales with time into mating, but not stimulation intensity. Error bars show the square root of the estimated parameter variance using the Cramér-Rao bound.

**(c)** Temporal integration is necessary to explain the behavior of flies during sustained optogenetic stimulation, as a model predicting no temporal integration (no $\tau$) ascribes a much lower likelihood to the data sets observed (bottom). Temporal integration is also needed to explain the increasing probability of termination as the stimulus goes on (top, data fit with a kernel density estimate).
**(d)** Termination times of CDN>Chr flies exposed to green light for sixty seconds (intensity indicated above graphs). Fitting the cumulative distribution to the model in **(a)** reveals a close fit. Solid black line: maximum likelihood fit. Error bars: pointwise 95% coverage intervals sampled according to estimated covariance of the parameters. The data used to fit the model for medium intensity light are the same as is plotted in **Figure 2k**.

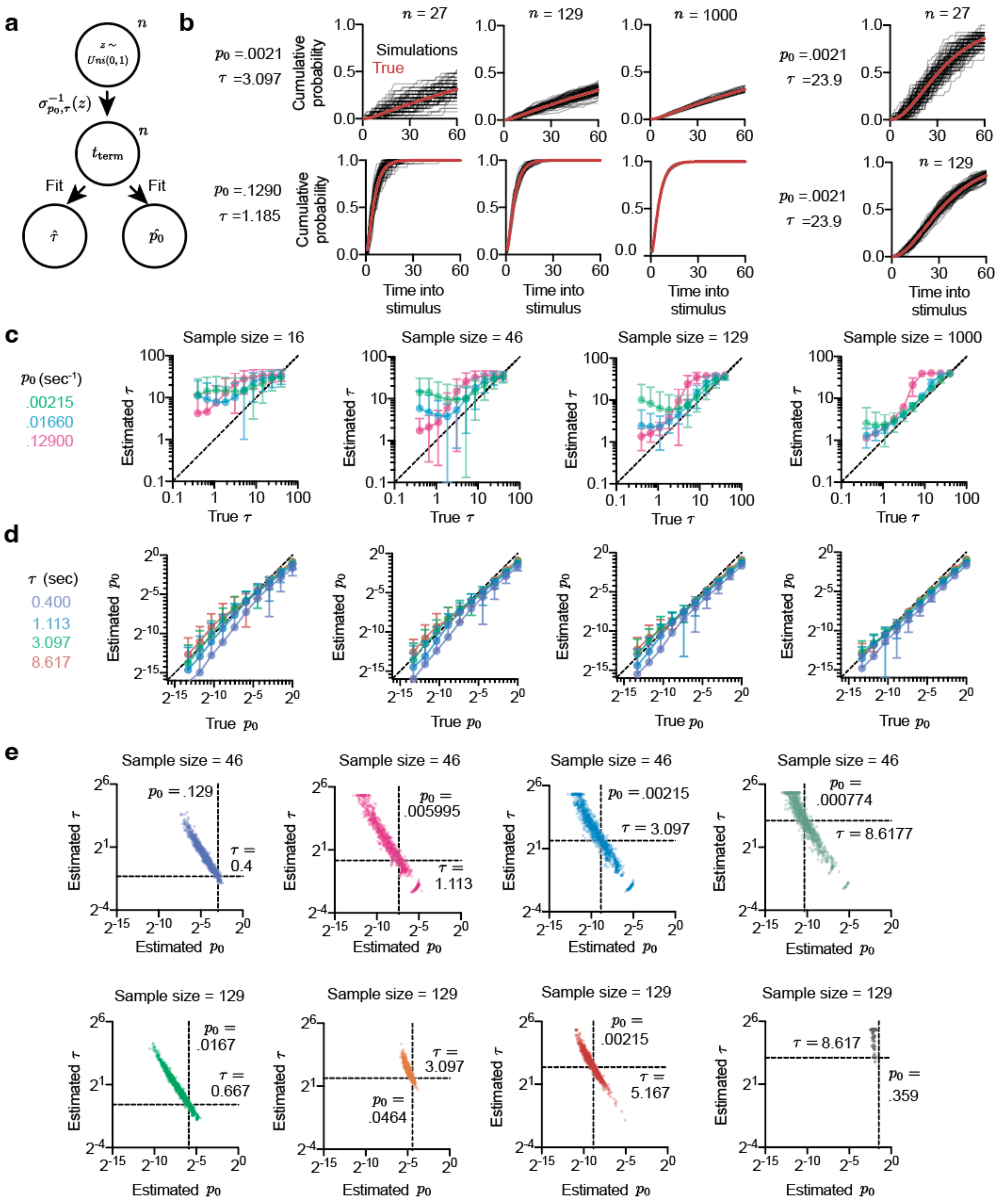


**Extended Data Figure 6: Analysis of the methodology for fitting temporal integration and sensitivity to sample size and underlying parameter values**
**(a)** Sampling scheme for generating data sets. $n$ samples were generated according to the given cumulative distribution function, $\sigma_{p_0,\tau}$, and these were used to fit estimates for the generating $\tau$

and $p_0$. The value of $n$ was varied logarithmically from 10 to 1000 to evaluate what sample size would be necessary to accurately estimate the parameters of the distribution.
**(b)** The cumulative distribution function can be qualitatively reconstructed with samples of size ~ 100 across a wide range of cumulative distribution function shapes. Smaller sample sizes (e.g. ~30) are highly variable, especially when the overall number of flies terminating the mating during the stimulation is low (top row).
**(c)** The sensitivity of the inference of the value of $\tau$ to sample size across a range of $p_0$ values. The closer a point is to the diagonal, the more likely the fitting procedure is to capture the correct $\tau$. The fitting procedure overestimates $\tau$ at low sample sizes, especially when the true value for $\tau$ is small. This may, to some extent, be explained by the fact that termination times are rounded to the nearest second (we find it is impossible to judge the time of termination more precisely than this value, given the complex motor sequence of terminating the mating). For larger sample sizes, the estimate is much better, so long as a large number of flies terminate the mating during the stimulation. When $p_0$ and $\tau$ are both small, however, the inference is considerably less reliable, because these conditions correspond to cases in which very few flies terminate the mating during the stimulus, providing very little information about $\tau$.
**(d)** As in ©, but instead examining the sensitivity of the estimate of $p_0$. The parameter $p_0$ is easier to estimate, because even flies that do not terminate the mating during the stimulation are still informative about its value to some extent (see **Methods**). However, we find that $p_0$ is systematically underestimated due to the bias towards overestimating $\tau$ and the fact that the two estimates show substantial anticovariance (elaborated in **Extended Data Figure 6e**).
**(e)** Covariance of $p_0$ and $\tau$ for various sample sizes. Dashed lines indicate the true parameter values, while independent points show individual sample estimates. The two parameters always anticovary, as indicated by the diagonal slant of each distribution. This reflects the fact that $p_0$ only appears in the cumulative distribution with $\tau$ in the form $p_0\tau$, and so this term is easier to fit than either value alone. If $\tau$ is overestimated, $p_0$ will tend to be underestimated to compensate. The multiplicative relationship is clear from the approximately linear covariance of the logarithm of the two parameters. When the data is more informative about $\tau$, i.e. many flies terminate the mating during the experiment, the cluster is much smaller (e.g. the orange data set). We therefore restricted our experiments to those conditions that would generate reliable estimates of the parameters, especially in cases where we expected $\tau$ or $p_0$ to be very small.

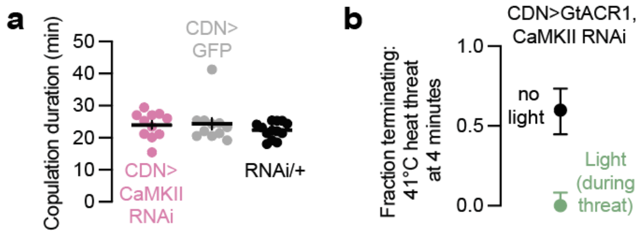


**Extended Data Figure 7: Further characterization of CaMKII knockdown in the CDNs**
**(a)** CaMKII knockdown in the CDNs does not alter mating duration.
**(b)** Mating termination is still dependent on CDN electrical activity when CaMKII is knocked down in the CDNs.

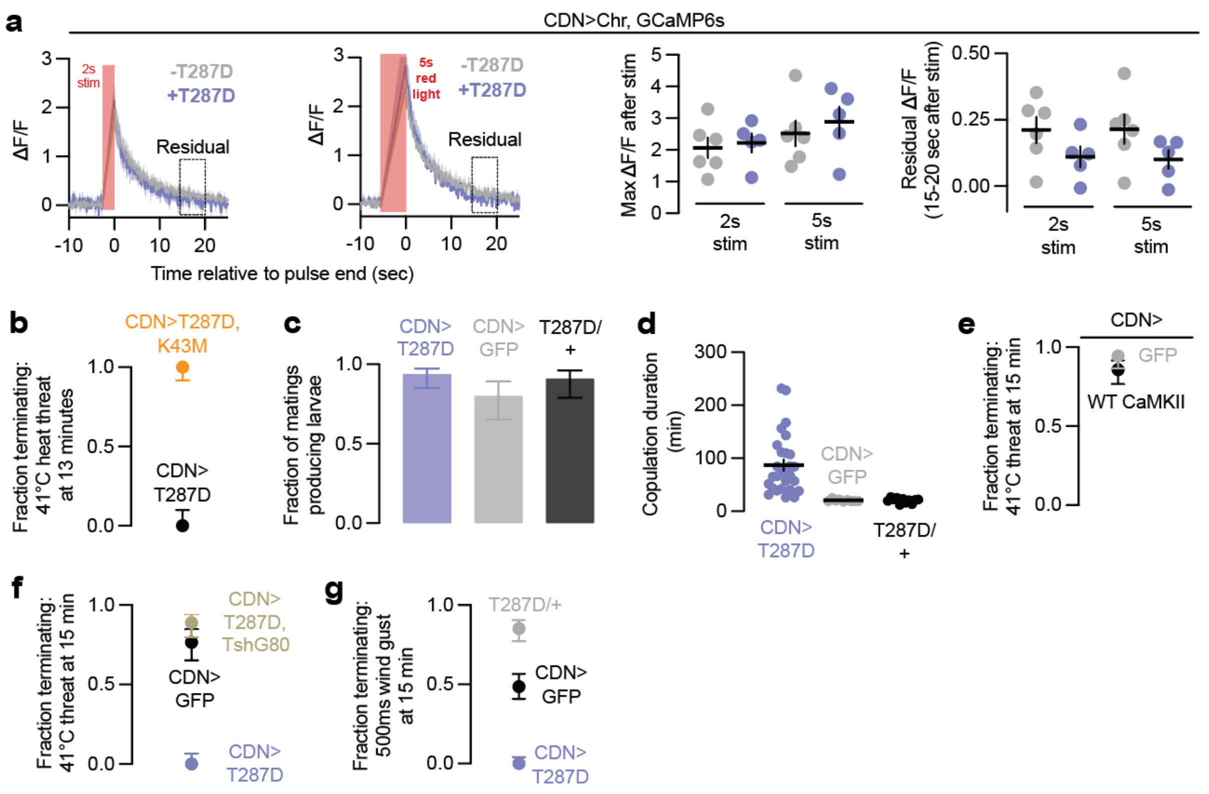


**Extended Data Figure 8: The catalytic activity of CaMKII specifically suppresses CDN response to demotivating inputs**

**(a)** Constitutively active CaMKII (T287D) has no obvious effect on the ability of CsChrimson stimulation to evoke calcium transients in the CDNs, as measured by changes in fluorescence of GCaMP6s. Left, middle left: average traces after 2 and 5 seconds of CsChrimson stimulation. Middle right: max fluorescence after stimulation. Right: average residual calcium 15-20 seconds after stimulation.

**(b)** CaMKII T287D in the CDNs cannot suppress the heat threat response with a non-functional catalytic domain (K43M). Both the T287D and K43M mutations are contained on the same UAS-CaMKII transgene.

**(c)** CaMKII T287D in the CDNs has no effect on fertility.

**(d)** CaMKII T287D in the CDNs extends mating duration.

**(e)** Overexpression of wildtype CaMKII in the CDNs does not decrease the likelihood of terminating mating in response to a heat threat.

**(f)** Subtracting the ventral nervous system expression from our CDN driver expressing CaMKII T287D prevents its effects on mating motivation.

**(g)** Expressing constitutively active CaMKII in the CDNs prevents termination in response to wind.

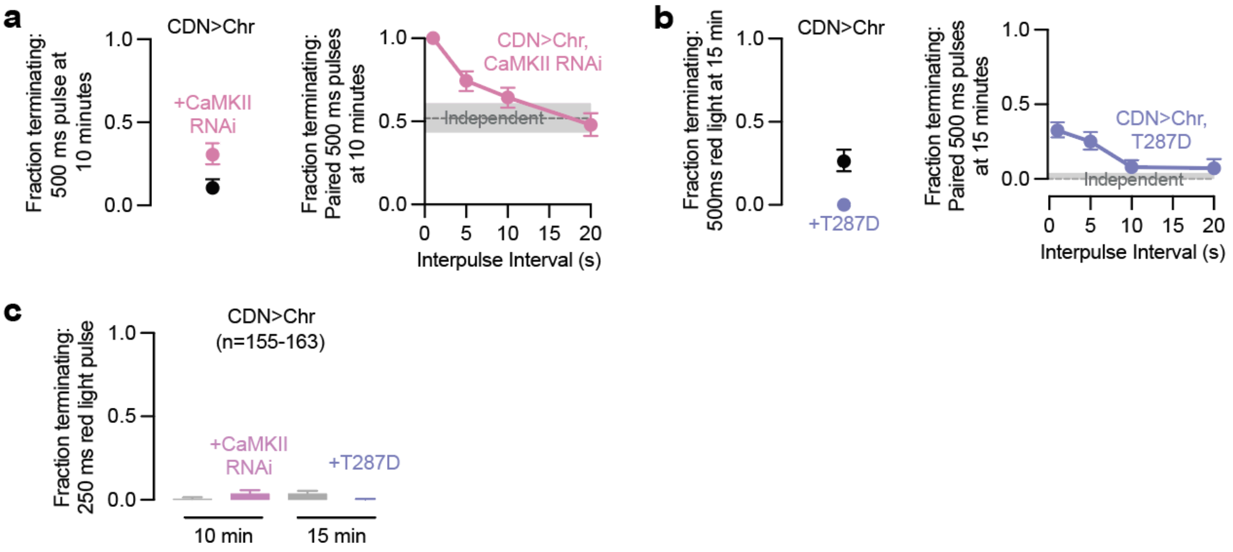


**Extended Data Figure 9: CaMKII manipulations alter the response to 500ms pulses of CDN stimulation**

**(a)** Left: termination response to 500ms red light stimulation of the CDNs with CaMKII knockdown. Right: paired pulse response with CaMKII knockdown in the CDNs.

**(b)** Left: termination response to 500ms red light stimulation of the CDNs with expression of CaMKII T287D. Right: paired pulse response with expression of CaMKII T287D in the CDNs.

**(c)** A single 250 millisecond pulse of CDN stimulation is very unlikely to cause termination regardless of time into mating or CaMKII manipulation. Plotted is the fraction of flies (for cycle lengths: 2.5, 5, and 10 sec) from **Figure 3g** that terminated in response to the first of 10 pulses.

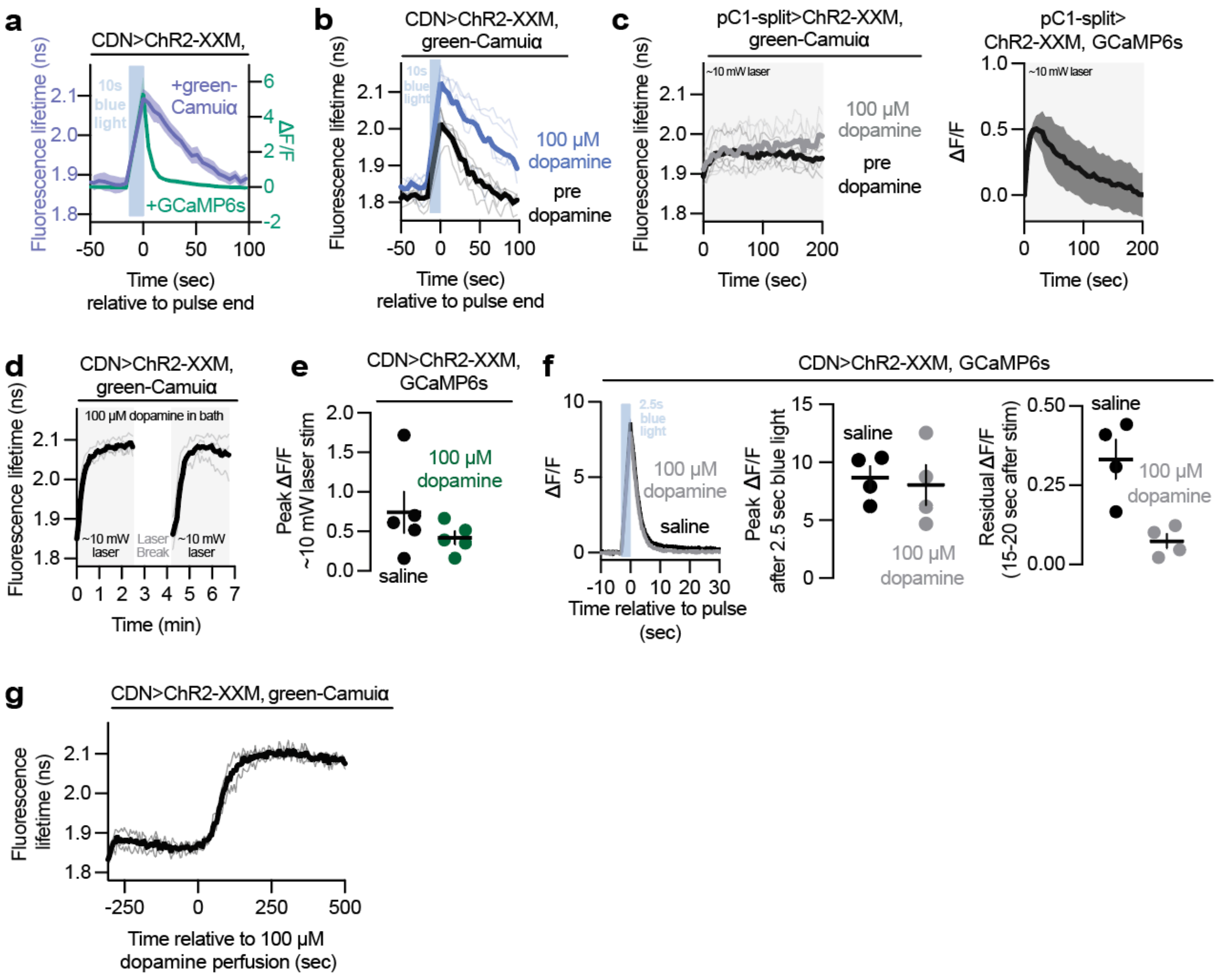


**Extended Data Figure 10: Dopamine potentiates CaMKII activity in the CDNs without increasing calcium influx**

**(a)** CaMKII activity in the axons of the CDNs (as reported by the fluorescence lifetime of the FRET sensor green-Camuiα) decays over ~1 minute after 10 seconds of blue light stimulation of the Channelrhodpsin-2 variant ChR2-XXM whereas calcium levels (as measured by changes in the fluorescence of GCaMP6s) decline over ~5 seconds. [31][12]Laser power was kept at ~5 milliwatts, to limit the basal excitation of ChR2-XXM. Error bar shading for all imaging data represents SEM.
**(b)** CaMKII activity, as reported by change in fluorescence lifetime of green-Camuiα (left), is more strongly activated by transient blue light stimulation of ChR2-XXM in the presence of dopamine perfusion (right). Laser power was kept at ~5 mW, to limit the basal excitation of ChR2-XXM.
**(c)** Left: Dopamine does not potentiate CaMKII activity in another set of Dsx+ neurons (female pC1 neurons[25]). Right: ~10 mW laser stimulation of ChR2-XXM in pC1 neurons induces an increase in GCaMP6s fluorescence that relaxes to baseline after a few minutes.
**(d)** Constant stimulation is required to keep CaMKII activity high in the presence of dopamine. After stopping laser stimulation for ~100 seconds (“laser break”), CaMKII levels return to baseline and then ramp up again once the laser is turned back on.
**(e - f)** Dopamine does not increase calcium influx in the CDNs. (e) Peak calcium levels (from **Figure 4e**) in response to ~10 mW laser stimulation of ChR2-XXM before and after dopamine perfusion. (f) Left: average traces of 2.5 seconds blue light stimulation of ChR2-XXM with saline

and dopamine perfusion. Middle: peak calcium after stimulation. Right: residual calcium 15-20 seconds after stimulation (note the small y-axis to highlight a potential effect on residual calcium). **(g)** Dopamine potentiates CaMKII activation under continuous ~10 mw infrared laser ChR2-XXM stimulation.

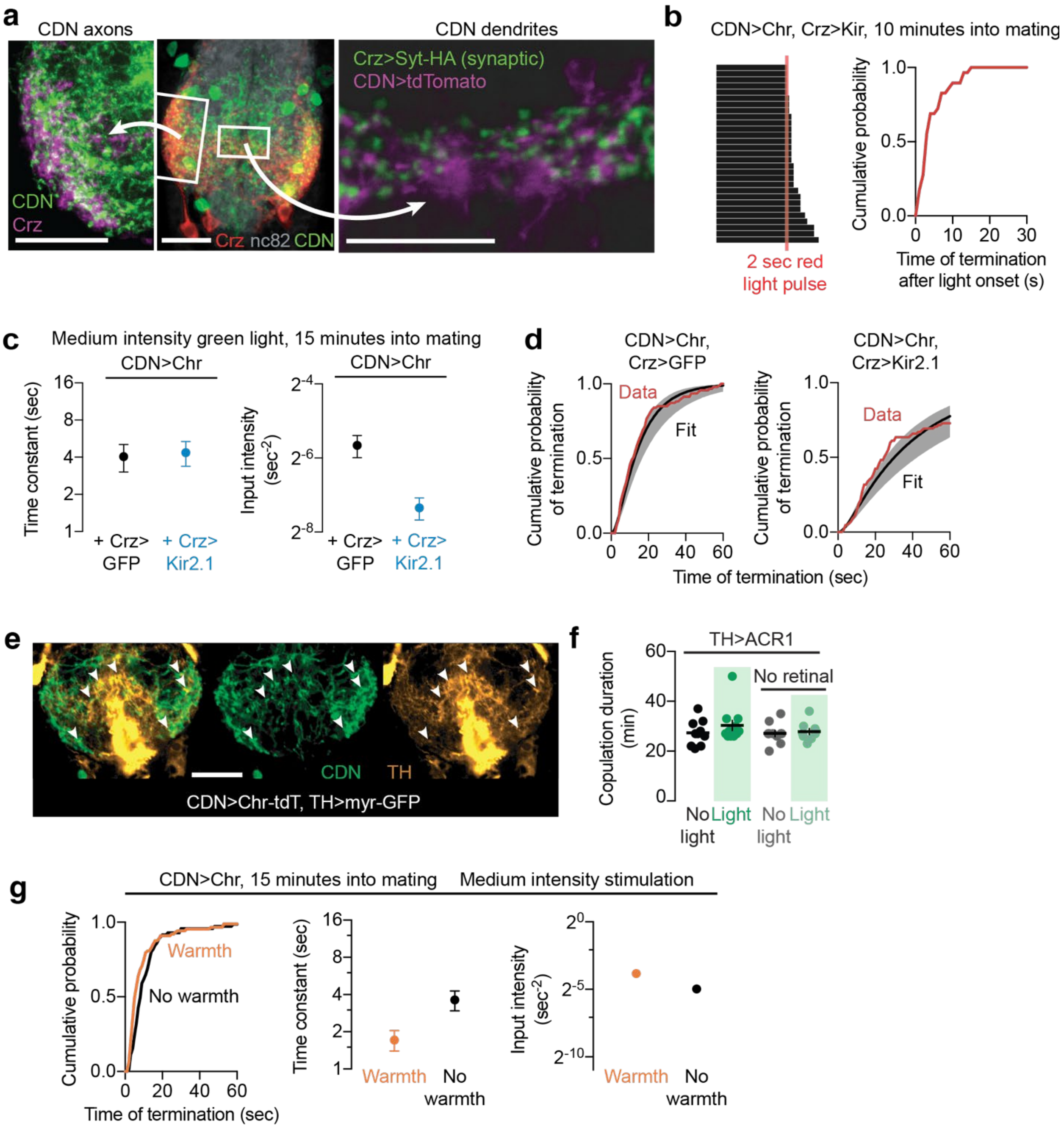


**Extended Data Figure 11: The interactions of dopaminergic and Corazonin neurons with the CDNs**

**(a)** The Crz neurons project throughout the abdominal ganglion, with processes closely apposed to those of the CDNs, both near their axons (left) and dendrites (right), though synaptic connectivity cannot be concluded.

**(b)** Optogenetic stimulation of the CDNs while the Crz neurons are silenced results in termination of the mating, demonstrating that the CDNs operate downstream of the Crz neurons in determining the motivational state of the fly.

**(c)** Silencing the Crz neurons reduces the response to sustained stimulation of the CDNs by selectively decreasing the gain on the input (~8-fold), leaving the time constant of integration largely unaffected.
**(d)** Cumulative distribution functions used for estimating the parameters of panel c
**(e)** Dopaminergic neurons (orange) send projections throughout the abdominal ganglion, often forming varicosities near CDN (green) processes (indicated by white arrowheads). Images are obtained from a single optical plane.
**(f)** Silencing the dopaminergic neurons does not affect overall copulation duration.
**(g)** Warmth alone decreases $\tau$ but increases $p_{!}$, showing that heat cannot account for the effects of stimulation the dopaminergic neurons.

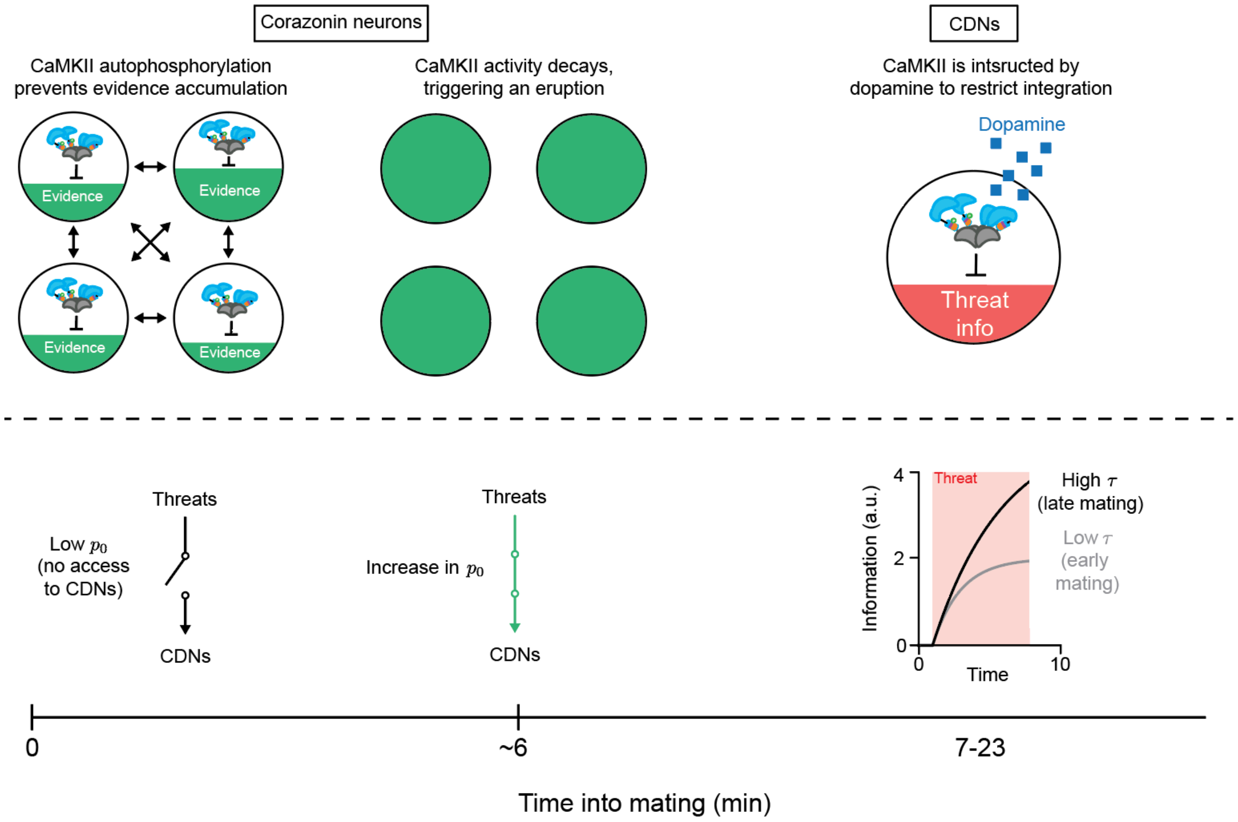


**Extended Data Figure 12: The circuitry controlling motivational dynamics during mating.** For the first ~6 minutes of mating, high CaMKII activity in the Corazonin neurons prevents the network eruption that triggers sperm transfer[13,14]. Before the eruption, males that have not mated recently are impervious to challenges of apparently all varieties and severities. At 6 minutes, the eruption increases p0, allowing threat information to be delivered to the Copulation Decision Neurons (CDNs), through mechanisms not yet understood. After the eruption, dopaminergic inputs to the CDNs, increase intracellular CaMKII levels to restrict tau, the timescale of competing information retention.

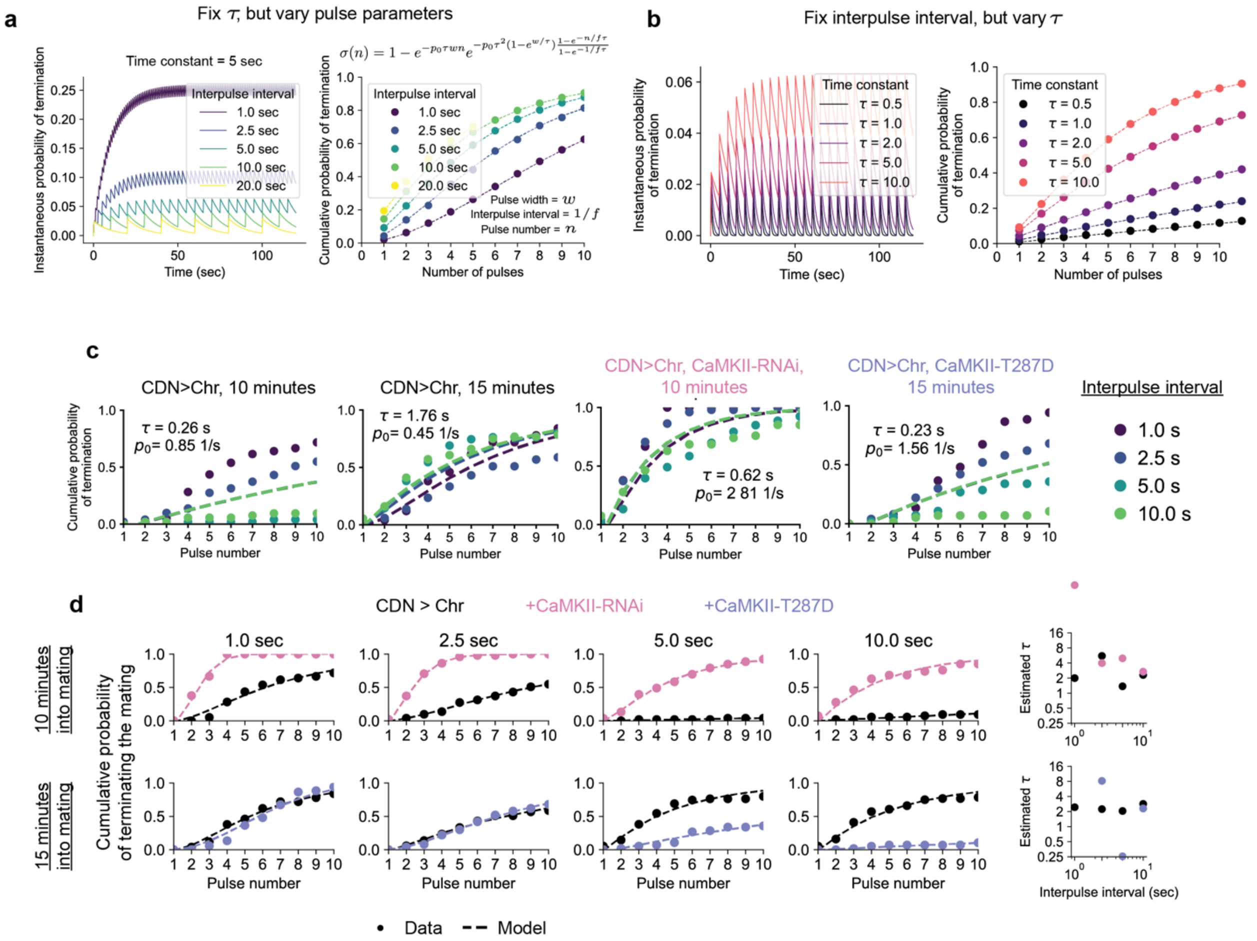


**Extended Data Figure 13: Linear systems analysis of the multi-pulse experiments**

**(a)** Left: The instantaneous probability of terminating the mating increases with successive pulses when those pulses are spaced out by a time shorter than $\tau$. Right: When the data are quantified by which pulse occurred most recently before the mating ends, the apparent effect of the first several pulses is higher when the pulses are spaced farther apart in time, even though the instantaneous termination probability is lower, because there is only a short opportunity to terminate in response to any given pulse before the next arises. This confounds the interpretation of termination probability in the multi-pulse experiments, because an increased response for a given interpulse interval does not necessarily reflect better temporal integration (this simulation fixes $\tau$ but shows a substantial difference in termination probability by pulse, with a seemingly-paradoxical *smaller* response probability when the pulses are spaced apart by less than $\tau$.

**(b)** However, for any given interpulse interval, the termination probability per pulse increases monotonically with $\tau$, and so differences like those observed in **Figure 3G** are still consistent with increasing time constants of integration.

**(c)** Analyzing the data from **Figure 3G** and attempting to fit a single pair of parameters for each genotype and time point results in curves which do not fit any data set particularly well, but make predictions consistent with our interpretations: loss of CaMKII function increases $\tau$, as does testing at 15 minutes as compared to 10 minutes, while constitutively active CaMKII suppresses $\tau$.

**(d)** Fitting each experiment independently shows that the data are not consistent with a single estimate of $\tau$ or $p_0$, illustrating the numerical instability and inadequacy of the first order linear model for this experiment.

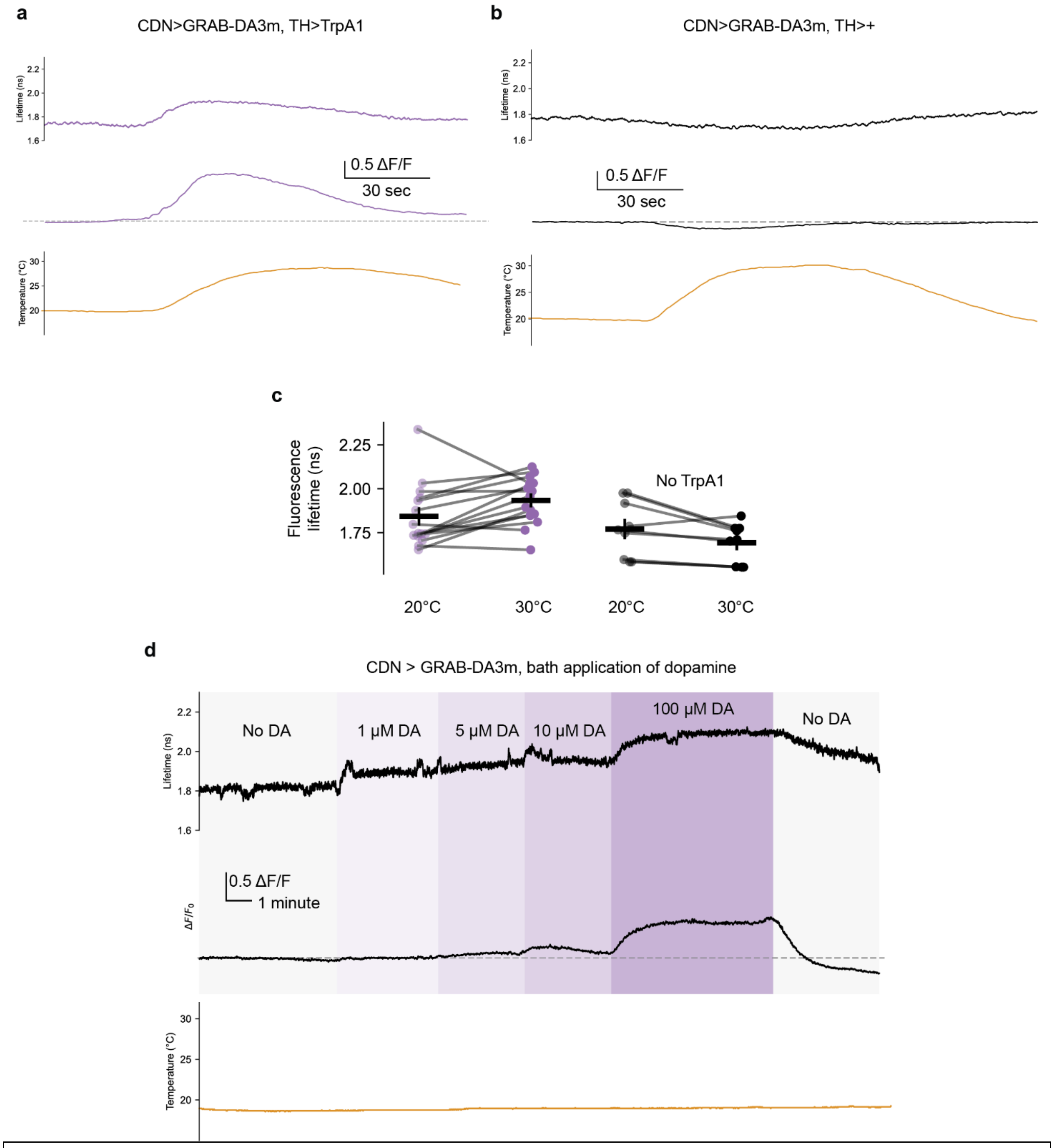


**Extended Data Figure 14: Further analysis of GRAB-DA3m experiments**
**(a)** Thermogenetic stimulation of the dopaminergic neurons of the abdominal ganglion results in an increase in fluorescence and fluorescence lifetime. Note that 1) the lifetime is not linearly related to the increase in fluorescence (and so the two measures have differential sensitivity across concentration) and 2) both signals begin to decrease before the temperature is decreased. Because bath application of dopamine resulted in stable fluorescence (see **Extended Data Figure 14**), this seems unlikely to be bleaching of the indicator. We speculate it results either from habituation of the TrpA1 channel or rapid depletion of the dopaminergic

neurons, at least at the scale of the sensor's dynamic range. Allowing several minutes of recovery at 20°C permitted a second stimulation to be equally efficacious (not shown).

**(b)** Warming the abdominal ganglion without expression of TrpA1 in dopaminergic neurons resulted in a small but consistent decrease in both fluorescence lifetime and fluorescence itself. The GRAB-DA3m protein itself is likely temperature sensitive, but the effect of temperature produces a change in fluorescence signal opposite to that observed during stimulation of dopaminergic neurons, arguing that signals as in **Figure 4b** are not artifacts of the temperature ramp.

**(c)** Data as in **Figure 4b** but plotting lifetime instead of fluorescence. The measurement was more variable but there is a consistent ~200 picosecond increase in lifetime with thermogenetic stimulation.

**(d)** Bath application of dopamine in the concentration range used in **Figure 4** results in increases in fluorescence lifetime and fluorescence quantitatively similar to that evoked by thermogenetic stimulation of dopaminergic neurons, arguing that these bath concentrations result in physiologically-plausible exposure to dopamine at the CDN axons. These values differ substantially from the reported sensitivity of GRAB-DA3m[30] which we speculate arises from the protective glial sheath surrounding the abdominal ganglion, which may buffer the exogenous dopamine levels or rapidly degrade it. Supporting this conclusion, in unpublished experiments, we found that single application of dopamine to a still bath (rather than continuous perfusion) only transiently increased the excitability of CaMKII, unlike the sustained excitability increase observed in **Figure 4**.

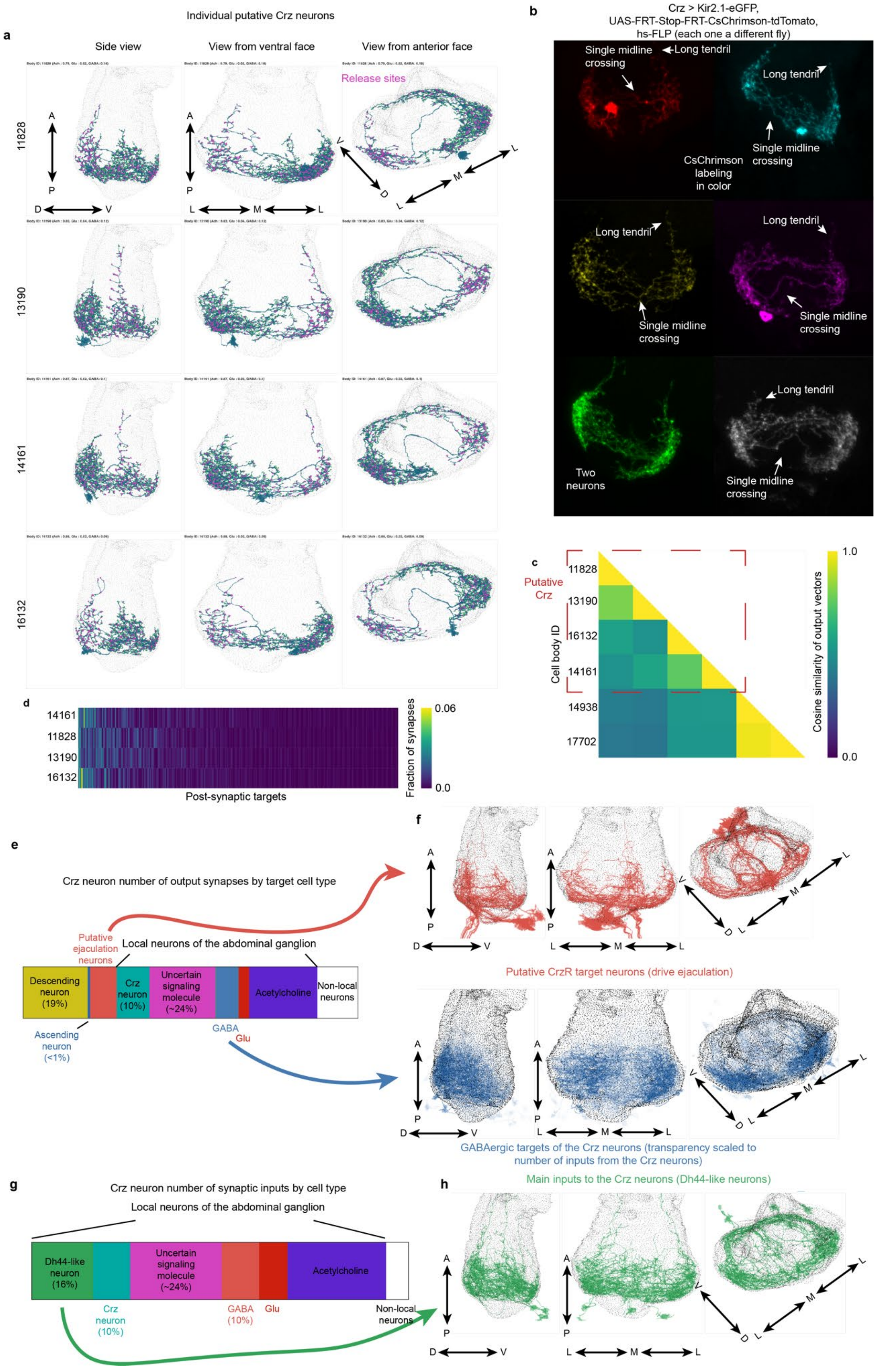

Individual putative Crz neurons
a
Side view
View from ventral face
View from anterior face
Release sites
11828
13190
14161
16132
A
P
D
V
L
M
b
Crz > Kir2.1-eGFP,
UAS-FRT-Stop-FRT-CsChrimson-tdTomato,
hs-FLP (each one a different fly)
Single midline crossing
Long tendril
CsChrimson labeling in color
Two neurons
c
Putative Crz
Cell body ID
11828
13190
16132
14161
14938
17702
Cosine similarity of output vectors
1.0
0.0
d
Post-synaptic targets
Fraction of synapses
0.06
0.0
e
Crz neuron number of output synapses by target cell type
Putative ejaculation neurons
Local neurons of the abdominal ganglion
Descending neuron (19%)
Crz neuron (10%)
Uncertain signaling molecule (~24%)
Acetylcholine
Non-local neurons
Ascending neuron (<1%)
GABA
Glu
f
Putative CrzR target neurons (drive ejaculation)
GABAergic targets of the Crz neurons (transparency scaled to number of inputs from the Crz neurons)
g
Crz neuron number of synaptic inputs by cell type
Local neurons of the abdominal ganglion
Dh44-like neuron (16%)
Uncertain signaling molecule (~24%)
Acetylcholine
Crz neuron (10%)
GABA (10%)
Glu
Non-local neurons
h
Main inputs to the Crz neurons (Dh44-like neurons)

**Extended Data Figure 15: EM analysis of the Corazonin neurons**

**(a)** Four neurons in the MANC volume closely resembling the morphology of the Crz neurons. Individual release sites (localized to both sides of the AG) shown in pink. Three perspectives are shown: from the right side of the fly (left column), from the ventral side of the fly (central column), and from the anterior side (right column).

**(b)** Single cell labeling of individual Crz neurons using a heat-shock induced recombinase. Data from Thornquist *et al.*, 2021[14].

**(c)** Cosine similarity of the synaptic output vectors of six Crz-like neurons (body IDs shown) suggests four neurons are approximately equally similar to one another and a separate pair that is more similar internally than to the other two.

**(d)** Most postsynaptic targets of the four Crz-like neurons identified are shared.

**(e)** Most of the Crz neurons' postsynaptic targets are local neurons of the abdominal ganglion. ~30% of their output synapses are onto descending cells, likely to drive ejaculation and the accompanying abdominal movements. 10% of their synapses are reciprocal (innervating one another), and ~50% of their outputs are onto many different cell classes in the AG.

**(f)** Top: One of the primary classes of neurons targeted by the putative Crz neurons is a set of enervating neurons projecting down the abdominal trunk nerve. These neurons resemble the ejaculation-driving Tph2/CrzR expressing cells from Tayler *et al*. Bottom: The GABAergic neurons targeted by the Crz neurons are largely restricted to the posterio-lateral portion of the AG, but predominantly do not appear CDN-like.

**(g)** A large fraction of the inputs the Crz neurons receive come from either other Crz neurons, or four other local neurons of the AG.

**(h)** The primary source of synaptic input to the Crz neurons are four peptidergic cells in the AG resembling the neurons labeled by Dh44-Gal4 (unpublished data).

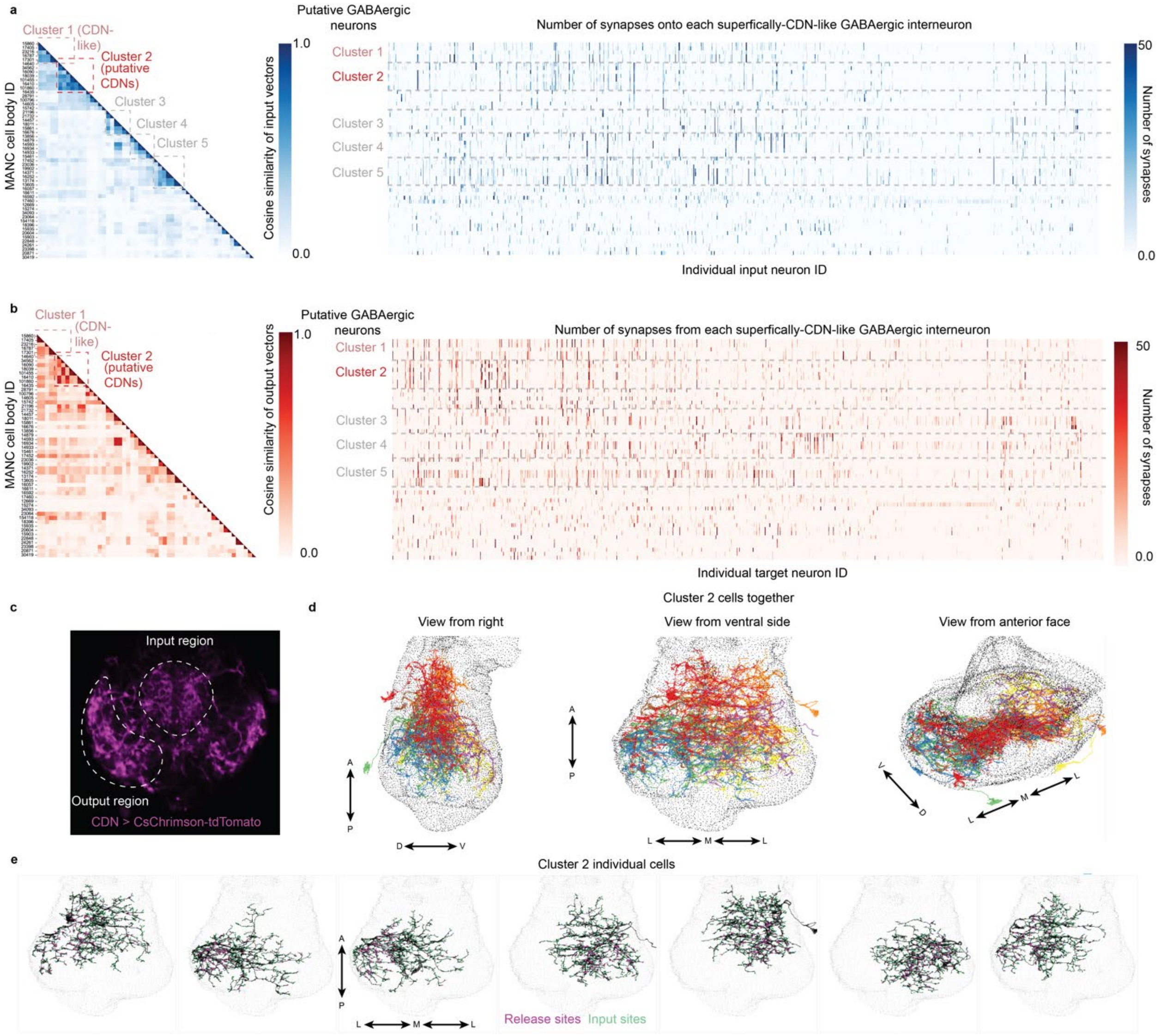


**Extended Data Figure 16: Identification of putative CDNs**

**(a)** The 54 predicted-GABAergic neurons of the abdominal ganglion that superficially resemble the CDNs cluster into five groups, as well as a miscellaneous collection, by comparing the proportions and identities of their various synaptic inputs. Left: the cosine similarity of the vector of synaptic inputs for each cell. Right: Each putative CDN's synaptic inputs across the collection of cells innervating any of the 54 possible candidates.

**(b)** As in **a**, but for the synaptic output vectors of the CDN candidates

**(c)** IHC of central planes of the abdominal ganglion highlights two features of the CDNs: synaptic release sites along the dorsolateral AG and inputs in the central AG.

**(d)** The overlaid anatomy of the 7 neurons identified in Cluster 2, presumed to be the CDNs, from three perspectives. Left: viewed from the fly's right side. Middle: viewed from the ventral side of the fly, Right: viewed from the anterior side of the fly.

**(e)** The anatomy of each individual CDN is varied, innervating different portions of the antero-posterior extent of the later abdominal ganglion

**(d)** Each individual cell in Clusters 1 and 2 of **Extended Data Figure** 16 receives relatively little reciprocal innervation from its postsynaptic targets, especially as compared to the other GABAergic neuron classes.
**(e)** Pooling the CDNs suggests an increase in reciprocal innervation across the cell class, suggesting that many post-synaptic targets of each putative CDN innervates other CDNs. The diagonal of these plots is still much less dense in Clusters 1 and 2, implying that these cells are less recurrently connected than other morphologically-similar interneurons of the AG.

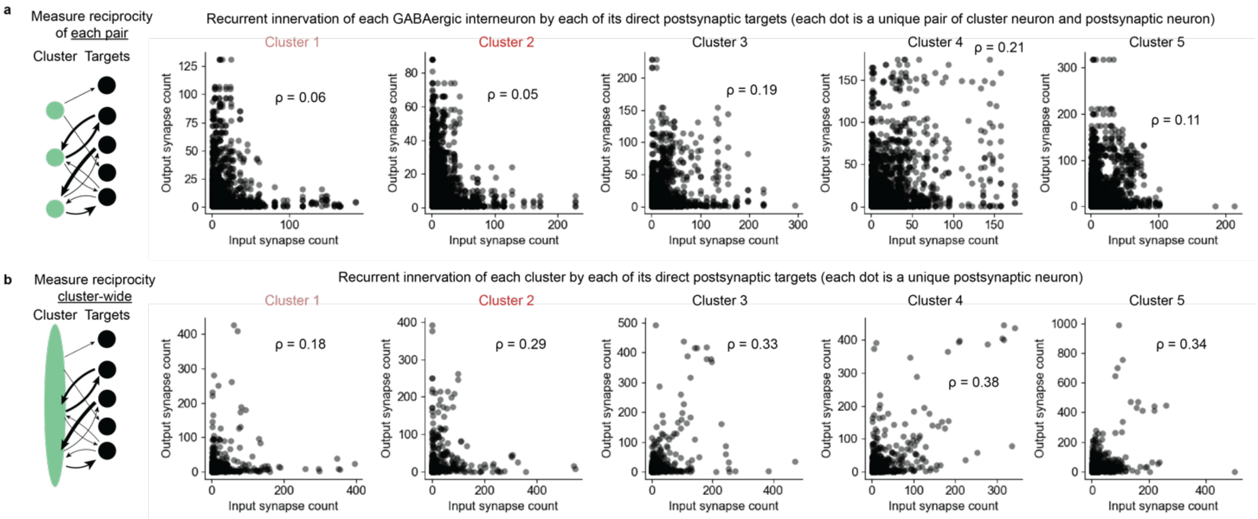


**Extended Data Figure 17: Analysis of recurrence between putative CDNs and their postsynaptic targets**

**(a)** Each individual cell in Clusters 1 and 2 of **Extended Data Figure** 16 receives relatively little reciprocal innervation from its postsynaptic targets, especially as compared to the other GABAergic neuron classes.

**(b)** Pooling the CDNs suggests an increase in reciprocal innervation across the cell class, suggesting that many post-synaptic targets of each putative CDN innervates other CDNs. The diagonal of these plots is still much less dense in Clusters 1 and 2, implying that these cells are less recurrently connected than other morphologically-similar interneurons of the AG.

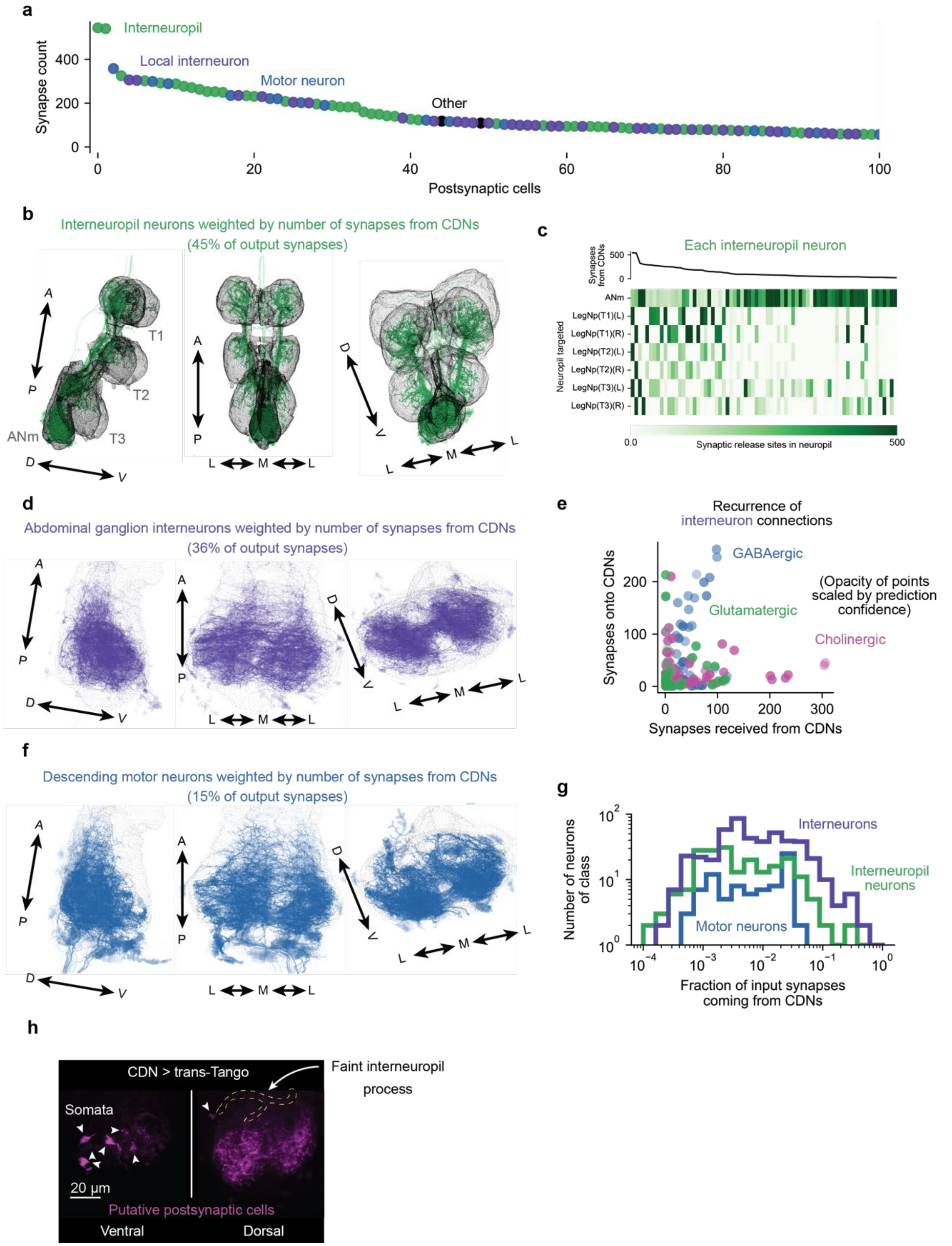


**Extended Data Figure 18: Putative CDNs principally target abdominal ganglion interneurons, and receive few direct recurrent inputs**

**(a)** The 100 cells receiving the most input from the presumed CDNs make up over 80% of all output synapses, and can be divided into three classes: interneuropil neurons of the VNS, local

interneurons of the abdominal ganglion, and motor neurons descending the abdominal trunk nerve.
**(b)** The interneuropil targets of the CDNs innervate all three leg neuropil.
**(c)** Most interneuropil targets of the CDNs that receive a substantial amount of CDN input do not innervate the abdominal ganglion, and instead transmit information to all of the leg neuropil (rather than innervating a single neuropil.
**(d)** The abdominal ganglion interneurons targeted by the CDNs densely innervate both halves of the AG with very little input or output in the medial third of the ganglion.
**(e)** The local interneurons of the AG do not strongly reciprocate synaptic input from the CDNs, though a subset of neurons weakly predicted to be GABAergic do synapse back onto the CDNs, providing the opportunity for recurrence through disinhibition.
**(f)** The minority descending neuron output of the CDNs.
**(g)** Motor neurons targeted by the CDNs receive only a very small fraction of their inputs from the CDNs. Most interneuropil neurons receive <10% of their synaptic input from the CDNs. In contrast, many AG interneurons receive a large fraction of their input from the CDNs.
**(h)** Trans-synaptic labeling of CDN targets identifies six large AG interneurons, matching the number of major interneuron targets in the EM volume, but mostly fails to resolve motor or interneuropil neurons.

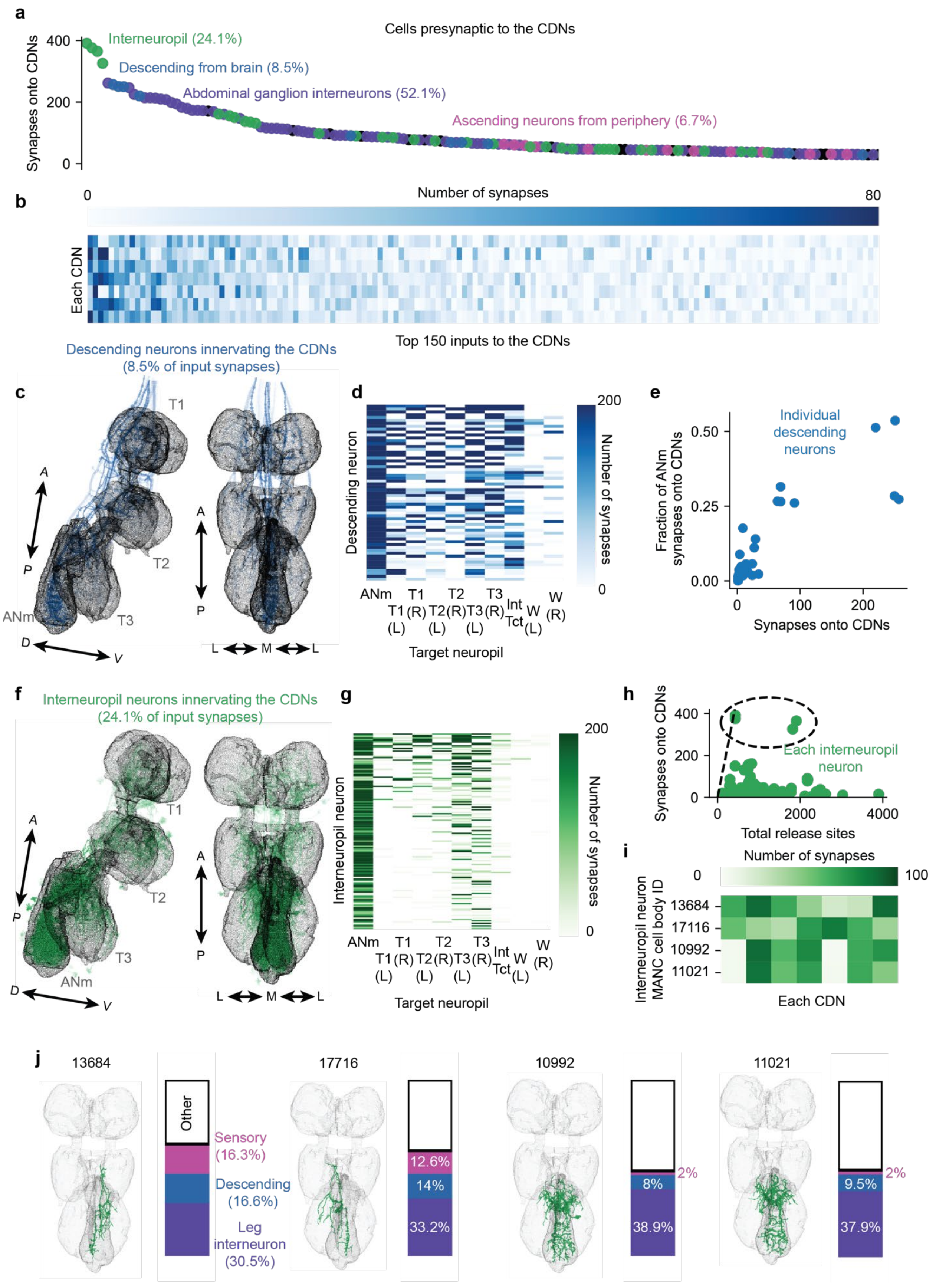
a
Cells presynaptic to the CDNs
Synapses onto CDNs
Interneuropil (24.1%)
Descending from brain (8.5%)
Abdominal ganglion interneurons (52.1%)
Ascending neurons from periphery (6.7%)
b
Number of synapses
Each CDN
Top 150 inputs to the CDNs
c
Descending neurons innervating the CDNs
(8.5% of input synapses)
d
Descending neuron
Number of synapses
Target neuropil
e
Individual descending neurons
Fraction of ANm synapses onto CDNs
Synapses onto CDNs
f
Interneuropil neurons innervating the CDNs
(24.1% of input synapses)
g
Interneuropil neuron
Number of synapses
Target neuropil
h
Synapses onto CDNs
Each interneuropil neuron
Total release sites
i
Number of synapses
Interneuropil neuron MANC cell body ID
13684
17116
10992
11021
Each CDN
j
13684
Other
Sensory (16.3%)
Descending (16.6%)
Leg interneuron (30.5%)
17716
12.6%
14%
33.2%
10992
2%
8%
38.9%
11021
2%
9.5%
37.9%

**Extended Data Figure 19: Putative CDNs receive integrative inputs from the brain and other VNS neuropil**

**(a)** The 150 primary inputs to the CDNs are divided into four classes: interneuropil neurons of the VNS, local interneurons of the abdominal ganglion, descending neurons from the brain, and ascending sensory neurons from the periphery.

**(b)** Most neurons that target the CDNs target them indiscriminately, with little preferential innervation of individual cells, arguing that the functional unit of the CDNs is the collective, and that their individual differences are not a primary feature.

**(c,d)** Descending neurons from the brain targeting the CDNs typically innervate other neuropil as well.

**(e)** A few descending neurons do preferentially target the CDNs, with ¼ to ½ of their synapses being specific to the CDNs, but other direct descending input to the CDNs is minimal.

**(f,g)** A quarter of input synapses to the CDNs come from interneuropil neurons of the VNS that receive input from multiple other leg neuropil

**(h)** Four interneuropil neurons heavily innervate the CDNs, making up their four largest inputs, and two of these almost exclusively synapse on the CDNs (dotted black line is the unity line, indicating all synapses being restricted to the CDNs)

**(i)** The four interneuropil neurons targeting the CDNs target them mostly indiscriminately, rather than singling out individual CDNs (other than hemispheric preferences).

**(j)** The interneuropil neurons innervating the CDNs receive input from multiple cell classes in other neuropil, with some pooling hundreds of synapses from descending neurons coming from the brain.

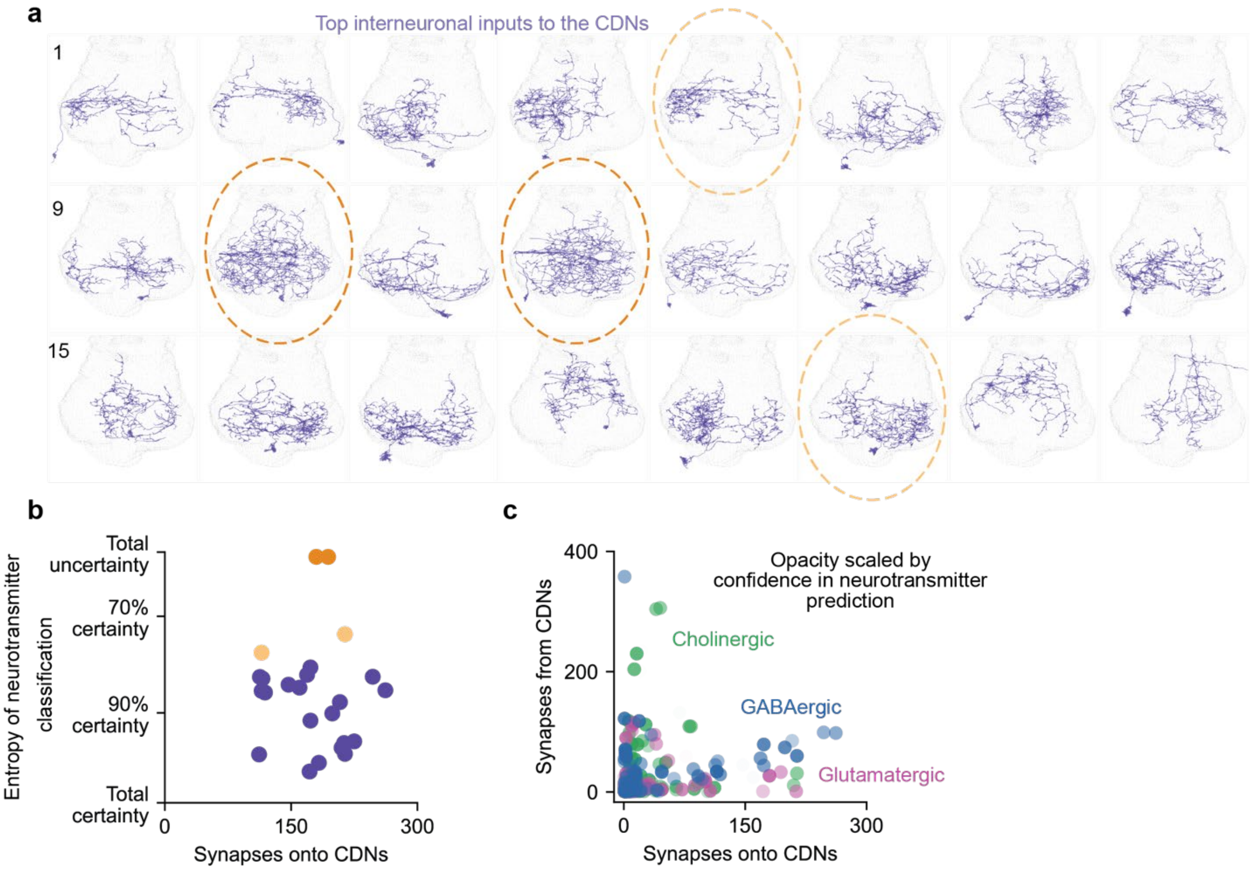


**Extended Data Figure 20: Local interneuron inputs to the putative CDNs are highly varied and likely inhibitory**

**(a)** The 24 strongest inputs to the CDNs from within the AG show highly varied anatomy, rather than resembling any specific cell class.

**(b)** The neurons with the most ambiguous neurotransmitter predictions (circled in panel **a**) do not have anatomy closely approximating the labeling by our TH-Gal4 line.

**(c)** The neurons that most strongly innervate the CDNs from within the AG are predicted to be inhibitory (either through GABA or glutamate) and do not receive reciprocal synapses from the CDNs, again arguing against a primary role for recurrence in their ability to integrate inputs.

#### VIDEO LEGENDS

**Video 1 | Silencing the CDNs prevents the termination of mating in response to heat threats.**
Silencing the CDNs with GtACR1 beginning 14.5 minutes into mating prevents the male from terminating the mating in response to a one-minute heat threat beginning at 15 minutes after the onset of copulation (12 seconds into the video). The male does not appear obviously stuck or unable to disengage his genitalia, but is also not paralyzed as evidenced by his attempts to retain the mating posture even as the female struggles to escape.

**Video 2 | Stimulation of the CDNs terminates the mating with variable latency.**
Several flies expressing CsChrimson in the CDNs are exposed to 2 seconds of light after 10 minutes of copulation. Each fly responds with a variable latency, often several seconds after the optogenetic stimulation.

**Video 3 | The termination procedure in response to threats resembles that of stimulation of the CDNs.**
A heat threat at 15 minutes into mating (beginning ~7 seconds into the video) results in termination behavior resembling optogenetic stimulation of the CDNs

**Video 4 | A short gust of wind strongly disturbs mating flies, but does not mechanically dislodge the male from the female**
Examples of CDN>GFP males challenged by a 350-millisecond gust of wind at 15 minutes into mating. In both examples, the gust of wind throws the mating pair around the chamber. Both male flies maintain their position on the female in the immediate aftermath of the wind gust. The male that terminates mating does so with a delay of ~3 seconds, indicating that it was not the force of the wind that caused the termination, but a decision made by the activity of the CDNs.

**Video 5 | Constitutively silencing the CDNs with tetanus toxin prevents termination in response to a gust of wind**
Example of a CDN>Tnt male that continues mating in response to a 500-millisecond gust of wind at 15 minutes into mating.

### METHODS

#### Fly stocks

Flies were maintained on conventional cornmeal-agar-molasses medium under 12 hour light/12 hour dark cycles at 25°C. Males were anesthetized with $CO_2$ and collected 0-10 days after eclosion and group-housed away from females for at least 3 days before testing. Flies expressing CsChrimson, GtACR1, or ChR2-XXM and all experimental controls were housed with rehydrated potato food (Carolina Bio Supply Formula 4-24 Instant Drosophila Medium, Blue) coated with all-trans-retinal (Sigma Aldrich R2500) diluted to 50 mM in ethanol for at least 1 day, unless marked as “no retinal.” These vials were kept inside aluminum foil sheaths to prevent degradation of the retinal due to light exposure. Virgin females used as partners for copulation assays were generated by heat-shocking a UAS-CsChrimson-mVenus stock with a hs-hid transgene integrated on the Y-chromosome (Bloomington stock #55135) in a 37°C water bath for 75 minutes. This stock was selected for mating partners because the females are highly receptive to courtship, resulting in many mating pairs shortly after initiation of assays, and because it has been shown that copulation duration is robust to variations in the female’s genetic background. Virgins were group-housed for at least 2 days before use. Optogenetic experiments were not performed at specific times relative to the light-dark cycle of the incubator because these animals were housed in constant dark conditions to preserve all-trans-retinal integrity. We did not observe any dependency of time of day on any of the behaviors described here. Flies containing TH-LexA and LexAop-TrpA1 (or flies used as controls for TH>TrpA1 experiments) were raised at 19°C and collected as adults before being stored at 25°C. Detailed genotypes of all strains used in the paper are listed in **Supplementary Table 3**.

#### BEHAVIORAL EXPERIMENTS

##### Evaluation of mating

A pair of flies was scored as “mating” when they adopted a stereotyped mating posture for at least 30 seconds. This posture consists of the male mounting the female and propping himself up on her abdomen using his forelegs, while curling his own abdomen and keeping the genitalia in contact. The posture is starkly different from anything exhibited during other naturalistic behaviors and is rarely, if ever, sustained for 30 seconds during unsuccessful attempts to initiate a mating. If the flies are physically pulled apart without disengaging the genitalia (such as if the female falls or if they collide with an obstacle), the male can climb back into place. This posture is also maintained in the presence of threats, unless the male elects to terminate the mating, differentiating it from a “stuck” phenotype. When stuck, the male dismounts the female, orients himself away from her, and attempts to walk away, but cannot decouple their genitalia. In the rare cases when we see a male become stuck in response to a threat, we record the copulation as having terminated. Occasionally in extremely long (>1hr) mating males, the male will become stuck, possibly because the seminal fluids harden and adhere the flies together. In this case too, the onset of the stuck posture is scored as the end of mating.

##### Assessing fertility

After a mating, the female fly was collected and placed in isolation in the above-described cornmeal food vials. One week later, the vial was visually inspected for the presence of larva as an indicator of successful fertilization.

##### Optogenetic stimulation during behavior

For more information on the behavioral arenas we used, see Thornquist et al., 2020.

*Non-screen experiments:*
For CsChrimson experiments: One male and one virgin female fly were placed in each 86" diameter 1/8" thick acrylic well sitting 4" above 655 nm LEDs (Luxeon Rebel, Deep Red, LXM3-PD01-0350) driven using 700 mA constant current drivers (LuxDrive BuckPuck, 03021-D-E-700) and passed through frosted collimating optics (Carclo #10124). This spot of light was scattered using a thin diffuser film (Inventables, 23114-01) under the wells to ensure a uniform light intensity of ~0.1 mW/mm$^2$. The LEDs were controlled using an Arduino Mega2560 (Adafruit) running a custom script, which itself was controlled by a Raspberry Pi (either 2 or 3, running Raspbian, a Debian variant). Flies were observed by recording from above using the Raspberry Pi with a Raspberry Pi NoIR camera (Adafruit) and infrared illumination from below using IR LED arrays (Crazy Cart 48-LED CCTV Ir Infrared Night Vision Illuminator reflected off the bottom of the box) while streaming the video to a computer for observation.

For GtACR1 experiments: The set-up was as above except using the green Luxeon Rebel, LXML-PM01-0100, and a pulse-width modulated signal to set the time-average intensity of the light to ~5 µW/mm$^2$ (approximately six times brighter than the ambient light) unless otherwise noted.

Matings must have ended within one minute after the end of CDN or grooming neuron stimulation in order to be scored as "terminated". Matings that did not stop within one minute of the stimulation's end almost always continued until ~23 total minutes of copulation (data not shown). Matings that continued longer than one minute after the end of stimulation but terminated earlier than the expected ~23 minutes were still scored as "not terminated".

Optogenetic experiments in which we recorded the exact time of termination in response to stimulation, or assigned mating termination to one of 10 pulses of red light, were scored *post hoc* from a recorded video of the experiment. We recognize that flies may make the decision to terminate mating before there are any visible signs, and that the termination motor process is typically executed over 1-3 seconds. Therefore, there may be a small amount of error in our observations of exactly how much stimulation time was needed to terminate matings, especially for the experiments in **Figure 3g** in which there was only a small amount of time (1 or 2.5 seconds) between red light pulses. It is possible there were flies that were recorded as terminating mating in response to a later red light pulse than was actually responsible. In order to be as accurate as possible, for flies that terminated mating in response to stimulation, we scored termination time as the first instance of the male pulling away from the female.

*Screen experiments (**Figure 3a**):*
Individual pairs of males and females were placed in single wells of 32 well recorded from a height of ~9'' using a Canon camera (VIXIA HF R600). Arenas were illuminated from below using a diffuse white light source (HSK A2 light pad on top of an Artograph LightPad 930 LX) with an illuminance of ~4500 lux. Copulation duration was scored *post hoc* using our automated scoring system[6] (https://github.com/CrickmoreRoguljaLabs/CaMKIICode).

**Thermogenetics (with and without optogenetics) and heat threats**
A similar device to the optogenetic arenas described above was constructed, with the addition of a ¼" thick water bath underneath each well. Room temperature water was continually passed through this bath, except when thermogenetic (TrpA1) or heat threat manipulations occurred. To change the temperature of the wells, hot water (controlled by a separate stopcock for each well) replaced the room temperature water. Heat threat temperatures refer to the temperature in the well after 10 seconds of hot water, as measured by a Bestdo 6802II digital thermometer. During

heat threat experiments, hot water was administered 10 seconds before intended time of the threat because it took ~10 seconds for the well to warm up to the right temperature. All heat threats lasted 1 minute, and matings were scored as "terminated" if they ended within 1 minute of the end of the heat threat. Like the optogenetic experiments, matings that did not stop within one minute of the heat threat's end almost always continued until ~23 total minutes of copulation (data not shown). The LEDs above were driven with 1A BuckPucks controlled by a pulse-width modulated signal selected to ensure the average intensity of illumination is the same as in the other behavioral experiments (~0.1 mW/mm$^2$ for red light, ~ 5 μW/mm$^2$ for green) despite having to pass through the water bath.

**Additional notes about acute optogenetics and heat threat experiments**

In most CDN/NP2719>Chr experiments, flies which did not terminate in response to the light pulse were probed with a four-second pulse of light 30-60s after the original light pulse to ensure that they expressed CsChrimson. In almost every mating pair, this was sufficient to induce termination, and in the few cases (<0.1%) in which it was insufficient, a subsequent 15 second pulse was likewise insufficient to terminate the mating, suggesting that these flies did not express CsChrimson (likely parental flies), and thus were not included in the reported data.

All reported termination probabilities are technically conditional: they are only the subset of flies which persisted in mating until the noted time of the stimulus. For the 15-minute and earlier time points, this accounts for 100% of experimental flies, but data at the 20-minute time point should be considered in this light, rather than as a cumulative termination probability that includes the flies that terminated without intervention.

**Experimental apparatus for delivering wind gusts to flies**

See **Extended Data Figure 2**. Flies were placed in modified cylindrical acrylic behavioral chambers (10mm diameter and ~9mm height) from Zhang et al., 2016. These chambers were placed on top of an acrylic box (9" x 9", 5.5" height, 0.125" thick, McMaster-Carr 8505K742), with the sides and area under the behavioral chambers cut out. This box was placed on top of another acrylic box (12" x 24", 4" height, 0.5" thick, McMaster-Carr 8560K266). In the floor of each behavioral chamber, there was a small hole (~3mm sided hexagon) that allowed a barbed tube fitting adapter (McMaster-Carr 5463K438) to fit in. To prevent flies from falling into the hole, a layer of mesh was placed on top of the floor. 0.125" inner diameter tubing (Firm Polyurethane, McMaster-Carr 5648K74) was inserted into each adapter, which then fed into 0.25" inner diameter tubing (Tygon PVC, McMaster-Carr 6516T21). This tubing fed through the cut-out sides of the acrylic box, with each individual tube connecting to a 2 way, normally closed solenoid (K-Rui H2O Solenoid, 12V, #997; 0.5" FNPT x 0.25" Nylon Female Adapter, Thogus Products TAF1084/N). Air was fed into the solenoid via two wall sources, with each source suppling air for five solenoids/chambers. Tubing from each air source fed into three airflow meters (Hilitand LZQ-7). Between each wall source and the airflow meters, a two-way stopcock was put in-line, to allow excess air to leak out. Of the three airflow meters, two meters fed into two solenoids each and one meter only fed into one solenoid. Between each airflow meter and the solenoids, another two-way stopcock was inserted in-line, to allow excess air to leak out. Solenoid-sized holes were cut into the 0.5" thick acrylic box to allow for solenoids to rest above the benchtop. To allow the air from the wall air sources to pass through the normally-closed solenoids, we connected each solenoid to an Arduino Mega 2560 and wrote custom Arduino and Java code to supply current to each solenoid. A camera (Canon Vixia HF R700) was set up above behavioral chambers and connected via HDMI cable to a computer monitor (Asus VS238H-P) to view and record experiments.

**Scoring mating termination in response to wind gusts**

Air strength was adjusted via air flow meters, and the strength of the wind delivered to each chamber was tested before starting experiments for the day with a handheld anemometer (BTMETER BT-100). All experiments were performed using a wind strength of ~1.5 meters/second. Wind speed was measured by removing each adapter from the bottom of its behavioral chamber – air was allowed to flow for 3.5 seconds and had to reach ~1.5 meters/second by the end of the 3.5 seconds.

Flies were scored as terminating mating if they stopped mating within 30 seconds of wind offset. For about 10% of matings, the wind gust either pinned flies against the wall (as opposed to throwing them around the chamber), or made flies get stuck in the door; these matings were excluded from the data.

**Note for paired wind gust experiments**
The Arduino code is very accurate in timing out how long to wait before opening solenoids in order to deliver gusts of wind to mating flies, as well as the length of interpulse intervals. However, it is not able to deliver a program of paired pulses to multiple behavioral chambers simultaneously. That means if delivering a paired pulse paradigm to two chambers, the first chamber must finish its full paired pulse cycle before the next one can begin. For example, if a paired pulse paradigm (500ms pulse, 10 sec interval, 500ms pulse) begins in chamber 1 at time 0:00 and is supposed to begin in chamber 2 at time 0:05, it will not begin in chamber 2 until time 0:11, after the whole cycle for chamber 1 has finished. This means that for flies in chamber 2, the pulse paradigm would not start until 6 seconds after the intended time. In order to correct for this, for paired pulse paradigms with an interpulse interval of 10 seconds or more, we started experiments by only loading flies into 5/10 mating chambers (so as to minimize the odds of flies mating within close time proximity of each other, and then loaded the other 5 chambers after the first 5 had already started mating. Since it was unlikely for flies to start mating within about 5 seconds of each other, we loaded all 10 chambers at the same time for experiments with an interpulse interval of 5 seconds. However, despite these precautions, there were still some flies whose paired pulse paradigms overlapped, and therefore caused one chamber's wind delivery to be delayed by a few seconds. Though not ideal, we consider these variations essentially negligible since our data indicates that the CDN time constant of integration changes over the timescale of many minutes.

#### IMAGING EXPERIMENTS

FLIM and intensity images were acquired, as in our previous work[13,14], using a modified Thorlabs Bergamo II. Samples were excited using a Coherent Chameleon Vision II Ti:Sapphire laser emitting a 920 nm beam and emission was detected using cooled Hamamatsu H7422P-40 GaAsP photomultiplier tubes, with light collected through a 16x water immersion objective (Nikon, 0.8NA N16XLWD-PF). The PMT signal was amplified using Becker-Hickl fast PMT amplifiers (HFAC-26) and passed to a PicoQuant TimeHarp 260 photon counting board, which was synchronized to the laser emission by a photodiode (Thorlabs DET110A2) inverted using a fast inverter (Becker-Hickl A-PPI-D). The TimeHarp signal was acquired by custom software (FLIMage, Florida Lifetime Imaging, Version 2.0.21) which was also used to control the microscope. Optogenetic stimulation was performed by excitation with a blue (Thorlabs M470L4) or red (Thorlabs M625L4) LED through a liquid light guide (Thorlabs LLG5-8H) for an incident intensity at the sample of ~0.2 $mW/mm^2$.

We acquired 64 x 64 pixel FLIM and intensity images at a rate of ~7.8 Hz (1 frame every 0.128 seconds). Each data point reported is the average of 40 frames (5.12 seconds). FLIM images were automatically averaged during acquisition. Intensity images were averaged post hoc.

**Dissection**
Flies were anaesthetized with ice, then dipped in 100% ethanol for about 15 seconds. The ventral nervous system was dissected out in chilled saline (103 mM NaCl, 3 mM KCl, 5 mM TES, 8 mM trehalose, 10 mM glucose, 26 mM $NaHCO_3$, 1 mM $NaH_2PO4$, 4 mM $MgCl_2$, 1.5 mM $CaCl_2$, pH ~7.25, 270-275 mOsm, dissolved in deionized water). The dissected out nervous system was then carefully transferred into a 35x10 mm VWR petri dish filled with 3 mL of the same chilled saline. The nervous system adhered to the bottom of the petri dish and was stable throughout the experiment.

*Note: If flies were anaesthetized with $CO_2$, baseline CaMKII activity was increased in the CDNs, and CaMKII was unresponsive to ChR2-XXM laser stimulation.*

**Pharmacology**
Dopamine was prepared as a 100 mM stock solution in $ddH_2O$ and kept frozen. From this, 10 mM aliquots were diluted in $ddH_2O$ were used for experiments, which were dissolved in chilled saline at different ratios depending on the experiment (for saline controls, the same amount of ddH2O was dissolved). The final volume was always 40 mL. For example, to get a 100 µM dopamine solution, 400 µL of 10 mM dopamine was dissolved into 39.6 mL of chilled saline — slightly modifying concentrations, osmolarity, and pH, which was not corrected.

Dopamine (or saline) was delivered by perfusion during experiments. Prior to application of dopamine, or in control cases, matched saline solution, data was collected with no perfusion. A 50 mL graduated cylinder (in which we put the dopamine solution) was kept elevated above the sample by a three-prong clamp attached to the top of a three-foot-tall stand. 1/8" internal diameter tubing (Firm Polyurethane, McMaster-Carr 5648K74) was fed into the graduated cylinder; it was then passed into the box that enclosed the objective, stage, scan mirrors, PMTs, etc. of our microscope. 1/32" internal diameter tubing (Clear Masterkleer Soft PVC, McMaster-Carr 5233K91) was then connected to the original tubing via a 1000 µL pipette tip (VWR 76322-154), allowing us to fit a 10 µL pipette tip (VWR 53509-130) at the end, which was held in place by a stand attached to the microscope stage. The pipette tip was then submerged in the saline in the petri dish. In order to move the dopamine from the graduated cylinder to the petri dish we inserted a 20 mL LiteTouch luer lock syringe in line with the 1/8" diameter tubing via a three-way stopcock. To deliver dopamine, we pulled about 5 mL of solution into the syringe from the graduated cylinder, and then gently pushed about 2 mL of solution from the syringe to the sample, which allowed the solution to naturally flow from the graduated cylinder to the sample at a rate of ~4 mL/min.

To remove solution from the petri dish, so as not to let it overflow, we attached another 10 µL pipette tip adjacent to the input pipette tip in such a way that it was positioned above the dopamine/saline solution but below the top of the petri dish. This allowed the dopamine/saline solution to remain at a constant volume within the petri dish, since it was removed once it rose up to the output pipette tip. The output pipette tip was attached to the 1/8" diameter tubing, which was then passed out of the microscope-enclosing box and into a 250 mL bolt neck flask with upper tubulation. A rubber stopper with a hole was inserted into the mouth of the flask – this was used to attach the flask to a vacuum. The vacuum allowed solution to be removed from the output pipette tip and be deposited into the 250 mL bolt neck flask.

After experiments were done for the day, 1-2 liters of deionized water were run through the input tubing of the perfusion system (emptying into an external flask) to wash out any residual dopamine.

**Region of interest**
For each sample, one region of interest (ROI) was drawn around the axons of the CDNs. We used the driver NP5270 Gal4 because it is a stronger driver and took less laser power to image from than NP2719, while still recapitulating the behavioral phenotypes of NP2719 and sparsely labelling the abdominal ganglion[12]. The neurons labeled by NP5270 project to either side of the abdominal ganglion. One set of axonal projections was chosen to record from, depending on its strength of fluorescence. Extra care was taken to not include any cell bodies in the region of interest. For intensity imaging, a small region of interest was also drawn outside of the abdominal ganglion and set as background.

**Optogenetic stimulation during imaging**
We used two types of optogenetic stimulation: stimulation with visible light (625 nm red light stimulation of CsChrimson and 470 nm blue light stimulation of ChR2-XXM) and stimulation of ChR2-XXM with the 920 nm infrared light emitted from our laser. When doing experiments not involving infrared optogenetic stimulation, we used a laser power of ~5 mW (as measured by a Coherent FieldMate Laser Power Meter 1098948; measurements using this power meter were slightly noisy – they varied by ~0.2 mW).

For one-photon stimulation experiments, we first recorded the baseline measurements for our sensor over a variable amount of time (always more than 30 seconds) depending on the experiment. Our 2-photon laser and PMTs were shuttered during stimulation to protect the PMTs using an Arduino, which closed the shutter 100 milliseconds before and opened the shutter 100 milliseconds after the stimulation.

For infrared laser optogenetic stimulation of CsChrimson or ChR2-XXM (the pre/post dopamine or saline experiments), we first found the region of interest (ROI) using ~5 mW laser power. We then shut the laser off and waited ~1 minute to allow the fluorescence to return back to baseline. We then recorded for 204.8 seconds at ~10 mW laser power, which constituted the "pre dopamine" period. We then turned the laser off and started perfusing in dopamine. We did not continue recording while perfusing dopamine because the start of perfusion sometimes moved the sample out focus, though we otherwise noticed no difference in CaMKII dynamics during these continuous imaging/stimulation trials (**Extended Data Figure 10g**). Once dopamine perfusion started, we quickly confirmed that the sample had not moved by turning the laser back on (~10 mW, but not recording) and brought it back into focus if it had moved. We then turned the laser off and waited until we could see the output tubing start to remove solution from the petri dish (see Pharmacology section above) - typically this whole process typically took ~2 minutes. Then we started recording again, which constituted the "dopamine" or "saline" (for **Figures 4e and f**) period. For some dopamine concentration experiments (**Figure 4f**) and the calcium imaging experiments in **Figure 4e**, the "pre dopamine/saline" period traces are not shown.

**Fluorescence lifetime imaging**
The FLIMage software recorded the time of arrival of each photon emitted (from the eGFP molecule of green-Camuiα) relative to the time of the excitation pulse delivered by our laser for each pixel in 64 time bins of width 200 picoseconds. The mean/average fluorescence lifetime was calculated using custom python code (https://github.com/CrickmoreRoguljaLabs/).

Briefly, we drew a ROI around the axons of the CDNs (see *Region of Interest*, above); the photon distributions for each pixel within the ROI across 40 consecutive frames (see *Imaging Experiments*, above) were pooled to increase photon count and reduce noise. To calculate the fluorescent lifetimes, these pooled photon distributions (represented as a vector n, where $n_i$ is the number of photons in the ith time bin) were used to fit the parameters of a double exponential model (1) that was convolved with a Gaussian Instrument Response Function (2) (resulting in (3)).

$$\boldsymbol{DExp(t) = \alpha_1\, exp\left(\frac{-t}{\tau_1}\right) + \alpha_2\, exp\left(\frac{-t}{\tau_2}\right)}$$

(1)

$$\boldsymbol{IRF(t) = \frac{1}{\tau_g\sqrt{2\pi}}exp\,(-\frac{(t-t_0)^2}{2\tau_g^2})} \quad (2)$$

$$\boldsymbol{(DExp * IRF)(t)} \quad (3)$$

Specifically, the parameters were calculated by minimizing the mean squared error between n and Eq. 3 as expressed below, subject to$\boldsymbol{\Sigma_n \alpha_n = 1, 0 \leq \alpha_n \leq 1, 0 < \tau_g, \tau_0, \tau_n}$.

$$\boldsymbol{\underset{\alpha_n,\tau_g,\tau_0,\tau_n}{argmin} \frac{1}{64}\sum_{i=1}^{64}(n_i - (DExp * IRF)(i))^2} \quad (4)$$

The mean/average fluorescence lifetime ($\boldsymbol{\tau_{mean}}$) was calculated as in our previous work[13,14]:

$$\boldsymbol{\tau = \frac{\sum_{i:t_i \geq t_0} n_i t_i}{\sum_{i:t_i \geq t_0} n_i} - t_0} \quad (5)$$

**Additional note about FLIM experiments**
In a very small number of trials (2 or 3), ChR2-XXM stimulation of the CDNs/NP5270 Gal4 did not increase green-Camuiα lifetime. These samples were subsequently probed with multiple 10 second blue light bouts, which also had no effect on lifetime. ChR2-XXM is not fused with a fluorescent protein so we could not easily verify its absence. However, due to the very reliable induction of CaMKII activity in the CDNs by strong ChR2-XXM stimulation (**Extended Data Figure 10a**, and dozens of other unreported experiments), we concluded that these samples likely did not express ChR2-XXM and were parental flies, so they were excluded from our data.

**Calcium imaging**
For calcium imaging, all detected photons within a pixel were summed together, regardless of arrival time relative to the excitation pulse. All data were likely entirely due to functioning GCaMP6s molecules, as opposed to autofluorescence, as their estimated fluorescence lifetime was 2.6-2.8 nanoseconds (data not shown).

For all experiments, change in fluorescence ($\boldsymbol{\Delta F/F}$) of GCaMP6s after optogenetic stimulation of the CDNs was calculated using custom MATLAB code (www.github.com/CrickmoreRoguljaLabs/) as:

$$\Delta \boldsymbol{F/F = \frac{F_1 - F_0}{F_0}}$$

where $\boldsymbol{F_0}$ is the mean number of photons (over at least 30 seconds) recorded at baseline before optogenetic excitation and $\boldsymbol{F_1}$ is the number of photons collected in each frame after excitation.

The code calculates $\Delta F$ from a .csv file that is created by the FLIMage software after an experiment. The file is in the format of photon count per frame (among other measurements), and accounts for background photon ROI subtraction.

The only exception is for the experiment in **Figure 4e** (further quantified in **Extended Data Figure 10e**) in which optogenetic stimulation (from the laser itself) is constantly present, so $F_0$ was calculated from the first 5.12 seconds of recording of the *pre* dopamine/saline period – the pre period is not shown in the traces. The original $F_0$ value from the pre period was also used for the post dopamine/saline period.

***<u>Experiments in Figure 4b and Extended Data Figure 14:</u>***

Physiology experiments in **Figure 4b** and **Extended Data Figure 14** were performed using an Olympus 20x 1.0 NA water immersion objective (XLUMPLFLN) on a customized Sutter MIMMs to be detailed in a later manuscript by SCT and Gaby Maimon. All data were plotted with `matplotlib` version 3.7.2. Extracellular saline was prepared as in Mussells Pires *et al.*, 2024[52] consisting of (in mM concentrations of the sats used): 103 NaCl, 3 KCl, 5 TES, 10 trehalose dihydrate, 10 glucose, 2 sucrose, 26 NaHCO3, 1 NaH2PO4, 1.5 CaCl2 and 4 MgCl2 titrated with Milli-Q water to an osmolarity of 280-285 mOsm. All salts were sourced from Sigma-Aldrich. Dopamine hydrochloride salt (Sigma H8502) was added to 50 mL aliquots of saline to the desired concentrations of 1, 5, 10, and 100 µM used for individual experiments. Images were acquired at a frame rate of ~61 Hz at 256 x 256 pixel resolution.

The physiology experiments were performed in a custom physiology chambers made of laser-cut ¼” thick black acrylic from McMaster-Carr (8505K755) cut in the shape of a standard sample slide (76 x 26 mm) with a 17 mm x 17 mm square hole in the center of the slide. The sides of one face of the hole were chamfered to provide access for a perfusion system and a thermocouple to measure bath temperature around the objective. The base of the hole (still 17.5 x 17.5 mm) was covered with a glass coverslip adhered to the chamber by placing a layer of Parafilm between the acrylic and coverslip, then heating the coverslip with a soldering iron to melt the Parafilm. The Parafilm was then cut away with a razor blade to provide a glass surface that contacted the dissected nervous system. The glass coverslips were covered with a poly-L-lysine solution overnight before use, which was washed away before experiments, to increase the stickiness of the glass surface. Saline was perfused driven by gravity with flow control set to a rate of ~3 mL/min through Tygon tubing formulation 2375. Saline was cleared from the chamber with a vacuum line connected to a thin glass wick with a hole through which saline was sucked to avoid any contact with potential contaminating plastics in the outlet tubing.

Closed-loop temperature control was achieved using a Warner Instruments temperature controller (CL-100) driving a Peltier device (SC-20) that enclosed the perfusion line and regulated by the LCS-1 heat exchanger. The set point was controlled, and probe temperatures were measured, with the BNC outputs of the CL-100 driven and measured by an MCC USB-1208FS DAQ controlled with custom Python3 code using the ROS2 Foxy Fitzroy distribution. The probe was placed in the bath of the physiology chamber.

Dopamine was applied by thawing 50 mL aliquots of each dose and switching the perfusion line into each dopamine solution sequentially (timestamped in the siff file stream to know when the solution was switched). Each solution was given ~5 minutes to fill the perfusion system and physiology chamber before switching solutions.

Male flies aged at least 3 days after eclosion were anesthetized on ice, and their VNS and brain was dissected out together in the extracellular saline solution above, with care taken not to contact the VNS with the forceps and disrupt the glial sheath. The VNS was then pipetted into the physiology chamber and allowed to settle for ~5 minutes before beginning perfusion of saline.

For thermogenetic stimulation, the explant was imaged for several minutes at ~20°C before increasing the Peltier controller's set point to ~39°C. The temperature was monitored until the water bath achieved ~30°C before resetting the set point to 20°. Each brain was exposed to two steps to 30°C to ensure repeatability of the result, but the first temperature ramp in each experiment was used for quantification. In flies expressing TrpA1 under the control of TH-LexA, we noticed that often the fluorescence signal would decline before the temperature was stepped back to 20°C – we interpret this result as habituation of the TrpA1 channel or a depletion of endogenous dopamine stores, as bath application of dopamine did not show the same effect (and so presumably does not result from bleaching of the fluorophore at high dopamine concentrations). To quantify fluorescence or lifetime, we took the mean intensity-weighted lifetime or ΔF/F over a 30 second window just preceding the command to increase the temperature (for the 20°C case) or during the period that the temperature was stably elevated to 30°C. We discarded approximately 1/3$^{rd}$ of flies because of either motion artifacts or formation of an air bubble directly above the sample during the temperature ramp. $F_0$ was computed using the 5$^{th}$-35$^{th}$ second of the recording, so sometimes the 20°C condition mean varied slightly from 0. Plotted data use a boxcar filter of width 0.5 seconds to smooth the traces.

Fluorescence lifetime for GRAB-DA3m experiments were computed by fitting the first 1200 frames to a biexponential distribution convolved with a Gaussian by minimizing the chi-squared statistic, as in Thornquist *et al.*, 2020. The offset of this fit was used to adjust the mean photon arrival time within each frame. Mean intensity-weighted lifetime refers to the following procedure: when taking the mean of a set of $\boldsymbol{n}$ frames, the lifetime was computed as

$$\langle \boldsymbol{L} \rangle = \sum_{\boldsymbol{k}=1}^{\boldsymbol{n}} \frac{\boldsymbol{I_k L_k}}{\boldsymbol{n}\langle \boldsymbol{I} \rangle}$$

where $\boldsymbol{I_k}$ refers to the intensity of the $\boldsymbol{k}$th frame, $\boldsymbol{L_k}$ refers to the mean lifetime of the $\boldsymbol{k}$th frame, and $\langle \circ \rangle$ refers to the average across frames.

**Antibodies and immunohistochemistry**

All samples were fixed in PBS with Triton X-100 and 4% paraformaldehyde for 20 minutes, then washed three times with PBS/Triton X-100 for 20 minutes each before application of antibodies. All samples were incubated with the primary antibody for two days, washed three times with PBS/Triton X-100 for 20 minutes each, incubated with the secondary antibody for two days, then washed three times as before and mounted on coverslips using VectaShield (Vector Labs). The exception is for MCFO staining, in which we followed the protocol of Nern et al.[53]
Antibodies used are as follows:

Rabbit anti-GFP (A-11122, Invitrogen, 1:1,000 dilution)
Chicken anti-GFP (GFP-1010, Aves Labs, 1:1,000 dilution)
Mouse anti-GFP (A11120, Invitrogen, 1:2,000 dilution)
Rabbit anti-DsRed (323496, Clontech 1:1000 dilution)
Mouse anti-nc82 (Developmental Studies Hybridoma Bank)
Donkey anti-chicken 488 (703-545-155, Jackson ImmunoResearch, 1:400)

Donkey anti-rabbit 488 (A11008, Invitrogen, 1:400 dilution)
Donkey anti-mouse 488 (A21202, Invitrogen, 1:400 dilution)
Donkey anti-rabbit 555 (A-31572, Invitrogen, 1:400 dilution)
Donkey anti-mouse Cy3 (715-166-150, Jackson ImmunoResearch, 1:400 dilution)
Donkey anti-rabbit 647 (711-605-152, Jackson ImmunoResearch, 1:400 dilution)
Donkey anti-mouse 647 (711-605-151, Jackson ImmunoResearch, 1:400 dilution)

**Confocal microscopy**
Confocal images were collected using a Zeiss LSM 710 through a 20x air objective (Olympus PLAN-APOCHROMAT) controlled by Zen software and analyzed using ImageJ.

**Connectomics Analysis**

All analyses were performed using Python 3.9 code using the NeuPrint Python API version 0.4.26 on the MANC volume v1.0. We first printed single-plane projections, flattening the dorsoventral axis, of all neurons whose `input_roi` and `output_roi` were only `'ANm'` and which did not enter or exit via any nerves (using `Bokeh` to generate visualizations of the neurons' skeletons and synapses). We manually inspected each neuron for features that qualitatively matched the CDNs or the Crz neurons, and created a candidate list as a starting point for further analyses. All visualizations of neuronal anatomy used `Bokeh 3.2.2`, but otherwise visualizations used `matplotlib`.

*Putative Crz neurons*

We queried the NeuPrint API for the number of release sites and synapses received from all other neurons in the MANC volume for each putative Crz neuron, and computed the cosine similarity of their output and input vectors using `scikit-learn 1.3.2`. We plotted the similarity between each candidate neuron as a heatmap, both for the presynaptic and the postsynaptic vectors, ordered using `sklearn`'s `linkage` (with the 'average' metric for cluster distances) and `dendrogram` functions to suggest clusters of neurons with similar connectivity, chosen subjectively. For each putative cluster, we examined the morphology of individual cells more closely, ruling out potential Crz neurons based on features like the presence of an extra midline branch crossing the anterior abdominal ganglion. This resulted in six neurons, with body IDs 14161, 11828, 13190, 16132 , 17702, 14938, that appeared to be putative Crz neurons. We chose the set of four of these which were most similar to one another to identify as Crz neurons, though our superficial analyses seem qualitatively robust to the selection of which set of four is used.

*Putative CDN anatomy*

We filtered the anatomically CDN-like cells by predicted neurotransmitter, taking all cells that were predicted to be more likely to be GABAergic than cholinergic or glutamatergic. We then used the same clustering approach as with the Crz neurons, focusing on the presynaptic vectors for each putative GABAergic CDN, to identify the five clusters delineated in **Extended Data Figure 16**.

Cluster 2 was selected based on the numerosity of the neurons (7 identified) and the closer consistency of their morphology with that of the CDNs than other potential clusters.

*Putative CDN outputs and inputs*

Overlaid images of neurons within a cell class were plotted using an opacity of $\mathbf{0.7}\frac{\text{synapses}}{\text{synapses}_{\boldsymbol{Max}}}$ where $\text{synapses}_{\boldsymbol{Max}}$ refers to the number of synapses of the cell within that class with the most synapses from or onto the CDNs. When scaling the opacity of markers within a scatter plot by neurotransmitter prediction confidence, the opacity was set to

$$\mathbf{1} - \frac{\text{Entropy}(\boldsymbol{x})}{\text{Log}_2(\mathbf{3})}$$

with

$$\text{Entropy}(\boldsymbol{x}) = -\boldsymbol{p}_{\boldsymbol{G}\text{ABA}}\text{Log}_2(\boldsymbol{p}_{\text{GABA}}) - \boldsymbol{p}_{\text{Glutamate}}\text{Log}_2(\boldsymbol{p}_{\text{Glutamate}}) - \boldsymbol{p}_{\text{Ach}}\text{Log}_2(\boldsymbol{p}_{\text{Ach}})$$

In **Extended Data Figure 20**, the assignment of entropy to confidence score was calculated mapping entropy to the confidence in the strongest neurotransmitter if the other two were equiprobable (i.e. the minimum possible confidence for a given entropy). Total uncertainty corresponds to an entropy of $\text{Log}_2(\mathbf{3}) = \mathbf{1.58}$, 70% confidence corresponds to an entropy of 1.181, 90% confidence corresponds to an entropy of 0.567, and total confidence corresponds to an entropy of 0.

#### MODELING

##### Estimation of parameters, $\boldsymbol{p_0}$ and $\boldsymbol{\tau}$

The instantaneous probability of termination, given that a fly has not yet terminated the mating by time $t$, was modeled as

$$p(t_{term} = t | \neg t_{term} < t) = p_0\tau(1 - e^{-\frac{t}{\tau}})$$

with $p_0$ the intensity of the stimulus and $\tau$ the time constant of integration, two model parameters we wish to fit. Each moment in time gives an approximate estimate of this value, which can be pooled together using the cumulative distribution $\sigma(t) = p(t_{term} \leq t)$, which can be derived by noting that

$$\frac{d\sigma}{dt} = p(t_{term} = t | \neg t_{term} < t)\big(1 - \sigma(t)\big) = p_0\tau(1 - e^{-\frac{t}{\tau}})(1 - \sigma(t))$$

this can be solved to yield

$$-\,log(1 - \sigma) = p_0\tau\left(t + \tau e^{-\frac{t}{\tau}}\right) + C$$

with $C$ the constant of integration. Because $\sigma(0) = 0$, we can immediately see that $C = -p_0\tau^2$ and solve to get

$$log(1 - \sigma(t)) = p_0\tau\left(\tau(1 - e^{-\frac{t}{\tau}}) - t\right)$$

finally yielding

$$\sigma(t) = 1 - e^{-p_0\tau\left(t - \tau(1 - e^{-t/\tau})\right)}.$$

The probability $p_{p_{0,}\tau}(x)$ of any particular observation $x$, given parameters, $p_0$ and $\tau$ is then

$$p_{p_0,\tau}(x) = \left.\frac{d\sigma}{dt}\right|_{t=x} = p_0\tau(1 - e^{-x/\tau})e^{-p_0\tau\left(x - \tau(1 - e^{-x/\tau})\right)}$$

We can now attempt to fit our data to this model to obtain estimates of the parameters $p_0$ and $\tau$. One form of estimate is the maximum likelihood estimate, the values of the parameters “least surprised” by the data (those which predict the greatest likelihood of generating the data set). The likelihood $\mathcal{L}_{\{x\}}(p_0, \tau)$ of any particular data set $\{x\}$, assuming all samples are independent, is

$$\mathcal{L}_{\{x\}}(p_0,\tau) = \prod_x p_{p_0,\tau}(x)$$

and we choose as our estimates $\hat{p}_0, \hat{\tau} = argmax_{p_0,\tau}\mathcal{L}_{\{x\}}(p_0,\tau)$.

Any values $\hat{p}_0$ and $\hat{\tau}$ that maximize $\mathcal{L}_{\{x\}}(p_0,\tau)$ also maximize $log\, \mathcal{L}_{\{x\}}(p_0,\tau)$, or

$$log\, \mathcal{L}_{\{x\}}(p_0,\tau) = \sum_x log\big(p_{p_0,\tau}(x)\big)$$

However, our data is subject to an additional constraint: the stimulation is terminated at some upper bound value $u$. Thus, rather than consisting of termination times $\{x\}$, the data takes the form $\{x\} \cup \{\emptyset\}^{n_{per}}$ with $n_{per}$ the number of data points that persevere through the threat. The probability of observing $\emptyset$ is the probability that the corresponding sample $x$ would be greater than $u$, or $1 - \sigma(u)$. Then we can then account for this constraint in the likelihood function by re-writing $\mathcal{L}_{\{x\}}(p_0,\tau)$ as

$$\mathcal{L}_{\{x\}}\big(p_0,\tau\big) = \left(\prod_{x\in\{x\}} p_{p_0,\tau}(x)\right)\big(1-\sigma(u)\big)^{n_{per}}$$

and thus

$$log\, \mathcal{L}_{\{x\}}(p_0,\tau) = \sum_x log\big(p_{p_0,\tau}(x)\big) + n_{per}\, log\big(1-\sigma(u)\big)$$

We can then maximize this expression to obtain our estimates $\hat{p}_0$ and $\hat{\tau}$.

To do so, we take the gradient of $log\, \mathcal{L}_{\{x\}}(p_0,\tau)$ and find where it equals 0.

$$\frac{\partial\, log\, \mathcal{L}_{\{x\}}(p_0,\tau)}{\partial p_0} = \sum_{x\in\{x\}} \frac{\partial\, log\big(p_{p_0,\tau}(x)\big)}{\partial p_0} + \frac{n_{per}\partial\, log\big(1-\sigma(u)\big)}{\partial p_0}$$

$$= \left[\sum_{x\in\{x\}} \frac{\partial}{\partial p_0}\left(log\left(p_0\tau\big(1-e^{-x/\tau}\big)e^{-p_0\tau\left(x-\tau\left(1-e^{-x/\tau}\right)\right)}\right)\right)\right] - n_{per}\tau\left(u-\tau\left(1-e^{-\frac{u}{\tau}}\right)\right)$$

$$= \left[\sum_{x\in\{x\}} \frac{\partial}{\partial p_0}\left(log(p_0\tau) + log\big(1-e^{-x/\tau}\big) - p_0\tau\left(x-\tau\big(1-e^{-x/\tau}\big)\right)\right)\right] - n_{per}\tau\left(u-\tau\left(1-e^{-\frac{u}{\tau}}\right)\right)$$

$$\frac{\partial\, log\, \mathcal{L}_{\{x\}}(p_0,\tau)}{\partial p_0} = \left[\sum_{x\in\{x\}} \frac{1}{p_0} - \tau\left(x-\tau\left(1-e^{-\frac{x}{\tau}}\right)\right)\right] - n_{per}\tau\left(u-\tau\left(1-e^{-\frac{u}{\tau}}\right)\right)$$

Similarly,

$$\frac{\partial\, log\, \mathcal{L}_{\{x\}}(p_0,\tau)}{\partial \tau} = \sum_{x\in\{x\}} \frac{\partial\, log\big(p_{p_0,\tau}(x)\big)}{\partial \tau} + \frac{n_{per}\partial\, log\big(1-\sigma(u)\big)}{\partial \tau}$$

Splitting into two pieces:

$$\frac{n_{per}\partial\, log\big(1-\sigma(u)\big)}{\partial \tau} = n_{per}p_0\left(2\tau\big(1-e^{-u/\tau}\big) - u\big(1+e^{-u/\tau}\big)\right)$$

and

$$\sum_{x\in\{x\}} \frac{\partial\, log\left(p_{p_0,\tau}(x)\right)}{\partial \tau} = \sum_{x\in\{x\}} \frac{\partial}{\partial \tau}\left(log(p_0\tau) + log\left(1-e^{-\frac{x}{\tau}}\right) - p_0\tau\left(x-\tau\left(1-e^{-\frac{x}{\tau}}\right)\right)\right)$$

$$= \sum_{x\in\{x\}} \left( \frac{1}{\tau} - \frac{xe^{-x/\tau}}{\tau^2(1+e^{-x/\tau})} - p_0 \left( 2\tau \left( e^{-\frac{x}{\tau}} - 1 \right) + x(1+e^{-x/\tau}) \right) \right)$$

$$= \sum_{x\in\{x\}} \left( \frac{\tau(1 + p_0\tau(2\tau - x)) - (x + \tau + 4p_0\tau^3)e^{-\frac{x}{\tau}} + p_0\tau^2 e^{-\frac{2x}{\tau}}(x + 2\tau)}{\tau^2(1 - e^{-x/\tau})} \right)$$

At the maximum likelihood values $\hat{p}_0$ and $\hat{\tau}$, both of these are 0, so

$$0 = \left[ \sum_{x\in\{x\}} \frac{1}{\hat{p}_0} - \hat{\tau}\left( x - \hat{\tau}\left(1 - e^{-\frac{x}{\hat{\tau}}}\right)\right)\right] - n_{per}\hat{\tau}\left( u - \hat{\tau}\left(1 - e^{-\frac{u}{\hat{\tau}}}\right)\right)$$

or

$$\frac{n_{term}}{\hat{p}_0} = \left[ \sum_{x\in\{x\}} \hat{\tau}\left( x - \hat{\tau}\left(1 - e^{-\frac{x}{\hat{\tau}}}\right)\right)\right] + n_{per}\hat{\tau}\left( u - \hat{\tau}\left(1 - e^{-\frac{u}{\hat{\tau}}}\right)\right)$$

where $n_{per}$ is the number of flies which terminate the mating in response to the stimulus. If $n$ is the total number of flies in the experiment, so that $\frac{n_{term}}{n} = p_{term}$ and $\frac{n_{per}}{n} = 1 - p_{term}$ then we have

$$\hat{p}_0 = \frac{p_{term}/\hat{\tau}}{(1 - p_{term})\left( u - \hat{\tau}\left(1 - e^{-\frac{u}{\hat{\tau}}}\right)\right) + \frac{\sum_{x\in\{x\}}\left( x - \hat{\tau}\left(1 - e^{-\frac{x}{\hat{\tau}}}\right)\right)}{n}}$$

which is the exact maximum likelihood estimate for $p_0$ given $\tau$.

Similarly, we have (after dividing by $n$) that

$$0 = (1 - p_{term})\hat{p}_0 \left( 2\hat{\tau}\left(1 - e^{-\frac{u}{\hat{\tau}}}\right) - u\left(1 + e^{-\frac{u}{\hat{\tau}}}\right)\right)$$
$$+ \frac{1}{n}\sum_{x\in\{x\}} \left( \frac{1}{\hat{\tau}} - \frac{xe^{-x/\hat{\tau}}}{\hat{\tau}^2\left(1 + e^{-x/\hat{\tau}}\right)} - p_0 \left( 2\hat{\tau}\left( e^{-\frac{x}{\hat{\tau}}} - 1\right) + x\left(1 + e^{-x/\hat{\tau}}\right)\right)\right)$$

We were unable to find an analytical solution to this equation, so to estimate $\hat{\tau}$ we performed grid search to maximize the log-likelihood, for each value using the analytical value for $\hat{p}_0$ given the current estimate for $\tau$. We used a step size of 0.001, and searched parameters ranging from 0.01 to 40.

Likewise, for the model in which there is no $\tau$ (**Extended Data Figure 2c**), the maximum likelihood estimate for $p_{0,null}$ is

$$\hat{p}_{0,null} = \frac{p_{term}}{(1 - p_{term})u + \frac{1}{n}\sum_{x\in\{x\}} x}$$

**Estimating variance of parameter fits**

To estimate the variance of the parameter fits, we use the Fisher Information and Cramér-Rao bound, which says

$$Cov(\boldsymbol{\theta}) \geq \mathcal{I}(\boldsymbol{\theta})^{-1}$$

for any consistent estimator of a parameter vector $\boldsymbol{\theta}$ where $\mathcal{I}(\theta)$ is the Fisher Information matrix defined as

$$\mathcal{I}(\theta)_{ij} = -\mathbb{E}\left[\left(\frac{\partial^2 \log p(\{x\}|\boldsymbol{\theta})}{\partial\theta_i \partial\theta_j}\right)\right]$$

For our model,

$$\mathcal{I}\begin{pmatrix}\hat{\tau}\\ \hat{p}_0\end{pmatrix} = -\mathbb{E}\left[\begin{pmatrix}\frac{\partial^2 \log p\left(\{x\}\middle|\hat{\tau},\hat{p}_0\right)}{\partial\tau^2} & \frac{\partial^2 \log p\left(\{x\}\middle|\hat{\tau},\hat{p}_0\right)}{\partial\tau\partial p_0}\\ \frac{\partial^2 \log p\left(\{x\}\middle|\hat{\tau},\hat{p}_0\right)}{\partial\tau\partial p_0} & \frac{\partial^2 \log p\left(\{x\}\middle|\hat{\tau},\hat{p}_0\right)}{\partial p_0^2}\end{pmatrix}\right]$$

The bottom right element is simply

$$\frac{\partial^2 \log p\left(\{x\}\middle|\hat{\tau},\hat{p}_0\right)}{\partial\tau^2} = -\frac{n_{term}}{\hat{p}_0^{\,2}}$$

so that the expectation is $\mathbb{E}\left[\frac{\partial^2 \log p\left(\{x\}\middle|\hat{\tau},\hat{p}_0\right)}{\partial\tau^2}\right] = -\frac{n\sigma(u)}{\hat{p}_0^{\,2}}$

The diagonal terms are

$$\frac{\partial^2 \log p\left(\{x\}\middle|\hat{\tau},\hat{p}_0\right)}{\partial\tau\partial p_0} = n_{per}\left(2\hat{\tau}\left(1-e^{-u/\hat{\tau}}\right) - u\left(1+e^{-\frac{u}{\hat{\tau}}}\right)\right) + \sum_{x\in\{x\}} 2\hat{\tau}\left(1-e^{-\frac{x}{\hat{\tau}}}\right) - x\left(1+e^{-\frac{x}{\hat{\tau}}}\right)$$

The expectation of the terms on the left (those resulting from the truncation) is

$$n(1-\sigma(u))\left(2\hat{\tau}\left(1-e^{-u/\hat{\tau}}\right) - u\left(1+e^{-\frac{u}{\hat{\tau}}}\right)\right)$$

The top left term of the Fisher Information is a little messy. The contribution from the values above the cutoff $u$ is

$$n(1-\sigma(u))p_0\left(\frac{(u^2+2u\tau)e^{-\frac{u}{\tau}}}{\tau^2} - 2\left(1-e^{-u/\tau}\right)\right)$$

while the contribution from the values within the range of the assay is the expectation of the term

$$\sum_{x\in\{x\}} \frac{e^{-x/\tau}}{\tau^4\left(e^{\frac{x}{\tau}}-1\right)^2}\left[-p_0\tau^2(x^2+2x\tau+2\tau^2) + e^{\frac{3x}{\tau}}\tau^2(2p_0\tau-1) + e^{\frac{x}{\tau}}\tau(-2x-\tau+2p_0x^2\tau+4p_0x\tau^2+6p_0\tau^3) - e^{2x/\tau}\left(-2\tau^2+6p_0\tau^4+2x\tau(p_0\tau-1)+x^2(1+p_0\tau^2)\right)\right]$$

For those elements of the Fisher Information for which we did not find a closed form solution of the expectation, we used the value at the maximum likelihood estimate (essentially assuming that the probability density of those values is peaked near our estimate). We then computed the Fisher Information and took the inverse to find the Cramér-Rao bound, which we used as our estimate of the covariance matrix. The plotted variances of the parameters correspond to the diagonal of this matrix.

The standard error of $p_{0,null}$, likewise, is

$$SE(p_{0,null}) = \left(\frac{p_{0,null}}{\sqrt{n}}\right)\left(\frac{1}{\sqrt{\left(1 - e^{-p_{0,null}u}\right) - \left(up_{0,null}e^{-up_{0,null}}\right)^{2}}}\right)$$

Because the maximum likelihood estimate is asymptotically normally distributed with variance given by the Cramér-Rao bound, for statistical comparisons of model fits we performed a Welch's t-test for unequal variances and unequal sample sizes:

**Supplementary Table 1: Statistical tests on model fits**

| **Figure, statistical test used** | **Null hypothesis** | **p-value (alpha = 0.05)**<br>**Bonferroni correction used to determine significance for multiple comparisons: (*0.05/n*), where *n* is number of comparisons made**<br>**Statistical significance indicated in red** |
|---|---|---|
| **Extended Data Figure 5** | | |
| B) Welch's t-test. 10 Minutes, low intensity = 1, medium = 2, high = 3, 15 minutes low intensity = 4, medium = 5, high =6 | No difference between the $\tau$ parameters across conditions | *<u>Corrected significance value</u>: 0.0033*<br><br>1-2: 0.101, 1-3: 0.535, 1-4: 0.0152, 1-5: 0.0059, 1-6: 0.057, 2-3: 0.186, 2-4: 0.019, 2-5: 0.00005, 2-6: 0.0292, 3-4: 0.017, 3-5: <0.000001, 3-6: 0.00619, 4-5: 0.036, 4-6: 0.2219, 5-6: 0.147 |
| B) Welch's t-test. 10 Minutes, low intensity = 1, medium = 2, high = 3, 15 minutes low intensity = 4, medium = 5, high =6 | No difference between the $p_0$ parameters across conditions | *<u>Corrected significance value</u>: 0.0033*<br><br>1-2: <0.000001, 1-3: <0.000001, 1- 4: 0.4376, 1-5: <0.000001, 1-6: 0.000001, 2-3: <0.000001, 2-4: <0.000001, 2-5: 0.0000158, 2-6: <0.000001, 3-4: <0.000001, 3-5: <0.000001, 3-6: 0.867, 4-5: <0.000001, 4-6: <0.000001, 5-6: <0.000001 |
| **Extended Data Figure 11** | | |

| | | |
|---|---|---|
| C) Welch's t-test | No difference between the $\tau$ parameters across conditions | 0.569 |
| D) Welch's t-test | No difference between the $p_0$ parameters across conditions | <0.000001 |
| G) Welch's t-test | No difference between the $\tau$ parameters across conditions | 0.00133 |
| G) Welch's t-test | No difference between the $p_0$ parameters across conditions | <0.000001 |

**STATISTICS (NON-MODELING DATA)**
Fisher's exact test, Mann Whitney U-test, and Kolmogorov-Smirnov test were performed using Prism 9. All tests are unpaired. All tests with multiple comparisons use post hoc Bonferroni corrections.

**Supplementary Table 2: Materials**

| REAGENT/RESOURCE | SOURCE | IDENTIFIER |
|---|---|---|
| **Antibodies** | | |
| Rabbit anti-GFP | Invitrogen | A-11122 |
| Chicken anti-GFP | Aves Labs | GFP-1010 |
| Mouse anti-GFP | Invitrogen | A11120 |
| Rabbit anti-DsRed | Clontech | 323496 |
| Mouse anti-nc82 | Developmental Studies Hybridoma Bank | N/A |
| Donkey anti-chicken 488 | Jackson ImmunoResearch | 703-545-155 |
| Donkey anti-rabbit 488 | Invitrogen | A11008 |
| Donkey anti-mouse 488 | Invitrogen | A21202 |
| Donkey anti-rabbit 555 | Invitrogen | A-31572 |
| Donkey anti-mouse Cy3 | Jackson ImmunoResearch | 715-166-150 |
| Donkey anti-rabbit 647 | Jackson ImmunoResearch | 711-605-152 |
| Donkey anti-mouse 647 | Jackson ImmunoResearch | 711-605-151 |

| **Chemicals, Peptides, Recombinant Proteins** | | |
|---|---|---|
| All-trans-retinal | Sigma Aldrich | R2500 |
| Dopamine | Alfa Aesar | A11136-06 |
| Sodium chloride | Fisher Chemical | S271-500 |
| Potassium chloride | Alfa Aesar | A11662.0B |
| TES | Thermo Fisher Scientific | B21819.18 |
| Trehyalose, Dihydrate | EMD Millipore Corp. | 625625-50GM |
| D-(+)-Glucose | Sigma Aldrich | G8270-1KG |
| Sodium bicarbonate | Acros Organics | 447102500 |
| Sodium phosphate | Research Products International | S23120-500.0 |
| Magnesium Chloride | Fisher Bioreagents | BP214-500 |
| Calcium Chloride | Fisher Chemical | C79-500 |
| **Experimental Models: Organisms/Strains** | | |
| NP2719 Gal4 | *Drosophila* Genome Resource Center | DGRC 113024 |
| NP5270 Gal4 | *Drosophila* Genome Resource Center | DGRC 113657 |
| R45G01 LexA | Bloomington Stock Center | BSC 54866 |
| TH LexA | Ron Davis lab | N/A |
| SS59854 AD, Dsx DBD | Barry Dickson lab | N/A |
| Repo-Gal80 | Tzumin Lee lab | N/A |
| UAS-Dicer2 | Bloomington Stock Center | BSC 24646 |
| UAS-CsChrimson-Tdtomato (attp2; attp40) | David Anderson lab | N/A |
| UAS-ChR2-XXM | Robert Kittel lab | N/A |
| UAS-GtACR1-eYFP | Adam Claridge/Chang lab | BSC 92983 |
| UAS-tntG | Sweeney et al., 1995 | N/A |
| LexAop2-CsChrimson-Tdtomato | Bloomington Stock Center | BSC 82183 |
| LexAop-TrpA1 | Scott Waddell lab | N/A |
| MCFO | Bloomington Stock Center | BSC 64086 |

| | | |
|---|---|---|
| UAS-myrGFP | Bloomington Stock Center | BSC 32197 |
| UAS-CD8-GFP | Lee and Luo, 1999 | N/A |
| UAS-green-Camuiα | Michael Crickmore lab | N/A |
| UAS-GCaMP6s (X; III) | David Anderson lab | N/A |
| RNAi and overexpression in **Figure 3A** | Bloomington Stock Center, Vienna Drosophila Resource Center, Michael Crickmore lab | Identifiers/stocks available upon request |
| 20x-UAS-CaMKII TT306/7DD | Michael Crickmore lab | N/A |
| 20x-UAS-CaMKII TT306/7AA | Michael Crickmore lab | N/A |
| 20x-UAS-CaMKII TT306/7SS | Michael Crickmore lab | N/A |
| 20x-UAS-CaMKII T287D | Michael Crickmore lab | N/A |
| CaMKII RNAi | Bloomington Stock Center | BSC 35330 |
| 20x-UAS-CaMKII T287D,K43M | Michael Crickmore lab | N/A |
| SS01593 ($AG_{Desc}$) | Gwyneth Card lab | N/A |
| Descending interneuron Gal4s from screen in **Extended Data Figure 4** | Gwyneth Card lab | Identifiers/stocks available upon request |
| Crz-LexA | Michael Crickmore lab | N/A |
| LexAop-Kir2.1 | | |
| LexAop-GFP | | |
| Gr28b, TrpA1 mutant | Marco Gallio lab | N/A |
| TH-LexA | Ron Davis lab | N/A |
| LexAop-GtACR1 | Michael Crickmore lab | N/A |
| Tshirt-Gal80 | Julie Simpson lab | N/A |

| | | |
|---|---|---|
| 10xUAS-GRAB-DA3m | Plasmid from Yulong Li, stock produced by SCT in Gaby Maimon lab | N/A |
| **Software and Algorithms** | | |
| Python FLIM code | This paper | https://github.com/CrickmoreRoguljaLabs/ |
| MATLAB calcium imaging code | This paper | https://github.com/CrickmoreRoguljaLabs/ |
| MATLAB copulation duration code | Thornquist et. al, 2020 | https://github.com/CrickmoreRoguljaLabs/CaMKIICode |
| Arduino solenoid control code | This paper | https://github.com/CrickmoreRoguljaLabs/ |
| Processing Java code | This paper | https://github.com/CrickmoreRoguljaLabs/ |
| SiffPy | Upcoming manuscript, SCT | https://github.com/MaimonLab/SiffPy |
| ScanImage-FLIM | Upcoming manuscript, SCT | To be made available at a later date |

**Supplementary Table 3: Genotypes and number of flies per experiment**

| **Figure** | **Label** | **Genotype (male, unless otherwise indicated)** | **Condition** | **N** |
|---|---|---|---|---|
| **Figure 1** | | | | |
| A) | - | - | - | - |
| B) | CDN>Chr-tdTomato | w-; NP2719-Gal4, repo-Gal80/+;UAS CsChrimson-tdTomato/+ | - | - |
| C) | CDN>GtACR1 | w-; NP2719-Gal4, repo-Gal80/+; UAS-GtACR1-eYFP/+ | No light | 19 |
| | | | Light | 17 |
| | CDN>GFP | w-; NP2719-Gal4, repo-Gal80/+; UAS-myrGFP/+ | No light | 19 |

| | | | | |
|---|---|---|---|---|
| | | | Light | 16 |
| | +/GtACR1 | w-; +/+; UAS-GtACR1-eYFP/+ | No light | 17 |
| | | | Light | 13 |
| D) | CDN>GtACR1 | w-; NP2719-Gal4/+; UAS-GtACR1-eYFP/+ | Light off at 0 min | 19 |
| | | | Off at 5 min | 11 |
| | | | Off at 20 min | 11 |
| | | | Off at 30 min | 18 |
| E) | CDN>Chr | w-; NP2719-Gal4/+;UAS CsChrimson-tdTomato/+ | No light | 89 |
| | 2 sec red light (no ret.) | | 1 min | 11 |
| | | | 5 min | 14 |
| | | | 10 min | 16 |
| | | | 15 min | 22 |
| | | | 20 min | 11 |
| | 2 sec red light | | 1 min | 35 |
| | | | 5 min | 34 |
| | | | 10 min | 20 |
| | | | 15 min | 37 |
| | | | 20 min | 14 |
| F) | CDN>Chr | w-; NP2719-Gal4, repo-Gal80/+;UAS CsChrimson-tdTomato/+ | | |
| | 10 μW/mm$^2$ green light | | 0.5 min | 17 |
| | | | 4 min | 10 |
| | | | 10 min | 31 |
| | | | 15 min | 29 |
| | 39°C | | 0.5 min | 10 |

| | | | | |
|---|---|---|---|---|
| | | | 4 min | 9 |
| | | | 10 min | 28 |
| | | | 15 min | 13 |
| | 8 μW/mm$^2$ green light | | 0.5 min | 9 |
| | | | 4 min | 11 |
| | | | 10 min | 30 |
| | | | 15 min | 33 |
| | 37°C | | 0.5 min | 7 |
| | | | 4 min | 12 |
| | | | 10 min | 23 |
| | | | 15 min | 25 |
| **Figure 2** | | | | |
| A) | CDN>Chr | w-; NP2719-Gal4/+;UAS CsChrimson-tdTomato/+ | | |
| | 1 sec | | 10 min | 51 |
| | | | 15 min | 60 |
| | 500 ms | | 10 min | 67 |
| | | | 15 min | 42 |
| B) | - | - | - | - |
| C) | CDN>Chr | w-; NP2719-Gal4/+;UAS CsChrimson-tdTomato/+ | | |
| | 10 min | | Single pulse | 53 |
| | | | 0.5 sec IPI | 70 |
| | | | 1 sec IPI | 72 |
| | | | 5 sec IPI | 79 |
| | | | 10 sec IPI | 88 |
| | | | 20 sec IPI | 67 |
| | | | 40 sec IPI | 69 |

| | | | | |
|---|---|---|---|---|
| | 15 min | | Single pulse | 56 |
| | | | 0.5 sec IPI | 65 |
| | | | 1 sec IPI | 64 |
| | | | 5 sec IPI | 75 |
| | | | 10 sec IPI | 71 |
| | | | 20 sec IPI | 67 |
| | | | 40 sec IPI | 81 |
| D) | CDN>GFP | w-; NP2719-Gal4, repo-Gal80/+; UAS-myrGFP/+ | 30 sec | 13 |
| | | | 4 min | 17 |
| | | | 10 min | 49 |
| | | | 15 min | 51 |
| E) | CDN>GFP | w-; NP2719-Gal4, repo-Gal80/+; UAS-myrGFP/+ | | 28 |
| | UAS tntG/+ | w-; UAS-tntG/+; +/+ | | 30 |
| | CDN>tntG | w-; NP2719-Gal4, repo-Gal80/UAS-tntG ; +/+ | | 25 |
| F) | CDN>GFP | w-; NP2719-Gal4, repo-Gal80/+; UAS-myrGFP/+ | | |
| | | 10 min | 5 sec IPI | 54 |
| | | | 10 sec IPI | 54 |
| | | | 20 sec IPI | 60 |
| | | 15 min | 5 sec IPI | 51 |
| | | | 10 sec IPI | 51 |
| | | | 20 sec IPI | 53 |
| G) | Groom>Chr, CDN>GFP | w-;R45G01-LexA,NP2719-Gal4/UAS-CD8-GFP;LexAop2-CsChrimson-tdTomato/+ | 1 min | 2 |
| | | | 5 min | 8 |

| | | | | |
|---|---|---|---|---|
| | | | 10 min | 25 |
| | | | 15 min | 15 |
| | Groom>Chr, CDN>Tnt | w-;R45G01-LexA,NP2719-Gal4/UAS-tntG;LexAop2-CsChrimson-tdTomato/+ | 1 min | 7 |
| | | | 5 min | 4 |
| | | | 10 min | 7 |
| | | | 15 min | 7 |
| H) | Groom>Chr | w-;R45G01-LexA/+;LexAop2-CsChrimson-tdTomato | | |
| | 10 min | | Single pulse | 57 |
| | | | 5 sec IPI | 36 |
| | 15 min | | Single pulse | 43 |
| | | | 5 sec IPI | 42 |
| I) | Groom>Chr, CDN>GtACR1 | w-;NP2719-Gal4, R45G01-LexA/UAS- GtACR1-eYFP;LexAop2-CsChrimson-tdTomato/+ | | |
| | Red only | | One pulse | 55 |
| | | | Two pulses | 44 |
| | Red + green | | One red pulse | 34 |
| | | | Two red pulses | 47 |
| | Groom>Chr, CDN>GFP | w-;NP2719-Gal4, R45G01-LexA/UAS- GFP;LexAop2-CsChrimson-tdTomato/+ | | |
| | Red only | | One pulse | 42 |
| | | | Two pulses | 41 |
| | Red + green | | One red pulse | 39 |

| | | | | |
|---|---|---|---|---|
| | | | Two red pulses | 44 |
| J) | - | - | - | - |
| K) | CDN>Chr | w-;NP2719-Gal4/+;UAS CsChrimson-tdTomato/+ | | |
| | 3.8 μW/mm$^2$ green light | | 10 min | 107 |
| | | | 15 min | 126 |
| | 9.2 μW/mm$^2$ green light | | 10 min | 116 |
| | | | 15 min | 110 |
| **Figure 3** | | | | |
| A) | CDN>Chr, genetic manipulation | Dicer-2; NP2719-Gal4, repo-Gal80/+;UAS CsChrimson-tdTomato/+ crossed into RNAi and UAS effector lines | | >2928 |
| B) | CDN> CaMKII RNAi | Dicer-2;NP2719-Gal4, repo-Gal80/NP2719-Gal4;UAS-CaMKII-RNAi/+ | 0.5 min | 26 |
| | | | 4 min | 23 |
| | | | 10 min | 20 |
| | CDN> GFP | Dicer-2;NP2719-Gal4, repo-Gal80/NP2719-Gal4;UAS-myrGFP/+ | 0.5 min | 14 |
| | | | 4 min | 17 |
| | | | 10 min | 17 |
| | RNAi/+ | w-;+/+;UAS-CaMKII-RNAi/+ | 0.5 min | 12 |
| | | | 4 min | 16 |
| | | | 10 min | 22 |
| C) | CDN>T287D | Dicer-2;NP2719-Gal4, repo-Gal80/+;UAS-20x-CaMKII-T287D/UAS-CD8-GFP | 0.5 min | 8 |

| | | | | |
|---|---|---|---|---|
| | | | 4 min | 11 |
| | | | 10 min | 9 |
| | | | 15 min | 11 |
| | | | 20 min | 12 |
| | CDN>GFP | Dicer-2;NP2719-Gal4, repo-Gal80/+;UAS-CD8-GFP/+ | 0.5 min | 9 |
| | | | 4 min | 7 |
| | | | 10 min | 20 |
| | | | 15 min | 8 |
| | T287D/+ | w-;+/+;UAS-20x-CaMKII-T287D/+ | 0.5 min | 7 |
| | | | 4 min | 7 |
| | | | 10 min | 16 |
| | | | 15 min | 18 |
| D) | CDN>Chr | | | |
| | - CaMKII RNAi | w-; NP2719-Gal4, repo-Gal80/+;UAS CsChrimson-tdTomato/+ | 10 min | 94 |
| | + CaMKII RNAi | Dicer-2; NP2719-Gal4, repo-Gal80/UAS CsChrimson-tdTomato; UAS-CaMKII-RNAi/+ | 10 min | 89 |
| | - CaMKII T287D | w-; NP2719-Gal4, repo-Gal80/+;UAS CsChrimson-tdTomato/+ | 15 min | 91 |
| | + CaMKII T287D | w-; NP2719-Gal4, repo-Gal80/UAS CsChrimson-tdTomato; UAS-20x-CaMKII-T287D/+ | 15 min | 87 |
| E) | CDN>Chr | | | |
| | - CaMKII RNAi | w-; NP2719-Gal4, repo-Gal80/+;UAS CsChrimson-tdTomato/+ | 60 sec green light | 27 |

| | | | | |
|---|---|---|---|---|
| | | | 0.5 sec red light | 51 |
| | + CaMKII RNAi | Dicer-2; NP2719-Gal4, repo-Gal80/UAS CsChrimson-tdTomato; UAS-CaMKII-RNAi/+ | 60 sec green light | 12 |
| | | | 0.5 sec red light | 50 |
| F) | CDN> CaMKII RNAi | Dicer-2;NP2719-Gal4, repo-Gal80/NP2719-Gal4;UAS-CaMKII-RNAi/+ | | |
| | | | Single pulse | 50 |
| | | | 10 sec IPI | 52 |
| | CDN> GFP | Dicer-2;NP2719-Gal4, repo-Gal80/NP2719-Gal4;UAS-myrGFP/+ | | |
| | | | Single pulse | 48 |
| | | | 10 sec IPI | 54 |
| | RNAi/+ | w-;+/+;UAS-CaMKII-RNAi/+ | | |
| | | | Single pulse | 50 |
| | | | 10 sec IPI | 50 |
| G) | CDN>Chr | | | |
| | - CaMKII RNAi | w-; NP2719-Gal4, repo-Gal80/+;UAS CsChrimson-tdTomato/+ | | |
| | 10 min | | 1 sec CL | 39 |
| | | | 2.5 sec CL | 51 |
| | | | 5 sec CL | 51 |
| | | | 10 sec CL | 53 |
| | + CaMKII RNAi | Dicer-2; NP2719-Gal4, repo-Gal80/UAS CsChrimson-tdTomato; UAS-CaMKII-RNAi/+ | | |
| | 10 min | | 1 sec CL | 45 |

| | | | | |
|---|---|---|---|---|
| | | | 2.5 sec CL | 51 |
| | | | 5 sec CL | 53 |
| | | | 10 sec CL | 54 |
| | - CaMKII T287D | w-; NP2719-Gal4, repo-Gal80/+;UAS CsChrimson-tdTomato/+ | | |
| | 15 min | | 1 sec CL | 50 |
| | | | 2.5 sec CL | 51 |
| | | | 5 sec CL | 55 |
| | | | 10 sec CL | 56 |
| | + CaMKII T287D | w-; NP2719-Gal4, repo-Gal80/UAS CsChrimson-tdTomato; UAS-20x-CaMKII-T287D/+ | | |
| | 15 min | | 1 sec CL | 52 |
| | | | 2.5 sec CL | 50 |
| | | | 5 sec CL | 56 |
| | | | 10 sec CL | 57 |
| H) | | | | |
| **Figure 4** | | | | |
| A) | CDN>Chr, TH>TrpA1 | w-; NP2719-Gal4/LexAop-TrpA1; TH-LexA/UAS-CsChrimson-tdTomato | No warmth | 108 |
| | | | Warmth | 119 |
| B) | CDN>GRAB-DA3m, TH>TrpA1 | w+; NP2719-Gal4, LexAop2-TrpA1/+; TH-LexA/UAS-GRAB-DA3m | | 15 |
| | | w+; NP2719-Gal4/+; TH-LexA/UAS-GRAB-DA3m | | 9 |
| C) | CDN>ChR2-XXM, green-Camuiα | w-;NP5270-Gal4/UAS-ChR2-XXM;UAS-green-Camuiα/+ | ~10 mW laser, pre/post 100 μM dopamine | 5 |

| | | | | |
|---|---|---|---|---|
| D) | CDN>green-Camuiα | w-;NP5270-Gal4, repo-Gal80/+;UAS-green-Camuiα/+ | pre/post 100 μM dopamine | 4 |
| E) | CDN>ChR2-XXM, GCaMP6s | w-;NP5270-Gal4/UAS-ChR2-XXM;UAS-GCaMP6s/+ | ~10 mW laser, saline | 5 |
| | | | ~10 mW laser, 100 μM dopamine | 5 |
| F) | CDN>ChR2-XXM, green-Camuiα | w-;NP5270-Gal4/UAS-ChR2-XXM;UAS-green-Camuiα/+ | 0 μM (saline) | 5 |
| | | | ~10 mW laser, 1 μM dopamine | 7 |
| | | | ~10 mW laser, 5 μM dopamine | 8 |
| | | | ~10 mW laser, 10 μM dopamine | 6 |
| **Extended Data Figure 1** | | | | |

| | | | | |
|---|---|---|---|---|
| A) | CDN>MCFO | pBPhsFlp2::PEST; NP2719-Gal4, repo-Gal80/+; pJFRC210-10XUAS-FRT>STOP>FRT-myr::smGFP-OLLAS, pJFRC201-10XUAS-FRT>STOP>FRT-myr::smGFP-HA, pJFRC240-10XUASFRT>STOP>FRT-myr::smGFP-V5-THS-10XUAS-FRT>STOP>FRT-myr::smGFP-FLAG/+ (MCFO-2 from Nern *et al.*[41], heat shocked twice for 15 minutes each at 37°C as an adult fly, separated by one day, and dissected at least three days later to allow for expression of the tags) | | |
| | CDN>SytGFP, Denmark | w-;NP2719-Gal4,repo-Gal80/+;UAS-SytGFP ,UAS-Denmark/+ | | |
| B) | CDN>GtACR1 | w-; NP2719-Gal4/+; UAS-GtACR1-eYFP/+ | No light | 8 |
| | | | Light on at 20 min | 18 |
| | | | Light | 9 |
| C) | CDN>Chr | w-; NP2719-Gal4/+;UAS CsChrimson-tdTomato/+ | No light | 89 |
| | | | No retinal | 11 |
| | | | Courtship | 7 |
| | | | Tonic | 30 |
| | | | Mating | 9 |
| D) | CDN>Chr | w-; NP2719-Gal4/+;UAS CsChrimson-tdTomato/+ | 10 min | 35 |
| | | | 15 min | 40 |

| | | | | |
|---|---|---|---|---|
| E) | CDN>GtACR1, Chr | w-; NP2719-Gal4/ UAS-GtACR1-eYFP;UAS CsChrimson-tdTomato/+ | No inhibition | 36 |
| | | | Inhibition of CDNs after sim | 38 |
| F) | | *Male genotype:* Dicer-2;NP2719-Gal4, repo-Gal80/+;UAS-CD8-GFP/+ | | |
| | Venus females | *Female genotype:* w;20xUAS-IVS-CsChrimson.mVenus/cyo;+/+ | 30 sec | 17 |
| | | | 4 min | 15 |
| | | | 10 min | 22 |
| | | | 15 min | 16 |
| | Heat insensitive females | *Female genotype:* w;Gr28bExc8/Gr28bExc8;TrpA1$^{1}$/ TrpA1$^{1}$ | 30 sec | 11 |
| | | | 4 min | 14 |
| | | | 10 min | 19 |
| | | | 15 min | 15 |
| **Extended Data Figure 3** | | | | |
| A) *see* ***Figure 2D*** | - | - | - | - |
| B) | CDN>GFP | w-; NP2719-Gal4, repo-Gal80/+; UAS-myrGFP/+ | 10 min | 50 |
| | | | 15 min | 49 |
| **Extended Data Figure 4** | | | | |
| A) | SS01593 ($AG_{Desc}$) | w-;R27E07-p65.AD/+;R20F03-Gal4.DBD/UAS-CsChrimson-tdTomato | | |
| | Unlabelled screen lines | ? | | |

| | | | | |
|---|---|---|---|---|
| B) | $AG_{Desc}$>Chrimson-tdTomato | w-;R27E07-p65.AD/+;R20F03-Gal4.DBD/UAS-CsChrimson-tdTomato | - | - |
| C) | $AG_{Desc}$>Chr | w-;R27E07-p65.AD/+;R20F03-Gal4.DBD/UAS-CsChrimson-tdTomato | 700ms at 10 min | 50 |
| | | | 300ms at 15 min | 54 |
| D) | $AG_{Desc}$>Chr | w-;R27E07-p65.AD/+;R20F03-Gal4.DBD/UAS-CsChrimson-tdTomato | | |
| | 10 min | | 1 sec IPI | 53 |
| | | | 5 sec IPI | 47 |
| | | | 10 sec IPI | 51 |
| | | | 20 sec IPI | 55 |
| | 15 min | | 1 sec IPI | 33 |
| | | | 5 sec IPI | 37 |
| | | | 10 sec IPI | 61 |
| | | | 20 sec IPI | 47 |
| **Extended Data Figure 5** | | | | |
| A) | - | - | - | - |
| B-D) | CDN>Chr | w-; NP2719-Gal4/+;UAS CsChrimson-tdTomato/+ | | |
| | 10 min | | Low intensity | 107 |
| | | | Medium | 116 |
| | | | High | 87 |
| | 15 min | | Low intensity | 126 |
| | | | Medium | 110 |
| | | | High | 85 |

| | | | | |
|---|---|---|---|---|
| **Extended Data Figure 7** | | | | |
| A) | CDN>CaMKII RNAi | Dicer-2;NP2719-Gal4, repo-Gal80/NP2719-Gal4;UAS-CaMKII-RNAi/+ | | 11 |
| | CDN>GFP | Dicer-2;NP2719-Gal4, repo-Gal80/NP2719-Gal4;UAS-myrGFP/+ | | 11 |
| | RNAi/+ | w-;+/+;UAS-CaMKII-RNAi/+ | | 13 |
| B) | CDN>GtACR1, CaMKII RNAi | Dicer-2;NP2719-Gal4, repo-Gal80/+;UAS-CaMKII-RNAi/ UAS-GtACR1-eYFP | No light | 10 |
| | | | Light | 11 |
| **Extended Data Figure 8** | | | | |
| A) | CDN>Chr, GCaMP6s | | | |
| | -T287D | UAS-GCaMP6s; NP5270-Gal4/UAS-CsChrimson-tdTomato;6B/+ | | 6 |
| | +T287D | UAS-GCaMP6s; NP5270-Gal4/UAS-CsChrimson-tdTomato;UAS-20x-CaMKII-T287D/+ | | 5 |
| B) | CDN>T287D | Dicer-2;NP2719-Gal4, repo-Gal80/+;UAS-20x-CaMKII-T287D/UAS-CD8-GFP | | 9 |
| | CDN>T287D,K43M | Dicer-2;NP2719-Gal4, repo-Gal80/if;UAS-20x-CaMKII-K43M, T287D/UAS-CD8-GFP | | 11 |
| C) | CDN>T287D | Dicer-2;NP2719-Gal4, repo-Gal80/+;UAS-20x-CaMKII-T287D/UAS-CD8-GFP | | 16 |
| | CDN>GFP | Dicer-2;NP2719-Gal4, repo-Gal80/+; UAS-CD8-GFP/+ | | 10 |

| | | | | |
|---|---|---|---|---|
| | T287D/+ | w-;+/+;UAS-20x-CaMKII-T287D/+ | | 11 |
| D) | CDN>T287D | Dicer-2;NP2719-Gal4, repo-Gal80/+;UAS-20x-CaMKII-T287D/UAS-CD8-GFP | | 28 |
| | CDN>GFP | Dicer-2;NP2719-Gal4, repo-Gal80/+; UAS-CD8-GFP/+ | | 9 |
| | T287D/+ | w-;+/+;UAS-20x-CaMKII-T287D/+ | | 11 |
| E) | CDN>GFP | Dicer-2;NP2719-Gal4, repo-Gal80/+; UAS-CD8-GFP/+ | | 18 |
| | CDN>WT CaMKII | Dicer-2;NP2719-Gal4, repo-Gal80/+;UAS-20x-CaMKII-WT/UAS-CD8-GFP | | 20 |
| F) | CDN>T287D, tshG80 | w-; NP2719-Gal4, repo-Gal80/tshirt-Gal80; UAS-20x-CaMKII-T287D/+ | | 18 |
| | CDN>GFP | w-; NP2719-Gal4, repo-Gal80/+; UAS-myrGFP/+ | | 17 |
| | CDN>T287D | w-; NP2719-Gal4, repo-Gal80/+; UAS-20x-CaMKII-T287D/+ | | 14 |
| G) | T287D/+ | w; +/+ ; UAS-20x-CaMKII-T287D/+ | | 27 |
| | CDN>GFP | w-; NP2719-Gal4, repo-Gal80/+; UAS-myrGFP/+ | | 37 |
| | CDN>T287D | w-; NP2719-Gal4, repo-Gal80/+; UAS-20x-CaMKII-T287D/+ | | 23 |
| **Extended Data Figure 9** | | | | |
| A) | CDN>Chr | w-; NP2719-Gal4, repo-Gal80/+;UAS CsChrimson-tdTomato/+ | Single pulse | 47 |

| | | | | |
|---|---|---|---|---|
| | CDN>Chr, CaMKII RNAi | Dicer-2; NP2719-Gal4, repo-Gal80/UAS CsChrimson-tdTomato; UAS-CaMKII-RNAi/+ | Single pulse | 49 |
| | | | 1 sec IPI | 28 |
| | | | 5 sec IPI | 51 |
| | | | 10 sec IPI | 59 |
| | | | 20 sec IPI | 50 |
| B) | CDN>Chr | w-; NP2719-Gal4, repo-Gal80/+;UAS CsChrimson-tdTomato/+ | Single pulse | 42 |
| | CDN>Chr, T287D | w-; NP2719-Gal4, repo-Gal80/UAS CsChrimson-tdTomato; UAS-20x-CaMKII-T287D/+ | Single pulse | 49 |
| | | | 1 sec IPI | 83 |
| | | | 5 sec IPI | 52 |
| | | | 10 sec IPI | 50 |
| | | | 20 sec IPI | 28 |
| C) | CDN>Chr | | | |
| | - CaMKII RNAi | w-; NP2719-Gal4, repo-Gal80/+;UAS CsChrimson-tdTomato/+ | 10 min | 155 |
| | + CaMKII RNAi | Dicer-2; NP2719-Gal4, repo-Gal80/UAS CsChrimson-tdTomato; UAS-CaMKII-RNAi/+ | 10 min | 158 |
| | - T287D | w-; NP2719-Gal4, repo-Gal80/+;UAS CsChrimson-tdTomato/+ | 15 min | 162 |
| | + T287D | w-; NP2719-Gal4, repo-Gal80/UAS CsChrimson-tdTomato; UAS-20x-CaMKII-T287D/+ | 15 min | 163 |
| **Extended Data Figure 10** | | | | |

| | | | | |
|---|---|---|---|---|
| A) | CDN>ChR2-XXM, green-Camuiα | w-;NP5270-Gal4/UAS-ChR2-XXM;UAS-green-Camuiα/+ | 10 sec blue light | 7 |
| | CDN>ChR2-XXM, GCaMP6s | w-;NP5270-Gal4/UAS-ChR2-XXM;UAS-GCaMP6s/+ | 10 sec blue light | 5 |
| B) | CDN>ChR2-XXM, green-Camuiα | w-;NP5270-Gal4/UAS-ChR2-XXM;UAS-green-Camuiα/+ | 10 sec blue light, pre/post 100 µM dopamine | 5 |
| C | pC1-split>ChR2-XXM, green-Camuiα | w-;*SS59854*-AD/UAS-ChR2-XXM;UAS-green-Camuiα/*Dsx*-DBD | ~10 mW laser, pre/post 100 µM dopamine | 4 |
| | pC1-split>ChR2-XXM, GCaMP6s | w-;*SS59854*-AD/UAS-ChR2-XXM;UAS-GCaMP6s/*Dsx*-DBD | ~10 mW laser, pre/post 100 µM dopamine | 3 |
| D) | CDN>ChR2-XXM, green-Camuiα | w-;NP5270-Gal4/UAS-ChR2-XXM;UAS-green-Camuiα/+ | | 4 |
| E) | CDN>ChR2-XXM, GCaMP6s | w-;NP5270-Gal4/UAS-ChR2-XXM;UAS-GCaMP6s/+ | ~10 mW laser, saline | 5 |
| | | | ~10 mW laser, 100 µM dopamine | 5 |
| F) | CDN>ChR2-XXM, GCaMP6s | w-;NP5270-Gal4/UAS-ChR2-XXM;UAS-GCaMP6s/+ | Saline | 4 |
| | | | 100 µM dopamine | 4 |
| G) | CDN>ChR2-XXM, green-Camuiα | w-;NP5270-Gal4/UAS-ChR2-XXM;UAS-green-Camuiα/+ | | 3 |
| **Extended Data Figure 11** | | | | |
| A) | Left and middle panels | w-; NP2719-Gal4, Repo-Gal80/LexAop-GFP; Crz-LexA/UAS-myr-tdTomato | | |

| | | | | |
|---|---|---|---|---|
| | Right panel | w-; NP2719-Gal4, Repo-Gal80/LexAop-SytGFP::HA; Crz-LexA/UAS-myr-tdTomato | | |
| B) | CDN>Chr, Crz>Kir | w-; NP2719-Gal4/LexAop-Kir2.1; Crz-LexA/UAS-CsChrimson-tdTomato | | 29 |
| C,D) | DIN>Chr, Crz>Kir2.1 | w-; NP2719-Gal4/LexAop-Kir2.1; Crz-LexA/UAS-CsChrimson-tdTomato | | 90 |
| | DIN>Chr, Crz>GFP | w-; NP2719-Gal4/LexAop-GFP; Crz-LexA/UAS-CsChrimson-tdTomato | | 91 |
| E) | | w-; NP2719-Gal4, Repo-Gal80/LexAop-myr-GFP; TH-LexA/UAS-CsChrimson-tdTomato | | |
| F) | TH>ACR1 | w-;+/+;TH-Gal4/UAS-GtACR1-eYFP | No light | 10 |
| | | | Light | 11 |
| | No retinal | | No light | 10 |
| | | | Light | 9 |
| G) | CDN>Chr | w-; NP2719-Gal4/+; UAS-CsChrimson-tdTomato | Warmth | 88 |
| | | | No warmth | 71 |

**Supplementary Table 4: Statistical tests**

| **Figure, statistical test used** | **Null hypothesis** | **p-value (alpha = 0.05)**<br>**Bonferroni correction used to determine significance for multiple comparisons: (*0.05/n*), where *n* is number of comparisons made**<br>**Statistical significance indicated in red** |
|---|---|---|
| **Figure 1** | | |
| C) Fisher's exact test, light vs. no light | No difference between termination probabilities | CDN>GtACR1: <0.0001, CDN>GFP: >0.9999, +/GtACR1: >0.9999 |
| D) Mann-Whitney U-test, groups numbered as 1=off at 0, 2=off at 5, 3=off at 20, 4=off at 30, 5=off at 200 | No difference between copulation durations | *Corrected significance value: 0.005*<br><br>1-2: 0.4449, 1-3: 0.7108, 1-4: <0.0001, 1-5: <0.0001, 2-3: 0.8044, 2-4: <0.0001, 2-5: <0.0001, 3-4: <0.0001, 3-5: <0.0001, 4-5: <0.0001 |
| E) Fisher's exact test, groups numbered as 1=CDN>Chr, 2=no light, 3=no retinal | No difference between termination probabilities within a timepoint | *Corrected significance value: 0.017*<br><br>*Time: 1 minute:* 1-2: <0.0001, 1-3: <0.0001, 2-3: >0.9999<br>*Time: 5 minutes:* 1-2: <0.0001, 1- 3: <0.0001, 2-3: >0.9999<br>*Time: 10 minutes:* 1-2: <0.0001, 1- 3: <0.0001, 2-3: >0.9999<br>*Time: 15 minutes:* 1-2: <0.0001, 1- 3: <0.0001, 2-3: >0.9999<br>*Time: 20 minutes:* 1-2: <0.0001, 1- 3: <0.0001, 2-3: >0.9999 |

| | | |
|---|---|---|
| F) Fisher's exact test, green light vs. heat | No difference between termination probabilities within a timepoint | ~10 $\mu W/mm^2$ green light vs. 39°C<br>*Time: 30sec:* >0.9999<br>*Time: 4min:* >0.9999<br>*Time: 10min:* 0.2893<br>*Time: 15min:* 0.1344<br><br>~8 $\mu W/mm^2$ green light vs. 37°C<br>*Time: 30 sec:* >0.9999<br>*Time: 4min:* >0.9999<br>*Time: 10min:* 0.6851<br>*Time: 15min:* 0.5964 |
| **Figure 2** | | |
| A) Fisher's exact test, 10 min vs. 15 min | No difference between termination probabilities | *500 ms pulse:* 0.7922<br>*1 sec pulse:* 0.7015 |
| C) Fisher's exact test, 10 min vs. 15 min | No difference between termination probabilities within an inter-pulse interval | *0.5sec IPI:* 0.3440<br>*1sec IPI:* 0.6545<br>*5sec IPI:* 0.0251<br>*10sec IPI:* 0.0988<br>*20sec IPI:* 0.3851<br>*40sec IPI:* 0.8711 |
| D) Fisher's exact test, groups numbered as 1= 30sec, 2= 4min, 3= 10min, 4=15min | No difference between termination probabilities | *Corrected significance value: 0.0083*<br><br>*1-2:* >0.9999, *1-3:* 0.1858, *1-4:* 0.0003, *2-3:* 0.1011, *2-4:* <0.0001, *3-4:* <0.0001 |
| E) Fisher's exact test, groups numbered as 1= CDN>GFP, 2= UAS tntG/+, 3= CDN>UAS tntG | No difference between termination probabilities | *Corrected significance value: 0.017*<br><br>1-2: 0.1844, 1-3: <0.0001, 2-3: <0.0001 |
| F) Fisher's exact test, 10 vs. 15 min | No difference between termination probabilities | *5sec IPI:* >0.9999<br>*10sec IPI:* 0.0237<br>*20sec IPI:* >0.9999 |

| | | |
|---|---|---|
| G) Fisher's exact test, between genotypes | No difference between termination probabilities within a timepoint | *Time*: *1 minute*: >0.9999,<br>*Time: 5 minutes*: >0.9999,<br>*Time: 10 minutes*: 0.0252,<br>*Time: 15 minutes:* 0.0002 |
| H) Fisher's exact test, 10 vs. 15 min | No difference between termination probabilities | *Single pulse:* 0.8151<br>*Two pulses, 5sec IPI:* 0.0013 |
| I) Fisher's exact test, red only vs. + green | No difference between termination probabilities | CDN>GtACR1<br>*One red pulse:* 0.0035<br>*Two red pulses:* 0.0119<br>CDN>GFP<br>*One red pulse:* >0.9999<br>*Two red pulses:* >0.9999 |
| J) Kolmogorov-Smirnov test, 10 vs. 15 min | No difference between distributions of termination times | 9.2 $\mu W/mm^2$ green light:<br><0.0001 |
| **Figure 3** | | |
| B) Fisher's exact test, groups numbered as 1= CDN>CaMKII RNAi, 2= CDN>GFP, 3= RNAi/+ | No difference between termination probabilities within a timepoint | *Corrected significance value: 0.017*<br><br>*Time: 30sec:* 1-2: 0.1426, 1-3: 0.1583, 2-3: >0.9999<br><br>*Time: 4min:* 1-2: 0.0009, 1-3: 0.0003, 2-3: >0.9999<br><br>*Time: 10min:* 1-2: 0.1590, 1-3: 0.0221, 2-3: 0.4940 |
| C) Fisher's exact test, groups numbered as 1= CDN>T287D, 2= CDN>GFP, 3= T287D/+ | No difference between termination probabilities within a timepoint | *Corrected significance value: 0.017*<br><br>*Time: 30sec:* 1-2: >0.9999, 1-3: >0.9999, 2-3: >0.9999<br><br>*Time: 4min:* 1-2: >0.9999, 1-3: >0.9999, 2-3: >0.9999<br><br>*Time: 10min:* 1-2: 0.0794, 1-3: 0.0732, 2-3: >0.9999<br><br>*Time: 15min:* 1-2: <0.0001, 1-3: <0.0001, 2-3: 0.5292 |

| | | |
|---|---|---|
| D) Kolmogorov-Smirnov test, CaMKII vs. no CaMKII manipulation | No difference between distributions of termination times | <u>10 min:</u><br><0.0001<br><u>15 min:</u><br><0.0001 |
| E) Fisher's exact test, 0.5 sec red light vs. 60 sec green light | No difference between termination probabilities within a genotype | *+CaMKII RNAi:* 0.0096<br>*-RNAi:* >0.9999 |
| F) Fisher's exact test, groups numbered as 1= CDN>CaMKII RNAi, 2= CDN>GFP, 3= RNAi/+ | No difference between termination probabilities within condition | *<u>Corrected significance value:</u>*<br>*0.017*<br><br>*Single pulse:*<br>1-2: 0.5150, 1-3: 0.8305, 2-3: 0.8248<br><br>*Paired pulse, 10sec IPI:*<br>1-2: 0.0049, 1-3: 0.0566, 2-3: 0.4376 |
| G) Kolmogorov-Smirnov test, CaMKII vs. no CaMKII manipulation | No difference between distributions of responses within cycle lengths | <u>10 min</u><br>*1s:* 0.0233<br>*2.5s:* 0.0059<br>*5s:* 0.0002<br>*10s:* 0.0006<br><u>15 min</u><br>*1s:* 0.8079<br>*2.5s:* 0.9934<br>*5s:* 0.0033<br>*10s:* 0.0006 |
| **Extended Data Figure 1** | | |
| B) Mann-Whitney U-test, no light vs. light on at 20 min | No difference between copulation durations | 0.0072 |
| C) Mann-Whitney U-test, groups numbered as 1=no light, 2=no retinal, 3=courtship, 4=tonic, 5=mating | No difference between copulation durations | *<u>Corrected significance value</u>:*<br>*0.005*<br><br>1-2: 0.2578, 1-3: 0.9529, 1-4: <0.0001, 1-5: <0.0001, 2-3: 0.4960, 2-4: <0.0001, 2-5: <0.0001, 3-4: <0.0001, 3-5: 0.0002, 4-5: <0.0001 |

| | | |
|---|---|---|
| D) Kolmogorov-Smirnov test | No difference between distributions of termination times | <0.0001 |
| E) Kolmogorov-Smirnov test | No difference between distributions of termination times | <0.0001 |
| F) Fisher's exact test, Venus vs. Heat insensitive females | No difference between termination probabilities within a timepoint | *Time: 30sec:* >0.9999<br>*Time: 4min:* >0.9999<br>*Time: 10min:* >0.9999<br>*Time: 15min:* >0.9999 |
| **Extended Data Figure 3** | | |
| B) Fisher's exact test, 10 vs. 15 min | No difference between termination probabilities | >0.9999 |
| **Extended Data Figure 4** | | |
| C) Fisher's exact test, 10 vs. 15 min | No difference between termination probabilities | 0.6655 |
| D) Fisher's exact test, 10 vs. 15 min | No difference between termination probabilities | *1sec IPI:* 0.1521<br>*5sec IPI:* 0.0071<br>*10sec IPI:* 0.0132<br>*20sec IPI:* 0.1562 |
| **Extended Data Figure 7** | | |
| A) Mann-Whitney U-test, groups numbered as 1=CDN>CaMKII RNAi, 2=CDN>GFP, 3=RNAi/+ | No difference between copulation durations | *Corrected significance value: 0.017*<br><br>1-2: 0.5190, 1-3: 0.2518, 2-3: 0.5215 |
| B) Fisher's exact test, light vs. no light | No difference between termination probabilities | 0.0039 |
| **Extended Data Figure 8** | | |
| A) Mann-Whitney U-test, between genotypes | No difference between values | 2 sec<br>*Peak:* 0.7922<br>*Residual:* 0.1255<br><br>5 sec<br>*Peak:* 0.5368<br>*Residual:* 0.1255 |

| | | |
|---|---|---|
| B) Fisher's exact test, between genotypes | No difference between termination probabilities | <0.0001 |
| C) Fisher's exact test, groups numbered as 1= CDN>T287D, 2= CDN>GFP, 3= T287D/+ | No difference between fertility probabilities | *Corrected significance value: 0.017*<br><br>1-2: 0.5385, 1-3: >0.9999, 2-3: 0.5865 |
| D) Mann-Whitney U-test, groups numbered as 1= CDN>T287D, 2= CDN>GFP, 3= T287D/+ | No difference between copulation durations | *Corrected significance value: 0.017*<br><br>1-2: <0.0001, 1-3: <0.0001, 2-3: 0.5516 |
| E) Fisher's exact test, between genotypes | No difference between termination probabilities | 0.6062 |
| F) Fisher's exact test, groups numbered as 1= CDN>T287D,tshG80; 2= CDN>GFP; 3= CDN>T287D | No difference between termination probabilities | *Corrected significance value: 0.017*<br><br>1-2: 0.4018, 1-3: <0.0001, 2-3: <0.0001 |
| G) Fisher's exact test, groups numbered as 1= T287D/+, 2= CDN>GFP, 3= CDN>T287D | No difference between termination probabilities | *Corrected significance value: 0.017*<br><br>1-2: 0.0035, 1-3: <0.0001, 2-3: <0.0001 |
| **Extended Data Figure 9** | | |
| A) Fisher's exact test, between genotypes (single pulse) | No difference between termination probabilities | 0.0229 |
| B) Fisher's exact test, between genotypes (single pulse) | No difference between termination probabilities | <0.0001 |
| C) Fisher's exact test, between genotypes | No difference between termination probabilities | 10 min: 0.0365<br>15 min: 0.0146 |
| **Extended Data Figure 10** | | |
| E) Mann-Whitney U-test, dopamine vs. saline | No differences between peaks | 0.3095 |
| F) Mann-Whitney U-test, dopamine vs. saline | No differences between values | *Peak*: 0.6857<br>*Residual:* 0.0286 |
| **Extended Data Figure 11** | | |

| | | |
|---|---|---|
| F) Mann-Whitney U test, conditions numbered from left to right | No difference between the distribution of copulation durations | 1-2: 0.357, 1-3:0.9681, 1-4: 0.778, 2-3: 0.289, 2-4: 0.497, 3-4: 0.653 |
